# Resource limitation rewires chromosome instability and ploidy evolution across in vitro and in vivo cancer models

**DOI:** 10.64898/2026.09.16.752131

**Authors:** Tao Li, Richard Beck, Vural Tagal, Xiaoqing Yu, Noemi Andor

## Abstract

Whole-genome doubling (WGD) and elevated ploidy are pervasive features of cancer that shape chromosomal instability (CIN), therapeutic response, and metastatic fitness. Yet ploidy is strikingly context dependent: many tumors remain near diploid in vivo despite the frequent emergence of highly polyploid states in vitro. Here, we estimate ploidy dependent chromosome missegregation tolerance using a mathematical model calibrated to growth and chromosome number data from matched near diploid and near tetraploid breast cancer cultures and xenografts. The model architecture reflects a tug of war between two opposing selective forces acting on ploidy–resource limitation that caps high ploidy by imposing energetic and biosynthetic costs, and CIN that can favor higher ploidy by buffering the fitness impact of chromosome gains and losses. The fitted model reproduced chromosome losses in 4N cultures and WGD followed by chromosome losses in 2N cultures. In vivo, the strongest determinants of ploidy shifted from stress associated death at low oxygen to baseline missegregation at higher oxygen. Joint calibration to both in vivo and in vitro contexts assigned tumors lower proliferation, an approximately tenfold higher stress-associated death scale, and an 11-16 fold larger maximal stress induced missegregation increment than cultures. Simulating populations across combinations of constant oxygen levels and missegregation settings showed that ploidy increased with oxygen under conditions supporting population growth. In both culture and tumors, mean ploidy above tetraploidy was associated with population decline. Together, the framework predicts when resource constraints favor chromosome loss and when missegregation tolerance permits high ploidy expansion, providing testable expectations for how CIN perturbations reshape ploidy evolution in different resource environments.

## Introduction

Whole-genome doubling (WGD) and aneuploidy are common features of cancer evolution, yet high-ploidy states are distributed unevenly across biological contexts^1–3^. Cultured cancer cell lines generally exhibit higher ploidy than primary tumors, raising the possibility that resource-rich culture conditions relax constraints on excess genomic material^4,5^. Oxygen provides a tractable first-order proxy for this resource landscape^4–6^. Across tissue sites, WGD prevalence correlates with peripheral oxygen partial pressure, consistent with the idea that well-oxygenated tissues are more permissive for WGD-positive cancers^5,6^. A cellular basis for this relationship is suggested by observations of increased mitochondrial content and membrane potential in tetraploid cancer-cell models^7^. Because mitochondrial respiration requires oxygen, these ploidy-associated mitochondrial changes provide an additional rationale for expecting oxygen availability to influence the relative fitness of high- and low-ploidy cells. Increased chromosome content carries biosynthetic demands associated with DNA replication, RNA production, and protein synthesis^4–6^. Aneuploidy can add further costs through gene-dosage imbalance and the proteostatic, replication, and other stress responses needed to maintain cellular function^8–10^. Resource limitation could therefore strengthen selection against some high-chromosome states. However, this expectation need not apply uniformly to every karyotype: experimentally introduced trisomies can impair proliferation under standard conditions yet confer a relative advantage under selected environmental stresses^11^.

At the same time, environmental stress can also increase genomic instability^4,5^. High mutation rates are usually costly because most new mutations are deleterious, but in adapting asexual populations mutator states can hitchhike with beneficial mutations during strong selection^12^. Stress-induced mutagenesis provides a way to reduce this cost by increasing mutation mainly in stressed or maladapted cells, thereby increasing adaptability while limiting mutational load in well-adapted cells^12,13^. Chromosome instability may represent a chromosome-scale analogue of this logic: a single segregation or cell-division error can alter many loci at once and generate large jumps through genotype and karyotype space^14,15^. Indeed, environmental stresses such as serum starvation, hypoxia, folate deficiency, and energy restriction have been linked to karyotypic instability, chromosome mis-segregation, polyploidization, and increased cellular heterogeneity. Chromosome missegregation can lead to arrest, senescence, or death, while genome doubling can reduce the relative dosage change associated with a segregation error^16–18^. Live-cell analyses of diploid and tetraploid colorectal clones showed that tetraploid clones tolerated segregation errors more readily without a detected increase in errors per chromosome^19^. Longitudinal karyotyping echoed these findings, showing that the fitness effects of chromosome alterations differ significantly by WGD status^20^. These observations identify a central tension: high ploidy can increase resource-stress while also helping cells retain viable descendants after stress-induced segregation errors^21^. Whether WGD-derived populations expand, persist, or remodel toward lower chromosome numbers should therefore depend on how the environment changes both resource costs and the production and survival of altered descendants.

Tagal et al. recently reported matched culture and tumor experiments in near-tetraploid and near-diploid SUM159 lineages that provide an opportunity to examine this balance^22^. In vivo, near-tetraploid cells formed tumors after a longer delay and exhibited substantial ploidy reduction toward near-diploid states, whereas tumors derived from near-diploid cells remained predominantly near-diploid. This reduction echoes an independent observation by Bloomfield et al.: cells recovered from xenografts initiated with the tetraploid CAL51-L3 breast cancer clone exhibited increased chromosome-number heterogeneity and a shift toward near-triploid karyotypes, although many cells in the other tested lines remained near-tetraploid^23^. In Tagal et al.’s oxygen-deprived cultures, near-tetraploid lineages likewise lost chromosomes, but near-diploid lineages followed a different trajectory: WGD intermediates emerged and expanded before undergoing subsequent chromosome loss^22^. This sequence is consistent with evolutionary models in which tetraploidy facilitates access to favorable near-triploid karyotypes by buffering the fitness costs of chromosome loss^24^. Together, these findings motivate a quantitative account of how environmental conditions influence both the generation of chromosome-number variation and the persistence of its descendants.

Here, to quantitatively interpret ploidy dynamics reported in Tagal et al., we develop a mathematical model of ploidy evolution under hypoxia. We integrate growth, chromosome-number, necrosis, and flow-cytometry observations from near-diploid and near-tetraploid SUM159 cultures and xenografts to ask why chromosome loss can accompany both the loss of an initial high-ploidy state and the expansion of a WGD-compatible component. We first test whether a common model of stress-induced CIN and ploidy-dependent survival can explain both in vitro and in vivo trajectories, then ask how resource costs and chromosome-error tolerance differ between tumors and cell lines. The selected fits point to stronger resource penalties in tumors but a larger ploidy-associated survival gain in culture, linking the observed contrast to different balances between chromosome-variant production and retention. Finally, fixed-oxygen and finite-time analyses examine how these differences shape transient trajectories and long-term ploidy composition.

## Results

### A model of ploidy evolution coupling stress-induced CIN with ploidy-dependent CIN buffering

To understand how resource stress reshapes ploidy from different starting states, we analyzed chromosome-number dynamics in matched SUM159 near-diploid (2N) and near-tetraploid (4N) in vitro cultures and orthotopic engraftments reported by Tagel et al^22^. The experimental design included control and oxygen-deprived culture lineages, together with repeated tumor-burden measurements and terminal chromosome sampling in vivo (Figure 1A). Under oxygen deprivation, the two starting-ploidy backgrounds exhibited distinct population-level trajectories: chromosome numbers decreased in the 4N lineage, whereas the 2N lineage showed an initial increase consistent with whole-genome doubling followed by chromosome loss (Figure 1B). In vivo, terminal chromosome-number distributions from four tumors per starting-ploidy cohort also showed substantial ploidy reduction in the 4N-derived tumors (Figure 1D).

**Figure 1:**
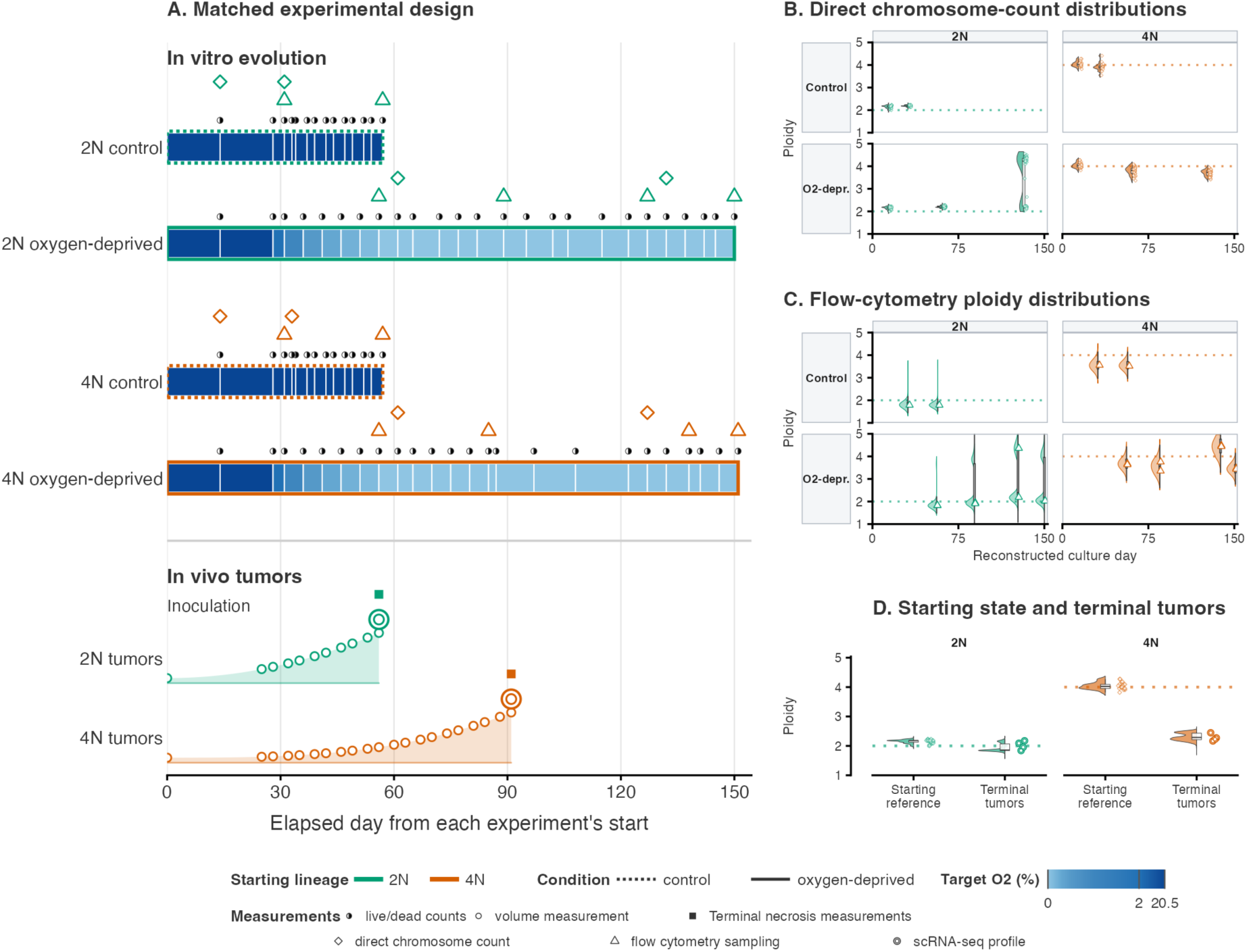
Matched culture and tumor measurements used to calibrate the ploidy-evolution model. Data from Tagal et al.^22^ comprise SUM159 near-diploid (2N) and near-tetraploid (4N) lineages studied during serial culture under control or oxygen-deprived conditions and after orthotopic engraftment. **(A)** Reconstructed culture-passage and tumor-sampling timelines, each retaining its own time origin. Cultures were maintained at 20.5% O_2_ or subjected to stepwise deprivation to 0% O_2_; tumor-growth wedges are schematic. In the 4N cohort, the exact day of terminal necrosis and scRNA-seq varied among mice; the corresponding 4N symbols mark the latest displayed sampling day, not a common sampling day. **(B)**Direct chromosome-count distributions in control and oxygen-deprived cultures, with autosomal counts divided by 22 to express ploidy. Shared starting measurements are repeated for alignment but not counted as additional samples. **(C)** G0/G1 DNA-content distributions on the reconstructed culture-day axis. Same-day profiles from parallel O1 and O2 deprivation lineages contribute equally; tails outside the displayed ploidy range of 1–5 are cropped without renormalization. **(D)** Direct chromosome counts from starting cultures (20 cells per cohort) and terminal RNA-derived chromosome estimates from 6,502 cells across eight tumors (four per cohort). Starting and terminal specimens are distinct samples. See Supplementary Table 1.

These observations motivated a model in which resource limitation can both oppose and promote high-ploidy states. We developed a deterministic model of ploidy evolution that tracks viable cells and dead biomass across autosomal chromosome-count states, coupling stress-induced CIN with ploidy-dependent CIN buffering and growth (Figure 2). The model contains 14 core biological response parameters present in both in-vivo and in-vitro settings (Table 1). Resource stress suppresses division and increases the death hazard, with both effects depending on chromosome number. The stress-associated death hazard, *µ*_eff_ (*N, O*_2_), also serves as the signal driving the per-chromosome-copy missegregation probability per division:

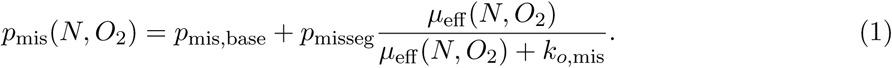

**Figure 2:**
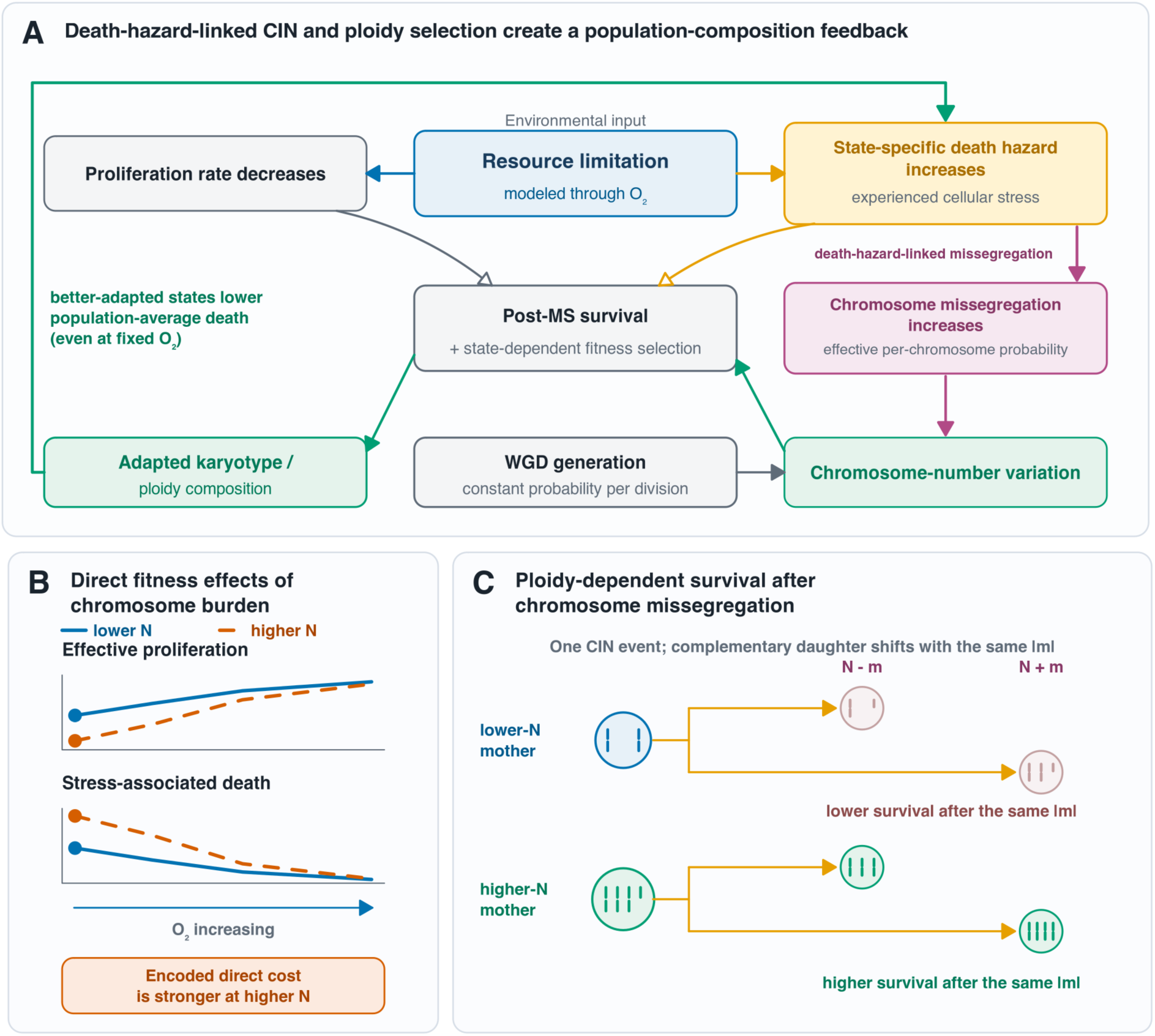
A model coupling stress-induced chromosome missegregation with ploidy-dependent survival and adaptive feedback. **(A)** Resource limitation suppresses proliferation and increases a chromosome-state-specific death hazard that also drives inducible chromosome missegregation. Survival filtering and differential growth change population composition; enrichment of better-adapted states can then reduce population-average death and chromosomal instability (CIN), even at unchanged oxygen. Whole-genome doubling (WGD) provides a separate chromosome-number transition with constant probability per division. Panels B–C illustrate relationships assumed by the model: **(B)** Schematic direct fitness effects: under oxygen deprivation, higher-chromosome states (orange) experience stronger proliferation suppression and stress-associated death than lower-chromosome states (blue). **(C)** Complementary daughters produced by missegregation contain *N ↑ m* and *N* + *m* chromosomes, where *N* is the maternal chromosome count and *m* the number of missegregated copies. Higher maternal ploidy increases survival for a given *m*; gain and loss daughters receive the same survival weighting. Faded daughters indicate lower survival.

**Table 1:**
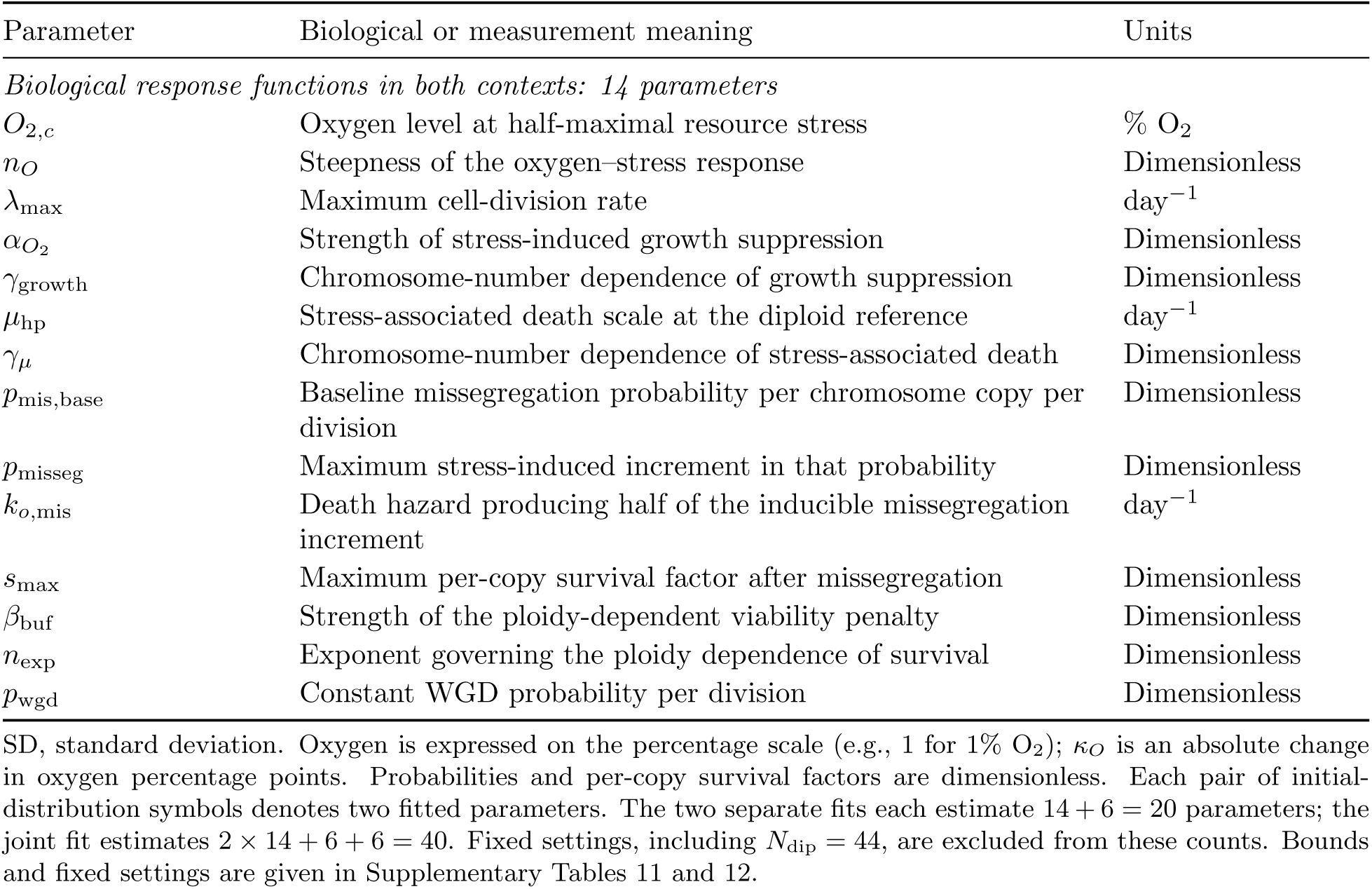
Parameters governing ploidy evolution and its calibration. The first 14 parameters describe biological response functions present in both contexts; their fitted values may differ under soft coupling. Additional parameters describe tumor oxygen supply, biomass handling, initial culture distributions, and measurement variability.

| Parameter | Biological or measurement meaning | Units |
| --- | --- | --- |
| *Biological response functions in both contexts: 14 parameters* |  |  |
| $O_{2,c}$ | Oxygen level at half-maximal resource stress | % O<sub>2</sub> |
| $n_O$ | Steepness of the oxygen–stress response | Dimensionless |
| $\lambda_{\text{max}}$ | Maximum cell-division rate | day<sup>−1</sup> |
| $\alpha_{O_2}$ | Strength of stress-induced growth suppression | Dimensionless |
| $\gamma_{\text{growth}}$ | Chromosome-number dependence of growth suppression | Dimensionless |
| $\mu_{\text{hp}}$ | Stress-associated death scale at the diploid reference | day<sup>−1</sup> |
| $\gamma_{\mu}$ | Chromosome-number dependence of stress-associated death | Dimensionless |
| $p_{\text{mis,base}}$ | Baseline missegregation probability per chromosome copy per division | Dimensionless |
| $p_{\text{misseg}}$ | Maximum stress-induced increment in that probability | Dimensionless |
| $k_{o,\text{mis}}$ | Death hazard producing half of the inducible missegregation increment | day<sup>−1</sup> |
| $s_{\text{max}}$ | Maximum per-copy survival factor after missegregation | Dimensionless |
| $\beta_{\text{buf}}$ | Strength of the ploidy-dependent viability penalty | Dimensionless |
| $n_{\text{exp}}$ | Exponent governing the ploidy dependence of survival | Dimensionless |
| $p_{\text{wgd}}$ | Constant WGD probability per division | Dimensionless |

SD, standard deviation. Oxygen is expressed on the percentage scale (e.g., 1 for 1% O<sub>2</sub>); $\kappa_O$ is an absolute change in oxygen percentage points. Probabilities and per-copy survival factors are dimensionless. Each pair of initial-distribution symbols denotes two fitted parameters. The two separate fits each estimate 14 + 6 = 20 parameters; the joint fit estimates 2 × 14 + 6 + 6 = 40. Fixed settings, including $N_{\text{dip}} = 44$ , are excluded from these counts. Bounds and fixed settings are given in Supplementary Tables 11 and 12.

Here, *N* is the mother’s autosomal chromosome count, *p*_mis,base_ is the baseline probability, *p*_misseg_ is the maximum stress-induced increment, and *k_o,_*_mis_ is the death hazard at which that increment is half-maximal. Because the death hazard depends on chromosome number, selection toward states experiencing less stress can reduce subsequent population-average CIN even at constant environmental oxygen.

Ordinary divisions generate complementary daughter states *N*–*m* and *N* +*m*, where *m* follows a binomial distribution with *N* trials and probability *p*_mis_(*N, O*_2_). For daughters within the modeled chromosome-count range, survival after a shift of Δ chromosomes is

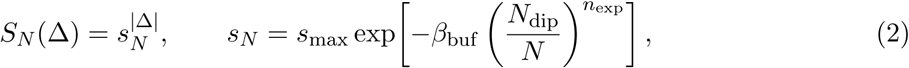

where *N*_dip_ = 44 is the diploid autosomal reference and *s_N_* is the per-copy survival factor. Thus, higher-ploidy mothers present more chromosome copies that can missegregate but confer greater survival for a given number of gains or losses. A separate WGD branch maps *N* to 2*N* with constant probability *p*_wgd_ per division. Ploidy evolution therefore reflects the interplay between resource-dependent proliferation and death, chromosome variation, and retention of altered descendants. The full population equations, oxygen supply–demand relation, passage rules, calibration, and evaluation are described in Supplementary Section 1.3; parameter bounds and fixed settings are listed in Supplementary Tables 11 and 12.

### Ploidy evolution model recapitulates direct ploidy reduction from 4N and genome doubling followed by reduction from 2N lineages

To test whether the same chromosome-generation and survival processes could explain the two culture trajectories (Figure 1A), we first fitted the model to passage-level growth rates, direct chromosome counts, and G0/G1 DNA-content profiles from three data streams: control and oxygen-deprived SUM159 2N and 4N cultures (Figure 1B). Populations were propagated through successive passages under their assigned oxygen conditions. We estimated 20 parameters by fitting the three data streams simultaneously, with equal weights for their averaged log-likelihoods, using 500 optimization runs combining differential evolution and local L-BFGS-B refinement (Supplementary Section 1.3.1, Eqs. (42)–(47)).

The lowest-objective fit, seed 144, reproduced the reduction in passage growth under oxygen deprivation and the broad remodeling of DNA-content profiles (Figure 3A–B). The 4N-derived population shifted progressively toward lower chromosome numbers. From the 2N starting state, the model generated a high-chromosome component compatible with WGD; this component expanded while its chromosome-number distribution shifted downward, alongside a persisting lower-chromosome mode (Figure 3C).

**Figure 3:**
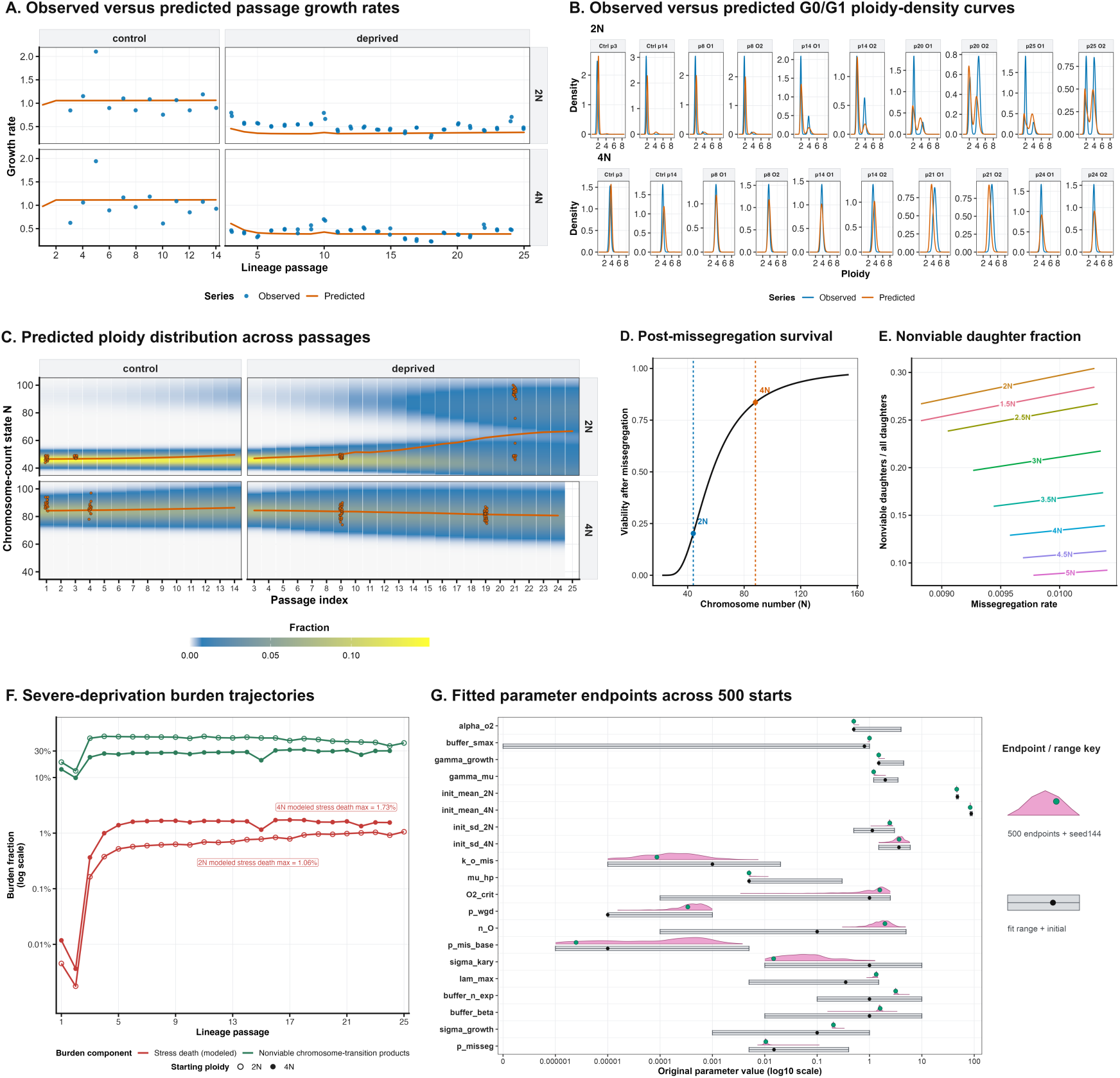
The model reproduces WGD expansion followed by ploidy reduction in the 2N and direct ploidy reduction in the 4N lineage. Panels A–F use the lowest-objective retained separate *in vitro* fit (seed 144). **(A)** Observed passage growth rates (blue points) and fitted predictions (orange), arranged by starting ploidy and control or deprivation condition. **(B)** Observed (blue) and fitted (orange) G0/G1 DNA-ploidy densities. Measurements from parallel O1 and O2 deprivation lineages are displayed separately with their corresponding predictions. **(C)** Predicted viable-cell fractions across chromosome states and passages. Orange curves show whole-population mean chromosome number; outlined orange points show direct single-cell counts at sampled passages. In the 2N lineage, subsequent chromosome loss occurs within the high-chromosome component compatible with whole-genome doubling. Intervening distributions are model predictions. **(D)** Fitted per-copy survival factor, *s_N_*, under the assumed nondecreasing dependence on maternal chromosome number. Dashed lines mark the 2N (*N* = 44, blue) and 4N (*N* = 88, orange) references. Survival after a shift of *m* chromosome copies is *s^m^*. **(E)** Expected nonviable misseg-regation products per division, divided by two, plotted against effective per-chromosome missegregation probability at the labeled maternal ploidies. This two-daughter normalization excludes out-of-grid losses. **(F)** Passage-end fractions of total modeled burden assigned to resource-stress death (red) or nonviable chromosome-transition products (green), including out- of-grid losses in the latter. Open and filled circles distinguish 2N and 4N lineages. The vertical axis is logarithmic. These retained compartments include clearance and are not cumulative death probabilities. **(G)** Numerical-search distributions for 20 fitted parameters across 500 starts. Pink half-violins show retained endpoint values; green points mark seed 144. Gray ranges and black points indicate fitting bounds and configured initial values. The log_10_ axis includes a dedicated position for zero bounds. Endpoint spread describes optimizer variation, not posterior uncertainty (Supplementary Tables 1 and 6).

The fitted survival function helps explain how high-ploidy expansion and downward remodeling can coexist. The per-copy post-missegregation survival factor increased from approximately 0.20 at *N* = 44 to 0.84 at *N* = 88 (Figure 3D–E). For a given chromosome-count change, daughters produced by high-ploidy mothers therefore have greater modeled survival, including daughters that have lost chromosomes. High-ploidy cells can thus persist while also supplying viable lower-chromosome descendants. As these descendants survive and proliferate, they can contribute to downward remodeling of the lineage without requiring direct resource-stress death to eliminate its high-ploidy ancestors.

Consistent with this interpretation, across the ten lowest-objective fits, mean chromosome number in the 4N lineage declined by 3.56–4.34 chromosomes while the maximum daily fraction of total burden assigned to resource-stress death remained at 1.69–1.77%. In seed 144, the maximum passage-end fractions were 1.06% for the 2N lineage and 1.73% for the 4N lineage; nonviable chromosome-transition products contributed the larger dead component (Figure 3F; Supplementary Table 6).

The model captured these broad trajectories but did not reproduce the full divergence between parallel deprived 2N lines. At the final sampled passage, 40% of counted O1 cells and 80% of counted O2 cells had *N ↑* 80, compared with fitted fractions of 32.7% and 32.4%, respectively (20 counted cells per observed sample; Supplementary Table 5). Different sequences of chromosome alterations can generate karyotypes with distinct fitness at the same ploidy, providing a source of replicate variation that a model tracking total chromosome number rather than individual karyotypes cannot resolve. Nevertheless, the same fitted model captured the central contrast: chromosome loss in the 4N lineage and high-ploidy expansion followed by reduction within the 2N lineage.

### The strongest ploidy association shifts from stress-associated death to baseline missegregation as oxygen increases

To ask how ploidy evolution differs in the tumor resource environment, we next fitted the model to a multimodal datasets consisting of three components: (i) longitudinal tumor-burden measurements, (ii) terminal single-cell chromosome-number estimates, and (iii) available histologic necrosis fractions from eight untreated SUM159 xenografts, four from each starting-ploidy cohort. We estimated 20 parameters using a maximum-a-posteriori objective combining the three data contributions with parameter-prior penalties, applying the same 500-run optimization strategy used for the culture fit (Supplementary Section 1.3.1, Eq. (40)). Tumor oxygen was inferred through the burden-dependent supply–demand model.

The lowest-objective fit, seed 25, captured the cohort-scale increase in tumor burden and the contrasting ploidy outcomes of the two starting populations (Figure 4A). The fitted 2N-derived population remained near diploidy, whereas the 4N-derived population shifted toward a state only slightly above diploid as inferred effective oxygen declined. Predicted terminal distributions captured the cohort-specific central ploidies but were broader than observed. Detailed growth, chromosome-distribution, and necrosis residuals are reported in Supplementary Table 2.

**Figure 4:**
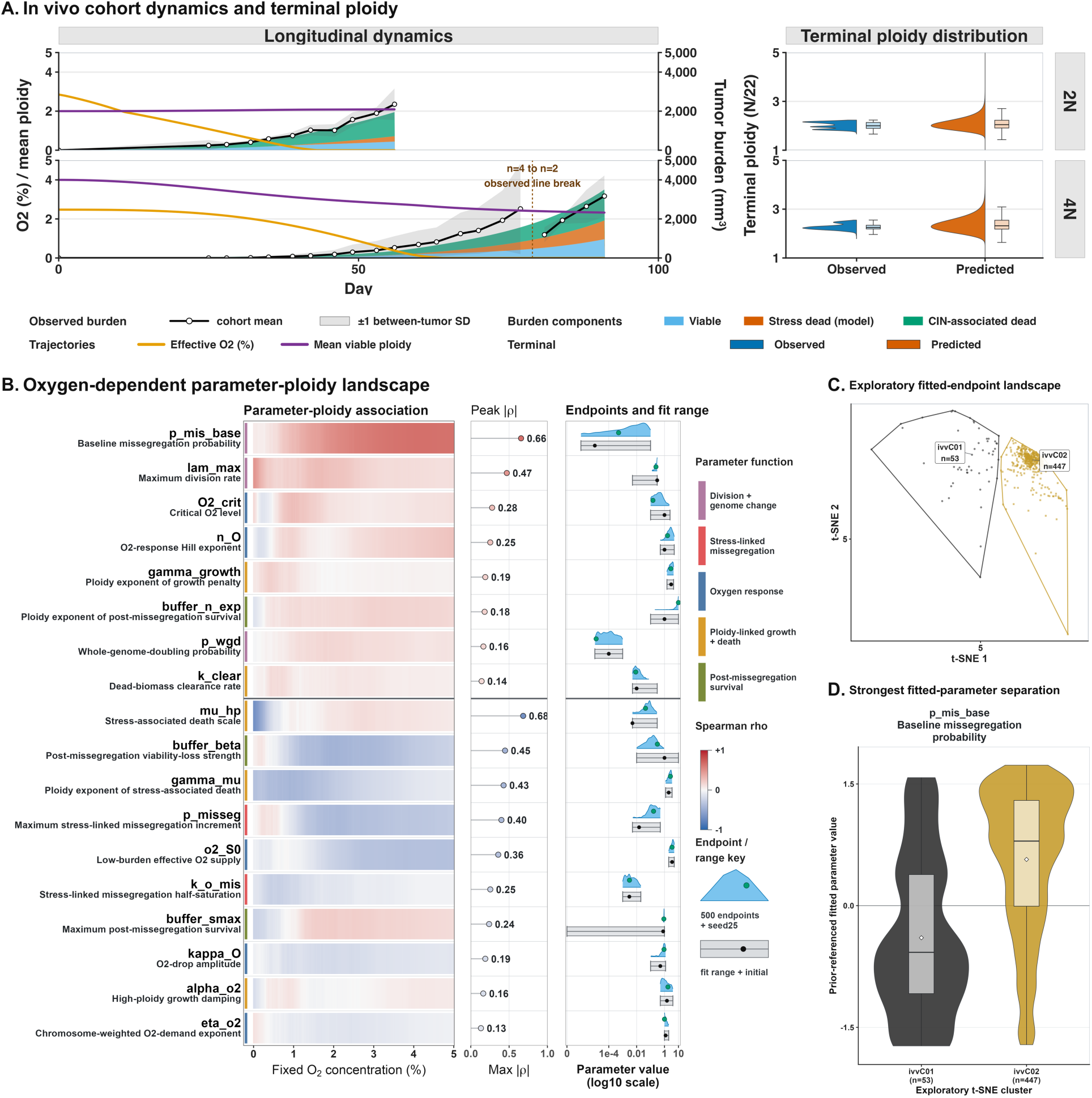
As oxygen increases, the strongest ploidy associations shift from stress-associated death to baseline missegregation. **(A)** Observed and predicted dynamics from the lowest-objective separate tumor fit (seed 25). Black points and gray ribbons show mean burden *±* between-tumor standard deviation (*n* = 4 mice per cohort; 4N *n* = 2 on days 81–91). Stacked areas partition predicted burden into viable, resource-stress-dead, and CIN-associated dead components (right axis, mm^3^). Latent effective oxygen (orange, %) and mean viable ploidy (purple, chromosome number/22) share the left axis. Terminal distributions compare RNA-derived observations (blue; 3,728 2N-derived and 2,774 4N-derived cells, four tumors per cohort) with predictions (orange), weighting tumors equally. **(B)** Spearman correlations between each of 18 fitted parameters and continuous mean asymptotic ploidy across 500 endpoints at 201 fixed oxygen concentrations (0–5%). Heatmap colors show signed correlations; dots show each parameter’s maximum absolute correlation, colored by its sign at that oxygen level. Red denotes positive and blue negative associations. Blue densities show fitted endpoint values; green points mark seed 25. Gray ranges and black points denote fitting bounds and initial values on a log_10_ scale. **(C)** t-distributed stochastic neighbor embedding (t-SNE) positions of fitted endpoints, with hulls outlining clusters of 53 (gray) and 447 (gold) solutions. **(D)** The largest contrast between the two solution clusters is in the fitted baseline mis-segregation rate (Kruskal–Wallis effect size *ε*^2^ among parameter comparisons meeting Benjamini–Hochberg *q <* 0.05). Panels B–D describe numerical-search variation, with cluster comparisons using the same parameters that defined the groups (Supplementary Tables 3 and 4). CIN, chromosomal instability.

To identify which modeled processes were most closely associated with ploidy at different resource levels, we evaluated each of the 500 fitted parameter sets at 201 fixed oxygen concentrations spanning 0–5%, without refitting. At each oxygen level, the normalized dominant eigenvector summarized the model’s long-term chromosome-state composition and its mean ploidy. We then examined each of 18 biological parameters separately, using Spearman correlation to ask whether larger parameter values accompanied higher or lower predicted ploidy (Figure 4B; Supplementary Section 1.3.1).

At the lowest oxygen level, the parameter controlling the overall strength of resource-stress-induced death (the stress-associated death scale, *µ*_hp_) had the strongest association with predicted ploidy: larger values accompanied lower ploidy (*ϱ* = *→*0.683 at 0% O_2_; Figure 4B). The parameter controlling how steeply this death hazard increases with chromosome number, *ϑ_µ_*, was also negatively associated with ploidy. At higher oxygen, the baseline probability of missegregation per chromosome copy per division, *p*_mis,base_, became the strongest correlate, with larger values associated with higher ploidy (peak *ϱ* = 0.656 near 4.2% O_2_; Supplementary Table 3). This pattern suggests a shift from ploidy outcomes dominated by differential survival under resource stress to outcomes shaped by proliferation, the segregation errors generated during division, and the survival of their products.

Multiple parameter combinations nevertheless remained compatible with the observed tumor trajectories. Clustering a two-dimensional t-SNE embedding of the transformed and standardized fitted parameters identified two groups of solutions (Figure 4C–D). Their strongest parameter contrast was *p*_mis,base_, with higher values in the larger group (Supplementary Table 4). This connects the cluster structure to the preceding oxygen analysis: the fitted regimes differed most along the same baseline-missegregation axis associated with ploidy at higher oxygen. Finite tumor-growth trajectories and harvest chromosome measurements can therefore constrain the observed remodeling while leaving alternative basal chromosome-error probabilities compatible with the data. These alternatives motivated joint fitting to determine which culture–tumor differences persisted across fitted regimes.

### Joint fits identify order-of-magnitude differences in stress-associated death and inducible missegregation between in vivo and in vitro environments

We next asked why the 2N lineage could expand a WGD-compatible component in culture but remain near diploidy in tumors, while the 4N lineage underwent chromosome loss in both settings. To distinguish differences in resource costs from differences in the survival of chromosome-altered descendants, we fitted both datasets jointly while penalizing cross-context parameter divergence. All 14 mechanistic parameters could take context-specific values, but their separation incurred a bounded penalty in the combined calibration objective. Two representative tumor solutions were paired with the same culture starting solution, generating families C01 and C02, each searched from 500 starts (Figure 5A, C; Supplementary Section 1.3.1, Eqs. (48)–(52)). The lowest-objective fit from each family retained the broad growth and ploidy patterns in both settings(Figure 5B).

**Figure 5:**
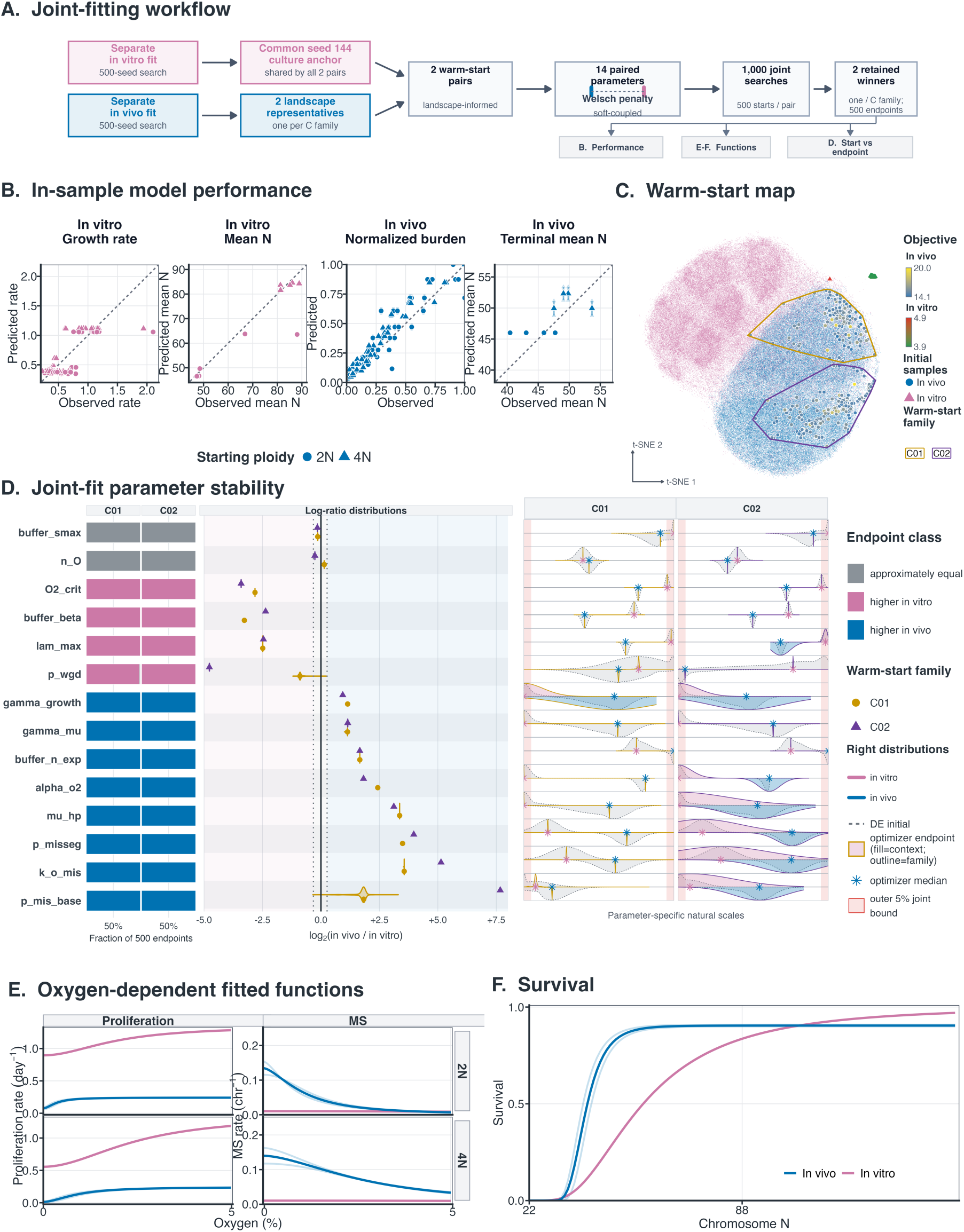
Joint fits distinguish higher baseline missegregation in vivo from faster proliferation and more frequent genome doubling in vitro. **(A)** Joint calibration penalizes differences between 14 paired mechanistic parameters while allowing context-specific values. C01 and C02 pair separate tumor fits from seeds 366 and 25, respectively, with the common culture seed-144 anchor. Five hundred starts per family yield 1,000 searches; B, E, and F use the lowest-objective winner from each family. **(B)** In-sample observed versus predicted culture growth, culture mean chromosome number, normalized tumor burden, and terminal tumor mean chromosome number. Pale points show the two winners; filled points and bars show their median and range. Circles and triangles distinguish 2N and 4N starts; dashed diagonals indicate equality. Blue and pink identify tumor and culture predictions. **(C)** Pooled 14-parameter t-distributed stochastic neighbor embedding (t-SNE) of separate-fit initial populations and endpoints used to select warm starts. Faded points show initial populations; circles and triangles denote tumor and culture endpoints, respectively, colored by objective value. Outlines identify the two tumorderived families. **(D)** Context contrasts across 500 endpoints per family. Left bars classify parameter ratios *r* = *Q*_vivo_*/Q*_vitro_ as higher in culture (*r ↓* 0.8, pink), approximately equal (0.8 *<r <* 1.2, gray), or higher in tumors (*r ↔* 1.2, blue). Central distributions show log_2_ *r*, with family medians marked by gold circles (C01) and purple triangles (C02); zero denotes equality. Right-hand distributions compare reconstructed differential-evolution initial populations (dashed) with fitted endpoints (filled); asterisks mark endpoint medians and shaded edge bands the outer 5% of each transformed fitting interval. These distributions summarize numerical searches, not posterior uncertainty. **(E)** Fitted proliferation rates (day*^→^*^1^) and complete per-chromosome missegregation probabilities, *p*_mis_(*N, O*_2_), at 2N (*N* = 44) and 4N (*N* = 88) over 0–5% O_2_. Common standardized oxygen coordinates do not imply equivalent resource environments. The evaluated probability differs from the maximal inducible increment, *p*_misseg_, scanned in Figure 6. **(F)** Per-copy survival factor *s_N_* versus maternal chromosome number, under the assumed nondecreasing relationship; survival after a shift of *m* copies is *s^m^*. In E–F, thin curves show the two family winners and thick curves their pointwise medians; blue denotes tumors and pink cultures (Supplementary Tables 7, 8, and 13).

Both families retained greater growth capacity in culture and stronger resource penalties in tumors (Figure 5D). Comparing median fitted parameter values, the maximum division rate was approximately 5.5-fold higher in culture, whereas the stress-associated death scale was 9–10-fold higher in tumors. Tumor fits also assigned stronger resource-dependent growth suppression and greater chromosome-number dependence to growth and death costs. Within the model, these differences increase the cost of maintaining chromosome-rich states in tumors, while the higher culture proliferative ceiling permits greater expansion when resource penalties are weak.

Stronger tumor resource stress was coupled to a larger capacity to generate chromosome errors, but not to a uniform increase in their effective probability. The maximal stress-induced missegregation increment was 11–16-fold higher in tumors, while the death hazard required to reach half of that increment was also 12–35-fold higher (Supplementary Table 8). Thus, the tumor fits allowed a larger inducible increase in errors but required a higher death hazard to reach the same fraction of that increase.

Although the fitted upper limits on survival were similar, chromosome-segregation errors were better tolerated in vivo, particularly near diploidy (Figure 5D, F). Following a one-chromosome change, predicted survival of daughters from near-diploid parents was 66–77% in tumors versus 20% in culture. For near-tetraploid parents, survival was more similar: approximately 90–91% in tumors versus 84% in culture (Supplementary Table 13). Thus, higher ploidy provided a substantially larger additional survival benefit in culture, whereas tumor fits already allowed relatively high survival of altered daughters from near-diploid parents.The fits also assigned a higher baseline per-chromosome missegregation probability to tumors and a higher probability of WGD per division to culture. Both differences were more pronounced in C02 than in C01, indicating agreement in their direction but variation in their magnitude between the two fitted families (Figure 5D; Supplementary Table 8). Together, these contrasts provide a coherent model-based explanation for the different in vivo ploidy trajectories. In tumors, stronger resource-related growth and death penalties can favor chromosome loss, while relatively permissive survival of missegregation products from near-diploid parents can help sustain low ploidy lineages. In culture, the higher proliferative ceiling, higher WGD probability, and larger survival gain associated with increasing ploidy can support expansion of the WGD-compatible component before its subsequent chromosome loss. Thus, the culture–tumor contrast reflects differences in both the resource costs of chromosome content and the retention of chromosome-altered descendants.

### The oxygen–ploidy relationship depends on missegregation and population history

The joint fits suggested that resource costs and ploidy buffering were balanced differently in culture and tumors. We therefore asked how these differences would alter ploidy when oxygen and the capacity for stress-induced missegregation changed. We varied oxygen and the maximal stressinduced increment, *p*_misseg_, while retaining each fit’s other parameters, first comparing long-term chromosome-state composition under fixed conditions and then following populations initialized at 2N or 4N (Figure 6A–B; Supplementary Table 9).

**Figure 6:**
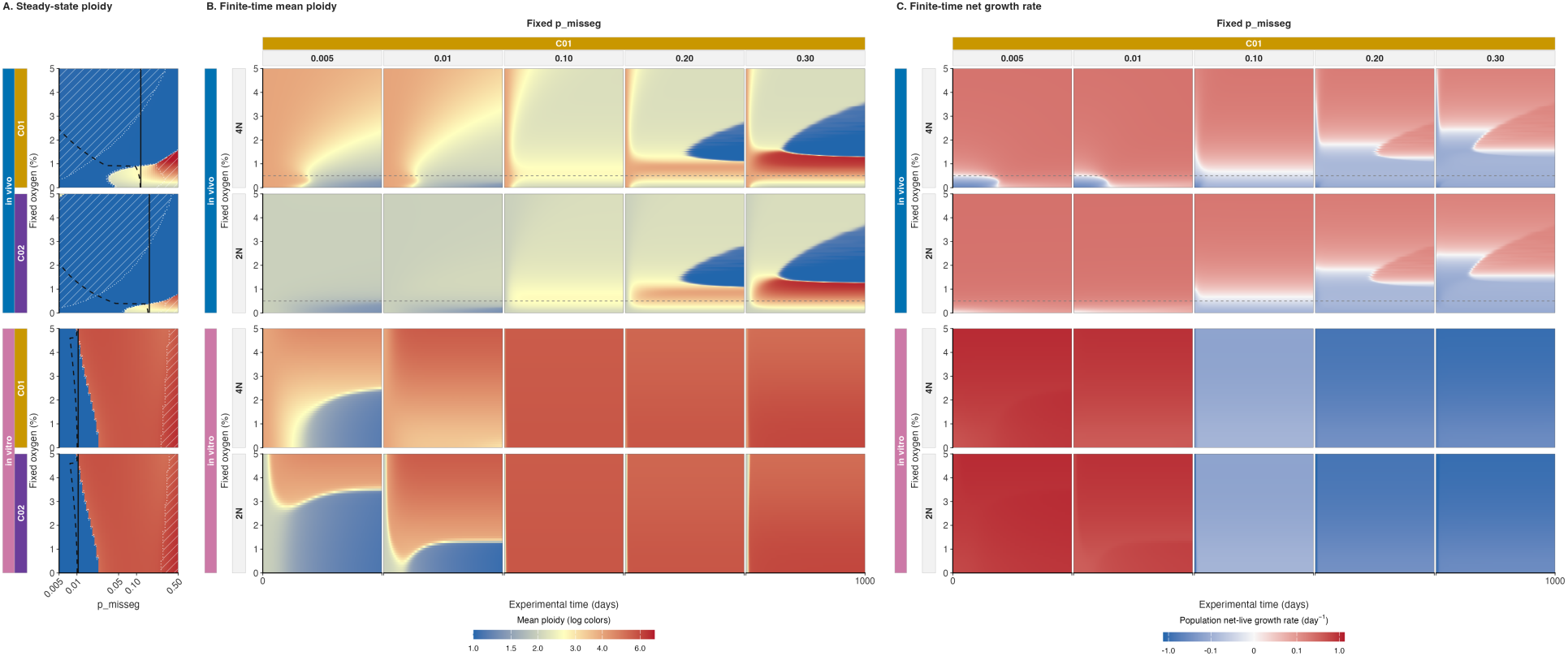
Missegregation alters the predicted oxygen–ploidy relationship differently in culture and tumors. (A-C) Ploidy composition evaluated at 201 fixed oxygen concentrations (0–5%) and 60 logarithmically spaced *p*_misseg_ inputs (0.005–0.5). Blue and pink side strips distinguish tumor and culture parameterizations. Panel A describes asymptotic composition; panels B–C retain initial-state and passage effects over a finite interval. **(A)** Color code reflects the seed-weighted median of mean ploidy calculated from each normalized dominant eigenvector. Black lines show ensemble means at fitted inputs: solid, the maximal increment *p*_misseg_; dashed, the population-averaged complete per-chromosome missegregation probability. The dashed reference is a different quantity from the scanned increment, not a fitted trajectory through that surface. White boundaries and hatching indicate weak separation between leading growth modes. **(B–C)** Finite-time simulations over 0–1,000 days from 4N and 2N initial populations, shown in upper and lower rows within each context. Tumor simulations use continuous growth; culture simulations include daily threshold-triggered stochastic passage with fixed-inoculum sampling. Horizontal dashed lines mark the 0.5% O_2_ reference. Color code reflects ploidy means across endpoints, after averaging stochastic repeats within endpoints **(B)** and population net-live growth rate (day*^→^*^1^), shown on a signed pseudo-log_10_ color scale with a 0.01 day*^→^*^1^ linear threshold **(C)**. Analysis design and robustness checks are reported in Supplementary Tables 9 and 10. All analyses use the lowest-objective decile of each joint-fit family (50 endpoints for C01), retaining fitted parameters except the scanned maximal stress-induced missegregation increment, *p*_misseg_.

Culture and tumor parameterizations produced distinct responses. Increasing *p*_misseg_ produced relatively sharp transitions from low to high dominant mean ploidy in culture, with similar patterns in C01 and C02. Tumor surfaces instead contained broader intermediate-ploidy regions whose extent and boundaries varied with oxygen and fit family (Figure 6A; Figure 8).

We next asked how rapidly populations starting at different ploidies approached the predicted long-term compositions. Tumor trajectories developed oxygen-dependent bands of low, intermediate, and high mean ploidy, with family-dependent timing and boundaries. Culture passage simulations showed similar family patterns, but trajectories from 2N and 4N initial states could remain distinct over the 1,000-day horizon, particularly at lower scanned missegregation increments (Figure 6B). Starting ploidy could therefore remain consequential even when the fixed-condition model specified a common long-term composition. In the continuous-propagation comparison, agreement with the asymptotic composition generally improved with elapsed time, although some conditions remained separated at day 1,000 (Supplementary Figure 9).

We then examined whether these ploidy compositions were accompanied by population expansion or decline. Paired maps of mean ploidy and net live-cell growth showed that, when mean ploidy was above four, growth was negative in both tumor and culture projections (Figure 6B–C; Supplementary Figure 4A–B). These high-ploidy compositions therefore characterized declining populations. Conditions combining low oxygen with high missegregation were particularly prone to population decline and extinction. The corresponding long-horizon growth maps for continuous tumor propagation and passage-aware culture simulations are provided in Supplementary Figures 11 and 14.

Because culture populations are repeatedly diluted and sampled, we also compared passage-aware and continuous culture projections. Continuous culture projections retained oxygen-dependent low/high-ploidy regions across a wider domain, including high-oxygen low-ploidy regions for selected inputs (Supplementary Figures 8, 12, and 13).

The qualitative patterns persisted across the tested optimizer-eligibility cutoffs and duplicate-endpoint weighting choices (Supplementary Figures 6 and 7; Supplementary Table 10). Detailed grids, ensemble definitions, and diagnostics are given in the Supplementary Methods.

Taken together, these analyses predict sharper transitions between low- and high-ploidy outcomes in culture and broader intermediate-ploidy regions in tumors, while starting ploidy and passaging influence which outcomes are reached over time. In the tumor projections at lower scanned missegregation increments, higher oxygen was associated with higher mean ploidy, directionally consistent with the reported association between tissue oxygenation and the ploidy of cancers arising in those tissues^4^. At higher increments, however, the oxygen–ploidy relationship became non-monotonic, with ploidy declining again as oxygen increased. One possible explanation is that proliferating high-ploidy cells continually generate viable chromosome-loss descendants, replenishing lower-ploidy compartments even under conditions that support high-ploidy growth. This mechanism could lower population-average ploidy without requiring the elimination of the high-ploidy population. Thus, resource availability does not specify a single ploidy outcome: its effects depend on the generation and survival of chromosome variants and on the population’s evolutionary history.

## Discussion

Our findings frame ploidy evolution as a balance between the resource costs of carrying additional chromosomes and the protection those chromosomes can provide against segregation errors. Integrating matched *in vitro* and *in vivo* datasets into a common mechanistic framework revealed how this balance changes between culture and tumors. Joint fits assigned tumors a lower proliferative ceiling, stronger stress-associated death, and a greater capacity for stress-induced missegregation. Baseline per-chromosome missegregation was also higher in vivo, whereas the probability of WGD was higher in vitro, with the magnitude of these contrasts varying between fitted families. Together, these differences suggest that tumor environments reshape both the production of chromosome-number variation and the fitness consequences of carrying it.

In vivo, altered daughters from near-diploid parents survived more readily than in culture, whereas higher ploidy provided a larger additional survival benefit in culture. This difference arose despite similar fitted upper limits on survival after a one-chromosome segregation error. The parameter controlling how sharply resource stress rises as oxygen falls was also approximately shared between contexts.

The ecological meaning of oxygen nevertheless differs between these settings. Experimentally assigned culture oxygen and the model’s inferred tumor oxygen represent different resource environments: anoxia in glucose-rich culture medium differs from tumor hypoxia accompanied by impaired perfusion and simultaneous shortages of other nutrients. Under this interpretation, the stronger fitted tumor responses reflect composite physiological stress, potentially including heterogeneous or episodic exposure, rather than an intrinsically stronger response to oxygen deprivation alone.

Within tumors, the oxygen-dependent parameter associations suggest a shift from death-dominated to proliferation-dominated ploidy evolution. At low oxygen, the most consequential difference between chromosome states is their ability to survive resource stress. Reducing chromosome burden can therefore be advantageous even when it diminishes protection against segregation errors. At higher oxygen, resource-associated death subsides, shifting the emphasis from avoiding death to producing viable descendants through proliferation. Both the rate of division and the chromosome errors generated during division then become more consequential. Ongoing missegregation makes the ability to sustain growth despite these errors an important determinant of ploidy composition. Both regimes therefore balance the benefits of chromosome-number reduction against the loss of protection from missegregation, but the dominant pressure shifts from surviving resource stress to successful proliferation. Fitted solutions on either side of this balance can reproduce the observed trajectories over the experimental interval while predicting different longer-term ploidy outcomes. Across tissues, this outcome can also be shifted by alternative processes not accounted for by the presented model. Differences in WGD generation, physiological polyploidization programs, immune-mediated selection, and tissue-specific mutagenic pressures provide complementary explanations for variation in tumor ploidy. A tissue with frequent polyploidization or mitotic errors may generate WGD-positive cells more often, while immune pressure or mutagenic exposure may change their subsequent advantage. These processes act on the same two components distinguished by the model: the generation of chromosome variants and their survival and expansion.

Our model does not explicitly account for differences in immune susceptibility among cells of different ploidy. This limitation is partly mitigated by the use of immunocompromised mice, although residual innate immune activity cannot be excluded solely on this basis. Experimentally induced polyploidization can enhance NK-cell recognition, while associated centrosome amplification can promote NK-mediated clearance and alter macrophage inflammatory responses^25,26^. Moreover, macrophages retain phagocytic potential even in profoundly immunodeficient NSG hosts^27^. These findings do not establish differential immune clearance of the near-diploid and near-tetraploid populations studied here, but leave open a contribution of residual immune selection to the inferred survival differences. Extending the framework to immunocompetent tumors may therefore require explicit consideration of ploidy-associated immune interactions.

This distinction has direct implications for therapies that increase chromosome-segregation errors. Anticancer agents can induce such errors by disrupting mitosis or by generating replication-associated and DNA-damage lesions that persist into mitosis^28–34^. Within the modeled survival framework, near-diploid cells are more vulnerable to the dosage consequences of individual chromosome losses, whereas higher-ploidy cells can retain some altered daughters as heritable karyotype variation. The therapeutic consequence of increasing missegregation should therefore depend on both the error burden imposed and the buffering advantage available in the treated environment.

The next step is to deploy and extend this framework to predict responses to drugs that induce chromosome missegregation and interventions that modify the tumor microenvironment. Incorporating drug-specific effects on segregation alongside treatment-induced changes in resource availability would allow us to generate testable predictions of how these perturbations jointly affect tumor growth and ploidy evolution. A central question is when increased missegregation overwhelms cellular buffering and when it instead generates viable variants capable of expansion. These predictions have the potential to guide testing of treatment combinations and schedules that couple chromosome instability to environmental constraints, steering populations toward states with reduced fitness or greater susceptibility to subsequent treatment. This connects the development of testable predictions of ploidy evolution with the prospect of directing it: exploiting both the context-dependent resource demands of chromosome-rich cells and the potential dosage vulnerability of lower-ploidy cells to further chromosome losses..

## Code and Data availability

Previously published karyotype and growth data for both in-vitro and in-vivo settings was accessed through the CLONEID database (dev.cloneid.org). The code used for model fitting and analysis is available at https://github.com/TaoLee0510/LTEE-Oxygen-Model. The corresponding modelfitting results and intermediate analysis data are archived in Zenodo (DOI: https://doi.org/10.5281/zenodo.22799665).

## Methods

### Study design and data streams

We studied near-diploid (2N) and near-tetraploid (4N) SUM159 breast cancer lineages in complementary *in vivo* and *in vitro* experiments. Eight untreated xenografts (four per starting-ploidy background) contributed longitudinal tumor-burden measurements and terminal RNA-derived chromosome-number estimates; terminal necrotic fractions were available for six tumors. Cultures were serially passaged under a defined oxygen schedule while population expansion, direct chromosome counts, and flow-cytometric ploidy profiles were measured. O1 and O2 denote separately propagated oxygen-deprivation lineages within each starting-ploidy cohort, not established independent biological replicates. These data support a lineage-matched cross-context comparison, not an estimate of a randomized environmental effect. Experimental procedures and observation inventories are described in the Supplementary Information.

### Model state and oxygen context

Viable cells are indexed by total autosomal chromosome number, from 22 to 154, with 44 chromosomes as the diploid reference. Two additional compartments track retained dead-cell populations assigned to resource-stress death and to nonviable chromosome-transition products. Tumor burden is linked to the sum of live and dead compartments, whereas chromosome profiles are linked to the viable distribution. Tumor oxygen is a fitted, time-dependent mean-field variable; culture oxygen is externally assigned and piecewise constant within passage segments. Neither representation is a direct measurement of intracellular oxygen.

### Mechanistic modules

Oxygen-linked stress modifies state-specific division and death rates. The per-chromosome missegregation probability, *p*_mis_(*N, O*_2_), depends on chromosome state and stress; its baseline, *p*_mis,base_, and maximal stress-induced increment, *p*_misseg_, are distinct fitted quantities. A separate constant per-division WGD probability allocates a division event to an *N* → 2*N* conversion. The per-copy survival factor *s_N_* weights missegregated daughter products, with survival 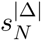 for a count shift Δ. This parameterization assumes nondecreasing survival with maternal chromosome number; calibration estimates its magnitude and ploidy dependence. Out-of-grid products enter the chromosome-transition dead pool, so that compartment includes finite-domain loss as well as modeled biological nonviability.

### Observation models

Burden, terminal chromosome distributions, and available terminal necrosis contribute separate tumor-calibration terms. Culture growth, direct chromosome counts, and flow-density profiles are compared with passage-level predictions under their respective observation models. The culture comparison state is selected by proximity to the recorded cell-count target, as detailed in the Supplementary Methods; it is not automatically the state at the recorded passage duration. Components measured in the same tumor or culture lineage constitute related evidence.

### Joint calibration

Fourteen mechanistic parameters were soft-coupled, allowing context-specific values under a bounded Welsch penalty with transformed-scale coupling scale 0.65 and saturation constant 0.4. Two tumor warm-start families, C01 and C02, used separate-fit seeds 366 and 25, respectively, and the common culture seed 144. Each family had 500 optimization starts. Main fitted-function comparisons use the lowest-objective retained winner from each family; optimization details, bounds, and endpoint distributions are reported in Supplementary Tables 7 and 8.

### Post-fit analyses and interpretation

Fixed-oxygen scans evaluate the normalized dominant eigenvector of the live-state operator, whereas finite-time simulations retain the specified initial state and continuous or passage-aware propagation rule. A stationary normalized composition does not by itself establish a nonzero equilibrium population size. These scans are model counterfactuals, not additional oxygen-intervention observations. Ensemble cutoffs, duplicate-endpoint handling, and weak-gap diagnostics assess numerical-search robustness within the selected families, not posterior uncertainty, biological replication, or unique identification of a mechanism.

