## supplementary figures, tables and methods for "Resource limitation rewires chromosome instability and ploidy evolution across in vitro and in vivo cancer models"

### 1 Supplementary Information

#### 1.1 *In vivo* experiments

The *in vivo* analysis used near-diploid and near-tetraploid variants of the SUM159 triple-negative breast cancer cell line, denoted `SUM159_NLS_2N_A6M` and `SUM159_NLS_4N_A4M`, respectively. For each tumor, $1 \times 10^6$ cells were injected orthotopically into mice. The subset analyzed here comprised eight tumors: four initiated from the near-diploid 2N line and four initiated from the near-tetraploid 4N line (Supplementary Table 1).

- **2N cohort:** four mice injected with `SUM159_NLS_2N_A6M`.
- **4N cohort:** four mice injected with `SUM159_NLS_4N_A4M`.

Tumors were harvested at predefined endpoints, and single-cell RNA sequencing was performed on all eight tumors. Chromosome-number distributions used for model calibration were derived from these endpoint single-cell profiles and analyzed together with the corresponding tumor-burden trajectories. Only this untreated subset was used for the modeling analyses presented here. Processed data and associated experimental metadata were organized and retrieved through CLONEID, a framework for structuring clone-specific genotypic and phenotypic measurements[35].

Supplementary Table 1: Experimental data inventory for the analyzed SUM159 culture and tumor datasets. Counts distinguish biological specimens, measurement records, and display summaries. Individual burden measurements, chromosome estimates, culture counts, passage records, and flow-density values are provided in the accompanying machine-readable supplementary tables.

| Setting | Data stream or design quantity | Biological or measurement units | Count or value | Scope and interpretation |
|---|---|---|---|---|
| *In vivo* | Orthotopic inoculum | Cells per tumor | $1 \times 10^6$ | Experimental input for each analyzed tumor |
| *In vivo* | Untreated tumor cohort | Tumors | 8 | Four near-diploid initiated and four near-tetraploid initiated tumors |
| *In vivo* | Longitudinal burden | Tumor measurements | 120 included; 112 in burden loss | Records from the eight tumors within the configured cohort and time scope; the eight day-0 records were excluded from the burden-loss calculation |
| *In vivo* | Terminal chromosome-number profiles | Single-cell estimates | 6,502 | 3,728 estimates from four near-diploid initiated tumors and 2,774 from four near-tetraploid initiated tumors |
| *In vivo* | Matched terminal necrosis | Tumor-level fractions | 6 | Three tumors per starting-ploidy cohort had matched necrosis measurements used by the observation model |
| *In vitro* | Passage-structured fit records | Passage entries | 131 | All entries retained assigned oxygen, timing, and linked population measurements |
| *In vitro* | Passage growth | Finite estimates | 114 | Entries with a finite segment-level growth estimate |
| *In vitro* | Direct chromosome counts | Samples; cells | 12; 220 | Passage-linked chromosome-number samples and their total single-cell observations |
| *In vitro* | G0/G1 DNA-content profiles | Profiles; grid points per profile | 20; 200 | Normalized profiles used in the flow-density comparison |
| *In vitro* | Oxygen schedule | Target $O_2$ values (%) | 20.5, 2, 1, 0.5, 0.3, 0.2, 0.1, 0 | Control and serial deprivation levels; a post-anoxia branch was evaluated at 1% |
| Figure 1 display | Culture population markers | Displayed endpoints | 77 | Representative cohort, condition, and reconstructed-day summaries |
| Figure 1 display | Flow summaries | Profiles; grouped time points | 20; 12 | Individual profile peaks and equal-weight grouped density summaries |
| Figure 1 display | Ploidy conversion | Autosomal chromosomes per ploidy unit | 22 | Used only to express chromosome counts on the plotted ploidy scale |
| Figure 1 display | Flow plotting viewport | Ploidy range; minimum retained mass | 1–5; 85.43% | Full normalized profiles were used for calculation; only the display viewport was cropped |
| Figure 1 display | Starting chromosome counts | Directly counted cells | 40 | Twenty cells from each starting-ploidy cohort |

*Note:* Counts summarize the analyzed data and do not imply equal replication across data streams. "Included" denotes records inside the configured cohort and time scope, whereas "in burden loss" denotes records that contribute to the longitudinal burden objective. Display counts can exceed the number of unique biological samples when a shared starting distribution is repeated for alignment. The machine-readable companion tables preserve the individual observations and identifiers needed to reconstruct these totals.

Supplementary Table 2: Absolute fit-quality metrics for the ten lowest-objective separate *in vivo* fits from the current 500-seed run. Burden errors are calculated on the log-burden scale; terminal chromosome-number and necrosis metrics are averaged equally across tumors or harvests.

| Rank | Fit | Objective | Burden log-RMSE | Tumor-balanced log-RMSE | Terminal mean-$N$ MAE | Terminal Wasserstein-1 | Terminal total variation | Necrosis-fraction MAE |
|---|---|---|---|---|---|---|---|---|
| 1 | `seed25` | 14.0508 | 0.6677 | 0.6330 | 3.03 | 4.38 | 0.682 | 0.083 |
| 2 | `seed74` | 14.1304 | 0.6673 | 0.6333 | 3.38 | 4.83 | 0.699 | 0.089 |
| 3 | `seed366` | 14.1358 | 0.6676 | 0.6337 | 2.51 | 4.74 | 0.708 | 0.079 |
| 4 | `seed292` | 14.1372 | 0.6724 | 0.6384 | 2.79 | 4.59 | 0.698 | 0.064 |
| 5 | `seed392` | 14.1466 | 0.6790 | 0.6433 | 3.51 | 4.92 | 0.686 | 0.093 |
| 6 | `seed165` | 14.1469 | 0.6789 | 0.6430 | 3.43 | 4.84 | 0.684 | 0.094 |
| 7 | `seed391` | 14.1547 | 0.6796 | 0.6438 | 3.41 | 4.76 | 0.682 | 0.091 |
| 8 | `seed322` | 14.1613 | 0.6688 | 0.6340 | 3.01 | 4.96 | 0.712 | 0.078 |
| 9 | `seed6` | 14.1782 | 0.6783 | 0.6422 | 3.53 | 5.12 | 0.697 | 0.095 |
| 10 | `seed264` | 14.1786 | 0.6726 | 0.6395 | 2.43 | 4.18 | 0.684 | 0.081 |

*Note:* Positive-observation burden RMSE weights every retained measurement equally; tumor-balanced RMSE first averages squared residuals within each tumor. Wasserstein-1 and mean-$N$ MAE are in chromosome units. Necrosis fraction is reconstructed at the mapped harvest day as predicted dead volume divided by predicted total burden. These optimizer-selected fits are not biological replicates or confidence-interval samples.

Supplementary Table 3: Peak continuous fixed-$O_2$ fitted-parameter associations underlying Figure 4B. Parameters are grouped by the sign of the Spearman correlation at their maximum absolute association, with the positive group shown before the negative group, and ordered by decreasing maximum $|\rho|$ within each group. Signed peak $\rho$ and peak $O_2$ report the first oxygen value attaining that maximum. Sign stability is the percentage of oxygen values sharing the predominant correlation sign and is not used for ordering.

| Within-group rank | Parameter | Max $|\rho|$ | Signed peak $\rho$ | Peak $O_2$ (%) | Sign stability (%) |
|---|---|---|---|---|---|
| *Positive peak $\rho$* | | | | | |
| 1 | `p_mis_base` | 0.656 | +0.656 | 4.175 | 100.0 |
| 2 | `lam_max` | 0.470 | +0.470 | 0.025 | 100.0 |
| 3 | `O2_crit` | 0.278 | +0.278 | 0.975 | 92.5 |
| 4 | `n_0` | 0.254 | +0.254 | 5.000 | 90.5 |
| 5 | `gamma_growth` | 0.189 | +0.189 | 0.400 | 99.5 |
| 6 | `buffer_n_exp` | 0.180 | +0.180 | 3.025 | 94.5 |
| 7 | `p_wgd` | 0.165 | +0.165 | 1.800 | 97.5 |
| 8 | `k_clear` | 0.141 | +0.141 | 0.450 | 100.0 |
| *Negative peak $\rho$* | | | | | |
| 1 | `mu_hp` | 0.683 | -0.683 | 0.000 | 83.6 |
| 2 | `buffer_beta` | 0.447 | -0.447 | 2.050 | 93.0 |
| 3 | `gamma_mu` | 0.428 | -0.428 | 0.900 | 100.0 |
| 4 | `p_misseg` | 0.400 | -0.400 | 2.125 | 85.6 |
| 5 | `o2_S0` | 0.358 | -0.358 | 3.825 | 97.5 |
| 6 | `k_o_mis` | 0.254 | -0.254 | 0.450 | 100.0 |
| 7 | `buffer_smax` | 0.239 | -0.239 | 0.000 | 83.6 |
| 8 | `kappa_0` | 0.189 | -0.189 | 2.025 | 100.0 |
| 9 | `alpha_o2` | 0.163 | -0.163 | 0.000 | 68.7 |
| 10 | `eta_o2` | 0.132 | -0.132 | 2.450 | 93.0 |

*Interpretation:* The display group uses the sign at maximum $|\rho|$, which can differ from the predominant grid sign when a strong association is localized to a narrow oxygen range. For example, `mu_hp` has a negative peak at 0% $O_2$ although 83.6% of its grid correlations are positive. The full signed heatmap, rather than the grouped peak order alone, should therefore be used to assess oxygen dependence. The gray fitting ranges, black initial values, endpoint densities, and seed25 markers in Figure 4B are optimizer-derived quantities, not posterior samples, credible intervals, measurement bounds, or biological confidence intervals.

Supplementary Table 4: Separate *in vivo* fitted-landscape and numerical-search summary. Cluster comparisons are descriptive because the groups were constructed from the same fitted parameters subsequently compared. Optimizer endpoints are numerical solutions, not biological replicates or posterior samples.

| Analysis | Quantity | Value | Definition or interpretation |
|---|---|---|---|
| Fixed-$O_2$ association audit | Evaluation dimensions | $500 \times 18 \times 201$ | 1,809,000 fitted-solution–parameter–oxygen evaluations from 0% to 5% $O_2$ |
| Embedding input | Numerical rows | $127,500 + 500$ | Retained initial-population rows plus fitted endpoints; 18 fitted parameters were embedded |
| Exploratory clustering | C01, C02 sizes | 53, 447 | Endpoint counts in the selected two-group solution |
| Exploratory clustering | Selected $k$; average silhouette | 2; 0.685 | Descriptive separation on the saved *in vivo*-only embedding |
| Omnibus comparison | BH-significant parameters | 11 of 18 | Kruskal–Wallis tests with Benjamini–Hochberg $q < 0.05$ |
| Top cluster contrast | Baseline missegregation probability, $p_{\text{mis,base}}$ | 0.079 | Kruskal–Wallis epsilon-squared; BH $q = 4.35e-09$ |
| Top cluster contrast | Post-missegregation-survival exponent, $n_{\text{exp}}$ | 0.072 | Kruskal–Wallis epsilon-squared; BH $q = 1.3e-08$ |
| Top cluster contrast | Oxygen-response Hill exponent, $n_O$ | 0.048 | Kruskal–Wallis epsilon-squared; BH $q = 4.09e-06$ |
| Top cluster contrast | Whole-genome-doubling probability, $p_{\text{wgd}}$ | 0.038 | Kruskal–Wallis epsilon-squared; BH $q = 3.69e-05$ |
| Top cluster contrast | Oxygen-drop amplitude, $\kappa_O$ | 0.032 | Kruskal–Wallis epsilon-squared; BH $q = 1.48e-04$ |
| Top cluster contrast | Maximum post-missegregation survival, $s_{\max}$ | 0.030 | Kruskal–Wallis epsilon-squared; BH $q = 1.95e-04$ |
| Objective proximity | Starts within 1%, 5%, and 10% | 13, 225, 391 of 500 | Relative excess above the lowest retained objective |
| Optimizer record | DE termination flag | 500 of 500 | Implementation-specific differential-evolution stopping flag |
| Optimizer record | Local refinement attempted; accepted | 500; 460 | Recorded local-refinement outcomes |
| Optimizer record | Local convergence code 0 | 281 | All endpoints carrying convergence code 0 |
| Selected solution | Active configured bounds | 1 of 20 | Exact active-bound count for the lowest-objective separate fit, seed 25 |

#### 1.2 *In vitro* experiments

The *in vitro* analysis used matched near-diploid (2N) and near-tetraploid (4N) SUM159 lineages. The O1 and O2 nomenclature defined in the main Methods is also used in Figures 1 and 3: O1 and O2 denote the two separately propagated oxygen-deprivation lineages within each starting-ploidy cohort. For each starting ploidy, a control branch was maintained at 20.5% $O_2$, and O1 and O2 were stepped over successive passages through target levels of 2%, 1%, 0.5%, 0.3%, 0.2%, 0.1%, and 0% $O_2$. The main deprivation branch was then maintained at target 0% $O_2$; an additional branch after the first 0% passage was evaluated at target 1% $O_2$. We describe O1 and O2 as parallel lineages because the available records do not establish whether they should be classified as biological replicates (Supplementary Table 1).

For calibration, each fit entry retained the assigned oxygen level, passage duration, initial and final corrected cell counts, and a segment-level growth-rate estimate. Chromosome-number mea-

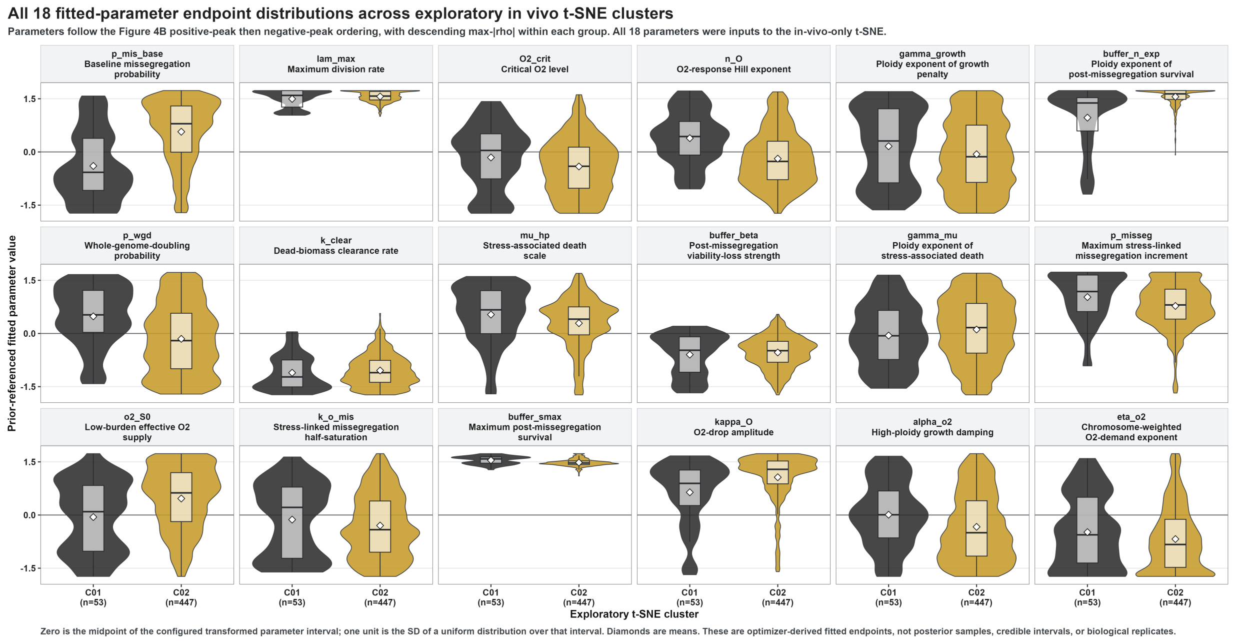


Supplementary Figure 1: **All 18 fitted-parameter endpoint distributions across the exploratory separate-*in vivo* t-SNE clusters.** The 18 facets are arranged in three rows and six columns in the same positive-peak then negative-peak order as Figure 4B, with decreasing maximum $|\rho|$ within each sign group. Each facet shows the prior-referenced fitted-value distribution for the selected exploratory clusters C01 (53 endpoints) and C02 (447 endpoints); boxplots show medians and interquartile ranges and diamonds show means. Figure 4D displays the six-parameter subset selected from these same values by the prespecified omnibus-test and effect-size rule. All displayed parameters contributed to the *in vivo*-only 18-parameter t-SNE and endpoint clustering. Zero is the midpoint of the configured transformed parameter interval and one unit is the standard deviation of a uniform distribution over that interval. Cluster sizes, tests, and effect sizes are reported in Supplementary Table 4. These optimizer-derived distributions quantify internal fitted-landscape structure rather than posterior uncertainty or biological replication.

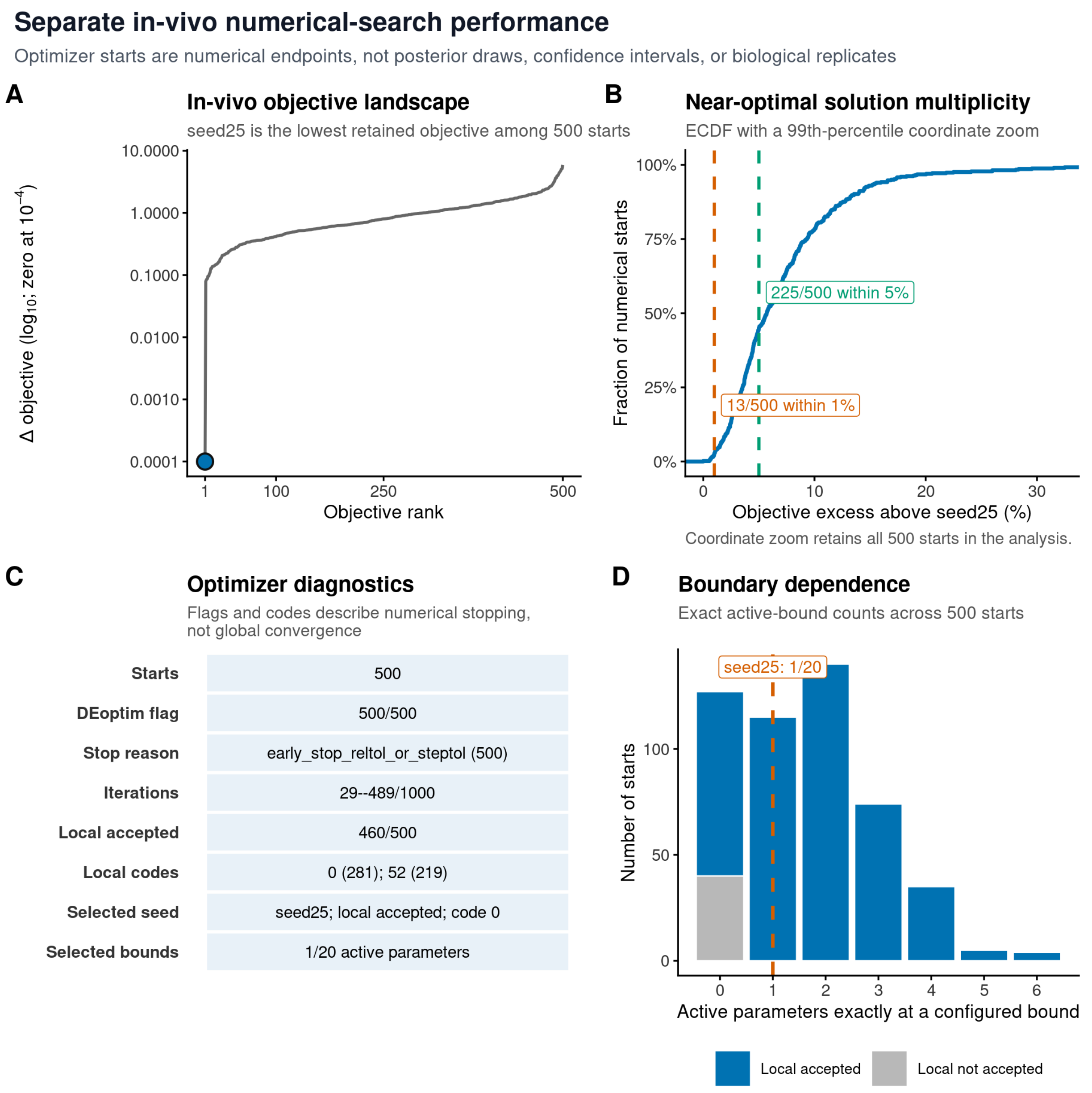


Supplementary Figure 2: **Complete numerical-search diagnostics for the separate *in vivo* fit ensemble.** **(A)** Objective difference by rank across 500 starts, with the lowest retained fit, seed 25, highlighted. **(B)** Empirical cumulative distribution of relative objective excess above seed 25. Vertical markers show that 13 starts were within 1% and 225 were within 5%; the displayed x-range ends at the 99th percentile by coordinate zoom without removing starts from the analysis. **(C)** Recorded differential-evolution termination, iteration, local-refinement, selected-seed, and boundary diagnostics. Flags and codes summarize implementation-specific numerical stopping. **(D)** Distribution of exact active-parameter boundary counts across all 500 retained endpoints, separated by local-refinement acceptance status; the dashed line marks seed 25, for which one of 20 active parameters was exactly at a configured bound. The corresponding counts are reported in Supplementary Table 4. The optimizer starts quantify numerical-search behavior rather than posterior uncertainty or biological replication.

surements and G0/G1 DNA-content density profiles were linked to the corresponding passage when available. The calibration dataset contains 131 entries, including 114 with finite growth estimates, 12 with chromosome-number measurements (220 single-cell values in total), and 20 with matched G0/G1 profiles. Each G0/G1 profile was represented on a 200-point ploidy grid and normalized before comparison with the simulated viable chromosome-number distribution (Supplementary Table 1).

For the Figure 1 timeline, population-size measurement availability was audited against the raw passage snapshot: all 131 passage IDs had a positive seeding count and at least one positive linked harvest count. The displayed representative tracks collapse matching records to cohort, condition, and reconstructed day; common pre-branch endpoints are repeated on both aligned eventual branches. For panel B, direct autosomal chromosome counts were converted to ploidy by division by $N_{\text{unit}} = 22$. For panel C, each observed flow profile was normalized to unit density mass. Profiles from parallel O1 and O2 lines at the same cohort, condition, and reconstructed day were averaged with equal sample weight for the half-violin and density-weighted box summary, while the global density maximum of each individual profile supplied its open-triangle marker. The complete normalized profiles were used to calculate peaks, quantiles, and whiskers before applying the ploidy 1–5 plotting viewport. This viewport retained at least 85.43% of normalized density mass in every sample; density above ploidy 5 was cropped without within-window renormalization (Supplementary Table 1). These display summaries did not alter the flow-density likelihood.

Within the passage-structured model, oxygen was held fixed at the assigned value for each segment. For segments represented by parallel records, passage duration and endpoint cell counts were summarized by the median, while all available growth, chromosome-number, and flow-density observations contributed to their corresponding likelihood terms.

Supplementary Table 5: Final karyotype distributions of the parallel 2N oxygen-deprivation lines and their matched fitted states from the current lowest-objective separate *in vitro* fit, seed144. O1 and O2 were propagated as separate lines from the same 2N precursor through the matched nominal oxygen schedule.

| Distribution | $n$ | Mean $N$ | Median $N$ | $N \geq 80$ | Component structure |
|---|---|---|---|---|---|
| O1 observed | 20 | 66.85 | 49 | 8/20 (40.0%) | 12 cells at $N = 46$–$49$; 8 cells at $N = 90$–$100$ |
| O2 observed | 20 | 88.05 | 95 | 16/20 (80.0%) | 4 cells at $N = 49$–$76$; 16 cells at $N = 93$–$99$ |
| O1 fitted state | — | 64.37 | 61 | 32.7% probability | dominant local modes at $N = 40$ and $N = 83$ |
| O2 fitted state | — | 64.17 | 60 | 32.4% probability | dominant local modes at $N = 41$ and $N = 83$ |

*Note:* $N \geq 80$ is a descriptive bin within the unoccupied interval between the low- and high-chromosome components of the final observed samples; it is not a fitted or inferential cutoff. Observed entries are single-cell counts. Fitted entries are probability distributions and therefore have no observed sample size. The current fit retains distinct O1 and O2 segment identifiers; their displayed summaries agree after rounding but they are audited separately.

Supplementary Table 6: Modeled 4N chromosome-number decline and the fitted contribution of the compartment representing death associated with resource stress across the ten lowest-objective separate *in vitro* fits. Initial and terminal mean chromosome numbers refer to the first and last modeled states of the selected 4N lineage; its terminal portion was maintained at target 0% $O_2$. The last column is the maximum daily burden assigned by the model to death associated with resource stress, expressed as a percentage of total burden over the full modeled 4N trajectory.

| Rank | Fit | Objective | Initial mean $N$ | Terminal mean $N$ | Decline in mean $N$ | Maximum modeled stress death burden (%) |
|---|---|---|---|---|---|---|
| 1 | `seed144` | 3.8556 | 84.25 | 80.69 | 3.56 | 1.77 |
| 2 | `seed325` | 3.8563 | 84.94 | 80.60 | 4.34 | 1.72 |
| 3 | `seed180` | 3.8566 | 84.58 | 80.62 | 3.95 | 1.76 |
| 4 | `seed362` | 3.8568 | 84.47 | 80.60 | 3.87 | 1.75 |
| 5 | `seed477` | 3.8568 | 84.38 | 80.59 | 3.78 | 1.69 |
| 6 | `seed475` | 3.8571 | 84.60 | 80.85 | 3.74 | 1.76 |
| 7 | `seed372` | 3.8572 | 84.45 | 80.60 | 3.85 | 1.75 |
| 8 | `seed64` | 3.8572 | 84.60 | 80.79 | 3.80 | 1.76 |
| 9 | `seed166` | 3.8573 | 84.56 | 80.86 | 3.70 | 1.76 |
| 10 | `seed207` | 3.8573 | 84.64 | 80.69 | 3.96 | 1.76 |

| Seed-144 audit quantity | Value | Scope |
|---|---|---|
| Maximum passage-end stress-associated dead-burden fraction | 2N: 1.06%; 4N: 1.73% | Severe-deprivation passage-end summaries shown in Figure 3F |
| Numerical starts and fitted parameters | 500; 20 | Separate *in vitro* calibration |
| Differential-evolution convergence flags | 500 of 500 | Recorded relative-tolerance/step-tolerance stop criterion |
| Selected local refinement | accepted, code 1 | Lowest-objective retained seed 144 |
| Selected active configured bounds | 5 of 20 | Exact active-bound count |

*Note:* Fits are ordered by the retained objective value. The decline is the initial minus terminal predicted mean chromosome number. The dead burden percentage is the maximum, over all daily 4N states, of the burden assigned by the model to death associated with resource stress as a percentage of total burden. It is therefore a trajectory maximum rather than an endpoint-only value. These optimizer-selected solutions are not biological replicates or a confidence interval.

Supplementary Table 7: Warm-start selection, joint numerical-search diagnostics, and objective eligibility for the 2 retained C families. Every family used the same separate *in vitro* seed144 anchor and 500 numerical starts. The primary lowest-objective decile retained 50 endpoints per family. Numerical endpoints and cluster assignments are search diagnostics rather than biological replicates, confidence sets, or biological groups.

| Family | Separate *in vivo* start | Joint winner | Minimum objective | Iterations of 1000 | DE flags of 500 | Local refinement | Winner bounds of 20 | Decile cutoff above minimum |
|---|---|---|---|---|---|---|---|---|
| C01 | 366 | 94 | 18.8810 | 26–95 | 500/500 | not accepted; code 52 | 2 | 0.063181 |
| C02 | 25 | 80 | 18.9350 | 26–45 | 500/500 | not accepted; code 52 | 1 | 0.002673 |

| Audit quantity | Value | Interpretation |
|---|---|---|
| Complete endpoints passing hard configuration and feasibility checks | 1,000 of 1,000 | 2 families, 500 endpoints per family |
| Widest retained objective set passing operator checks | 200 of 200 | Lowest-objective 20%, 100 endpoints per family |
| Primary complete context-specific response surfaces | 4 | Two contexts, 2 families, and 50 endpoints per family |
| Parameter-specific context-ratio records passing feasibility checks | 14,000 of 14,000 | 2 families, 500 endpoints, and 14 paired parameters |
| Pooled warm-start embedding | 256,000 vectors | 127,500 population points and 500 final solutions in each context |
| Saved primary clustering | $k = 2$; silhouette 0.673 | 2 primary C regions, with no secondary-cluster layer |
| Fixed-$k = 2$ 80% subsample agreement | Median ARI 1.000; 5th–95th percentile 1.000–1.000 | Agreement of resampled endpoint labels with the saved primary labels |
| Standardized 14-parameter-space $k = 2$ agreement | ARI 0.010 | Agreement with the saved t-SNE regions |

Supplementary Table 8: Context-specific differential-evolution initial-population and optimizer-endpoint distributions for the 14 paired parameters across the primary in-vivo clusters C01–C02. Each row reports the in vivo and in vitro natural-scale parameter distributions separately. The DE-initial distribution comprises 200,000 members reconstructed from the 500 runs (400 members per run) within a family; the endpoint distribution comprises the corresponding 500 feasible final solutions. Entries report median [5th, 95th percentile], followed by context-specific active-bound occupancy. Cross-family direction is an auxiliary sign-based audit of the paired endpoint ratios and does not apply the 0.8/1.2 equivalence band used for the family-specific classes in Figure 5D. These numerical-search distributions are not posterior or confidence intervals.

| Parameter | Family | In vivo: DE initial; endpoint | In vitro: DE initial; endpoint | Directional/search audit |
|---|---|---|---|---|
| `lam_max` | C01 | DE init 0.244 [0.206, 0.290]; endpoint 0.239 [0.239, 0.239]; bound=0% | DE init 1.33 [1.13, 1.48]; endpoint 1.34 [1.34, 1.34]; bound=0% | lower in vivo; max paired W_E/W_I=0; max bound=0% |
| `lam_max` | C02 | DE init 0.249 [0.211, 0.295]; endpoint 0.244 [0.244, 0.244]; bound=0% | DE init 1.33 [1.13, 1.48]; endpoint 1.34 [1.34, 1.34]; bound=0% | lower in vivo; max paired W_E/W_I=0; max bound=0% |
| `p_mis_base` | C01 | DE init 0.0000104 [0.00000163, 0.0000712]; endpoint 0.00000882 [0.00000503, 0.00000921]; bound=0% | DE init 0.00000292 [0.00000102, 0.0000200]; endpoint 0.00000248 [0.00000148, 0.00000361]; bound=0% | higher in vivo; max paired W_E/W_I=0.510; max bound=0% |
| `p_mis_base` | C02 | DE init 0.000504 [0.0000350, 0.00289]; endpoint 0.000506 [0.000506, 0.000506]; bound=0% | DE init 0.00000248 [0.00000107, 0.0000142]; endpoint 0.00000248 [0.00000248, 0.00000248]; bound=0% | higher in vivo; max paired W_E/W_I=0.510; max bound=0% |
| `p_wgd` | C01 | DE init 0.000174 [0.0000456, 0.000560]; endpoint 0.000182 [0.000182, 0.000202]; bound=0% | DE init 0.000323 [0.0000849, 0.000962]; endpoint 0.000339 [0.000304, 0.000339]; bound=0% | lower in vivo; max paired W_E/W_I=0.860; max bound=0% |
| `p_wgd` | C02 | DE init 0.0000136 [0.0000102, 0.000111]; endpoint 0.0000124 [0.0000124, 0.0000124]; bound=0% | DE init 0.000334 [0.0000229, 0.000941]; endpoint 0.000339 [0.000339, 0.000339]; bound=0% | lower in vivo; max paired W_E/W_I=0.860; max bound=0% |
| `p_misseg` | C01 | DE init 0.118 [0.0645, 0.212]; endpoint 0.118 [0.118, 0.118]; bound=0% | DE init 0.0105 [0.00593, 0.0188]; endpoint 0.0105 [0.0105, 0.0105]; bound=0% | higher in vivo; max paired W_E/W_I=0; max bound=0% |
| `p_misseg` | C02 | DE init 0.164 [0.0911, 0.290]; endpoint 0.164 [0.164, 0.164]; bound=0% | DE init 0.0105 [0.00601, 0.0186]; endpoint 0.0105 [0.0105, 0.0105]; bound=0% | higher in vivo; max paired W_E/W_I=0; max bound=0% |
| `k_o_mis` | C01 | DE init 0.00103 [0.000266, 0.00394]; endpoint 0.00103 [0.00103, 0.00103]; bound=0% | DE init 0.0000870 [0.0000225, 0.000334]; endpoint 0.0000870 [0.0000870, 0.0000870]; bound=0% | higher in vivo; max paired W_E/W_I=0; max bound=0% |

| Parameter | Family | In vivo: DE initial; endpoint | In vitro: DE initial; endpoint | Directional/search audit |
|---|---|---|---|---|
| `k_o_mis` | C02 | DE init 0.00306 [0.000852, 0.0109]; endpoint 0.00306 [0.00306, 0.00306]; bound=0% | DE init 0.0000870 [0.0000242, 0.000311]; endpoint 0.0000870 [0.0000870, 0.0000870]; bound=0% | higher in vivo; max paired W_E/W_I=0; max bound=0% |
| `O2_crit` | C01 | DE init 0.223 [0.186, 0.268]; endpoint 0.223 [0.223, 0.223]; bound=0% | DE init 1.57 [1.31, 1.89]; endpoint 1.57 [1.57, 1.57]; bound=0% | lower in vivo; max paired W_E/W_I=0; max bound=0% |
| `O2_crit` | C02 | DE init 0.148 [0.118, 0.186]; endpoint 0.148 [0.148, 0.148]; bound=0% | DE init 1.57 [1.25, 1.98]; endpoint 1.57 [1.57, 1.57]; bound=0% | lower in vivo; max paired W_E/W_I=0; max bound=0% |
| `n_O` | C01 | DE init 2.17 [1.83, 2.51]; endpoint 2.17 [2.17, 2.17]; bound=0% | DE init 1.97 [1.63, 2.31]; endpoint 1.97 [1.97, 1.97]; bound=0% | family-dependent or overlaps equality; max paired W_E/W_I=0; max bound=0% |
| `n_O` | C02 | DE init 1.65 [1.35, 1.95]; endpoint 1.65 [1.65, 1.65]; bound=0% | DE init 1.97 [1.67, 2.27]; endpoint 1.97 [1.97, 1.97]; bound=0% | family-dependent or overlaps equality; max paired W_E/W_I=0; max bound=0% |
| `alpha_o2` | C01 | DE init 2.53 [2.26, 2.73]; endpoint 2.69 [2.69, 2.69]; bound=0% | DE init 0.530 [0.503, 0.592]; endpoint 0.500 [0.500, 0.500]; bound=99.6% | higher in vivo; max paired W_E/W_I=0; max bound=100% |
| `alpha_o2` | C02 | DE init 1.69 [1.56, 1.77]; endpoint 1.77 [1.77, 1.77]; bound=0% | DE init 0.522 [0.502, 0.567]; endpoint 0.500 [0.500, 0.500]; bound=100% | higher in vivo; max paired W_E/W_I=0; max bound=100% |
| `gamma_growth` | C01 | DE init 3.23 [2.44, 3.73]; endpoint 3.31 [3.31, 3.31]; bound=0% | DE init 1.60 [1.51, 1.92]; endpoint 1.50 [1.50, 1.50]; bound=100% | higher in vivo; max paired W_E/W_I=0; max bound=100% |
| `gamma_growth` | C02 | DE init 2.80 [2.09, 3.24]; endpoint 2.86 [2.86, 2.86]; bound=0% | DE init 1.58 [1.51, 1.88]; endpoint 1.50 [1.50, 1.50]; bound=100% | higher in vivo; max paired W_E/W_I=0; max bound=100% |
| `mu_hp` | C01 | DE init 0.0467 [0.0123, 0.104]; endpoint 0.0515 [0.0515, 0.0515]; bound=0% | DE init 0.00574 [0.00506, 0.0101]; endpoint 0.00500 [0.00500, 0.00500]; bound=99.8% | higher in vivo; max paired W_E/W_I=0; max bound=100% |
| `mu_hp` | C02 | DE init 0.0397 [0.0104, 0.0885]; endpoint 0.0434 [0.0434, 0.0434]; bound=0% | DE init 0.00571 [0.00505, 0.0102]; endpoint 0.00500 [0.00500, 0.00500]; bound=100% | higher in vivo; max paired W_E/W_I=0; max bound=100% |
| `gamma_mu` | C01 | DE init 2.57 [1.95, 2.97]; endpoint 2.64 [2.64, 2.64]; bound=0% | DE init 1.28 [1.21, 1.54]; endpoint 1.20 [1.20, 1.21]; bound=94% | higher in vivo; max paired W_E/W_I=0.0169; max bound=98.4% |

| Parameter | Family | In vivo: DE initial; endpoint | In vitro: DE initial; endpoint | Directional/search audit |
|---|---|---|---|---|
| `gamma_mu` | C02 | DE init 2.59 [1.96, 2.99]; endpoint 2.66 [2.66, 2.66]; bound=0% | DE init 1.28 [1.21, 1.54]; endpoint 1.20 [1.20, 1.20]; bound=98.4% | higher in vivo; max paired W_E/W_I=0.0169; max bound=98.4% |
| `buffer_smax` | C01 | DE init 0.871 [0.735, 0.986]; endpoint 0.909 [0.909, 0.909]; bound=0% | DE init 0.961 [0.826, 0.999]; endpoint 1.00 [1.00, 1.00]; bound=99.6% | lower in vivo; max paired W_E/W_I=0; max bound=99.8% |
| `buffer_smax` | C02 | DE init 0.864 [0.728, 0.985]; endpoint 0.900 [0.900, 0.900]; bound=0% | DE init 0.964 [0.828, 0.999]; endpoint 1.00 [1.00, 1.00]; bound=99.8% | lower in vivo; max paired W_E/W_I=0; max bound=99.8% |
| `buffer_beta` | C01 | DE init 0.165 [0.133, 0.205]; endpoint 0.165 [0.165, 0.165]; bound=0% | DE init 1.60 [1.29, 1.98]; endpoint 1.60 [1.60, 1.60]; bound=0% | lower in vivo; max paired W_E/W_I=0; max bound=0% |
| `buffer_beta` | C02 | DE init 0.310 [0.267, 0.358]; endpoint 0.310 [0.310, 0.310]; bound=0% | DE init 1.60 [1.38, 1.85]; endpoint 1.60 [1.60, 1.60]; bound=0% | lower in vivo; max paired W_E/W_I=0; max bound=0% |
| `buffer_n_exp` | C01 | DE init 9.37 [7.41, 9.94]; endpoint 10.0 [10.0, 10.0]; bound=99.6% | DE init 3.32 [2.34, 5.84]; endpoint 3.16 [3.16, 3.16]; bound=0% | higher in vivo; max paired W_E/W_I=0; max bound=99.6% |
| `buffer_n_exp` | C02 | DE init 9.37 [7.41, 9.94]; endpoint 10.0 [10.0, 10.0]; bound=0% | DE init 3.32 [2.34, 5.84]; endpoint 3.16 [3.16, 3.16]; bound=0% | higher in vivo; max paired W_E/W_I=0; max bound=99.6% |

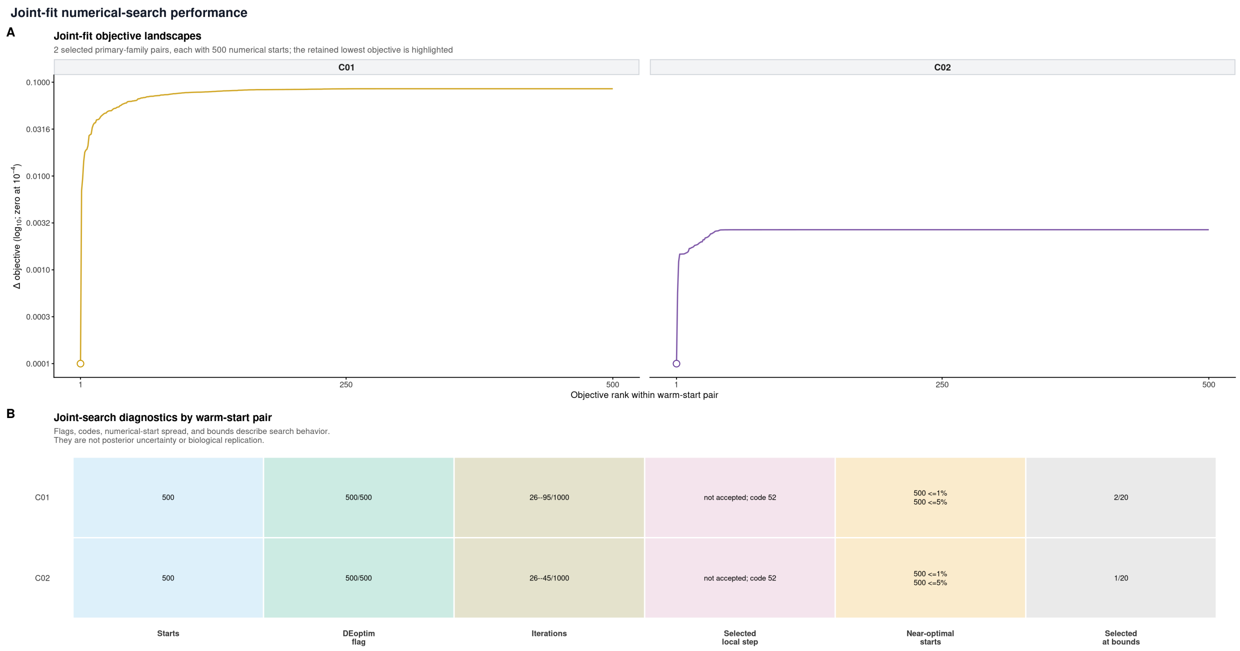


Supplementary Figure 3: **Joint-fit numerical-search performance across the two retained C-family ensembles. (A)** Objective-difference landscapes for the C01 and C02 500-start joint fits. Within each family, starts are ranked by objective, the lowest retained winner is highlighted, and exact zero is displayed at $10^{-4}$ on the $\log_{10}$ axis. **(B)** C-family-resolved run diagnostics, including numbers of starts, recorded differential-evolution convergence flags, completed-iteration ranges relative to the target, selected local-step status and code, near-optimal start counts, and exact active-bound counts for the selected winners. The corresponding C-family rows are reported in Supplementary Table 7. These diagnostics characterize the sampled numerical searches and optimizer stability across starts.

Supplementary Table 9: Current Figure 6 analysis design. The scanned input is the mechanistic $p_{\text{misseg}}$ model parameter while the other parameters retain their fitted context-specific values; state-specific effective missegregation remains a model-derived quantity.

| Component | Current specification |
|---|---|
| Displayed families | C01 and C02 |
| Primary ensemble | Lowest-objective q10; 50 optimizer endpoints per family |
| Exact endpoint multiplicity | C01: 50 unique/50 seeds, max multiplicity 1; C02: 16 unique/50 seeds, max multiplicity 35 |
| Steady-state mechanistic scan | 201 oxygen values from 0% to 5%; 60 log-spaced fixed mechanistic p_misseg values from 0.005 to 0.5 |
| Main oxygen display window | 0–2% oxygen in both steady-state and finite-time panels |
| Main finite-time display | Panels A-B distinguish steady state from finite time; displayed finite-time window is 0-1000 days |
| Full-range validation source | 0-10000 days; 5 initial-ploidy states, 201 oxygen values and 5 fixed p_misseg values per validated panel |
| Propagation modes | Continuous in vivo; continuous comparator and threshold-triggered stochastic passage in vitro |
| Interpretive status | Post-fit counterfactual/model simulation; optimizer endpoints are numerical solutions, not biological replicates or posterior samples |

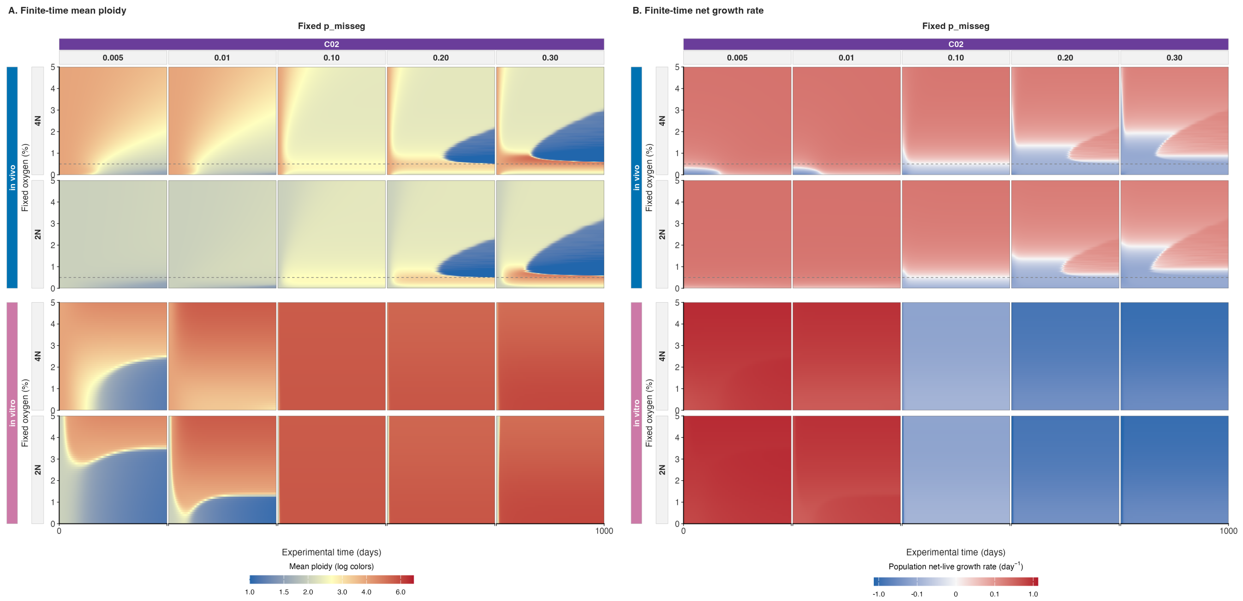


Supplementary Figure 4: **C02 finite-time mean ploidy and population net-live growth under fixed oxygen.** The C02 joint-fit family is evaluated from 4N and 2N initial populations (upper and lower rows within each context) over 0–1,000 days at fixed $p_{\text{misseg}} = 0.005$, 0.01, 0.10, 0.20, or 0.30 and 0–5% $O_2$. **(A)** Mean ploidy (autosomal chromosome number/22, logarithmic color spacing). **(B)** Population net-live growth rate (day$^{-1}$) on a signed pseudo-log$_{10}$ color scale with a 0.01 day$^{-1}$ linear threshold (red, positive; blue, negative; white, zero). Values are arithmetic means across the same 50 lowest-objective C02 optimizer endpoints used in Figure 6; stochastic culture repeats are averaged within endpoints first. Tumors grow continuously; culture uses daily threshold-triggered stochastic passage, with passage sampling and dilution excluded from the plotted growth rate. Dashed horizontal lines mark 0.5% $O_2$. The ploidy and growth maps describe the same model conditions; negative growth in high-ploidy regions denotes declining viable populations.

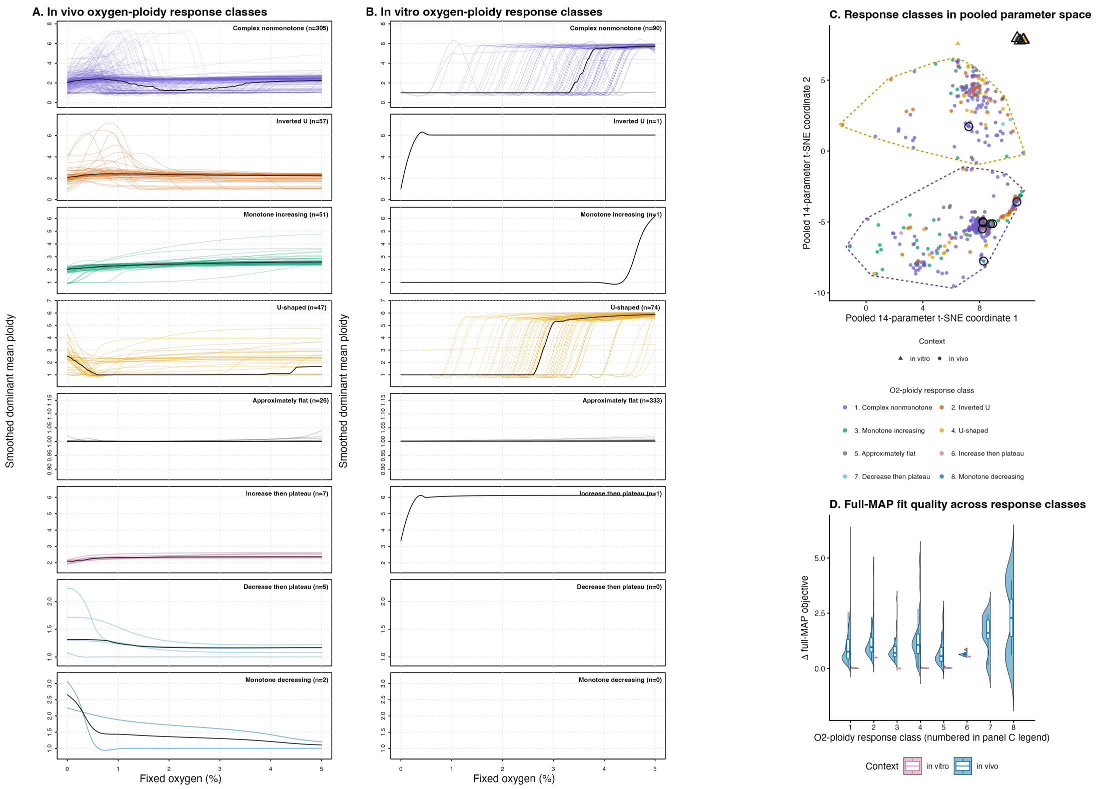


Supplementary Figure 5: **Separate-fit fixed-$O_2$ response classes and their fitted-landscape context. (A–B) Context-specific oxygen–ploidy response classes.** Analytical dominant-eigenvector ploidy curves from the 500 separate *in vivo* fits (A) and 500 separate *in vitro* fits (B) were grouped by the same regression-smoothed response-shape rule. Both columns use the same fixed eight-class order; an empty plotting region therefore records a class absent from that context rather than omitted data. Class names and seed counts are printed in the upper-right corner. Thin colored curves show individual optimizer-seed endpoints and the black curve is the pointwise class median. Panel-specific ploidy ranges preserve visibility of both broad and nearly flat responses. These class counts include reliable, caution, and unreliable spectral-gap strata. **(C) Response classes in pooled parameter space.** All 1,000 separate-fit endpoints are positioned in the pooled 14-parameter t-SNE coordinate system used for the joint warm-start landscape in Figure 5C and colored by fixed-$O_2$ response class. Circles denote *in vivo* and triangles denote *in vitro*. These unitless visualization coordinates describe local numerical neighborhoods and are distinct from the *in vivo*-only 18-parameter embedding in Figure 4C. A black outline marks the lowest-objective endpoint within every nonempty context–class combination. Family-colored dashed outlines delimit the C01 and C02 *in vivo* warm-start search-design regions. **(D) Context-relative full-MAP fit quality across response classes.** For each context, the minimum full MAP objective across its 500 separate fits was subtracted before comparison. Blue left and pink right half-violins show the *in vivo* and *in vitro* distributions, respectively, with narrow context-colored boxplots showing medians and interquartile ranges. Empty halves identify classes absent from that context. Because the two fits use different observation models, the objective difference compares relative fit quality within each context. Together, panels C–D diagnose fitted-landscape multiplicity across response shapes.

Supplementary Table 10: Current Figure 6 robustness and numerical-validation summary. Optimizer endpoints are numerical solutions rather than biological replicates, posterior samples, or confidence intervals.

| Check | Observed | Interpretation |
|---|---|---|
| q20 operator checks | 200/200 | All q20 endpoint/operator combinations passed |
| Cutoff invariance of qualitative results | 16/16 | Modal qualitative result unchanged from q05 through q20 |
| Seed-weighted versus unique-endpoint modal agreement | 48/48 | Exact deduplication did not change any audited modal conclusion |
| Primary q10 exact endpoint counts | C01: 50/50; C02: 16/50 | Multiplicity is substantial for C02 and must be reported |
| Full-range panel contracts | 6/6 passed | All current context/mode/family full-range panels passed |
| Stochastic-passage implementation tests | 30/30 passed | Determinism, resume, RNG, allocation and external-reference checks passed |
| Weak-gap cells with at least 90% endpoint regime consensus in vivo C01 | 99.96% | Fraction of weak-gap cells with at least 90% endpoint agreement |
| Weak-gap cells with at least 90% endpoint regime consensus in vivo C02 | 99.94% | Fraction of weak-gap cells with at least 90% endpoint agreement |
| Weak-gap cells with at least 90% endpoint regime consensus in vitro C01 | 99.45% | Fraction of weak-gap cells with at least 90% endpoint agreement |
| Weak-gap cells with at least 90% endpoint regime consensus in vitro C02 | 99.86% | Fraction of weak-gap cells with at least 90% endpoint agreement |

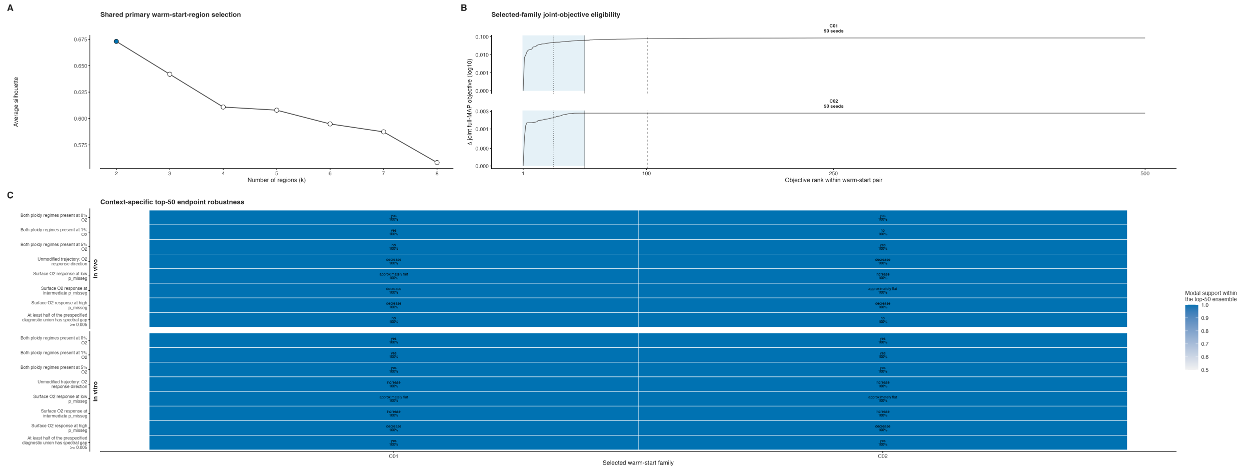


Supplementary Figure 6: **Cluster-selection, objective-eligibility, and multi-seed robustness diagnostics for Figure 6.** **(A)** Average silhouette over candidate primary-region counts in the frozen pooled 14-parameter t-SNE coordinates. The saved primary partition is $k = 2$, with average silhouette approximately 0.673; the displayed 80% subsample audit conditions on the saved embedding rather than re-estimating t-SNE. **(B)** Full joint MAP objective differences ordered across all 500 numerical endpoints within C01 and C02. The blue region is the primary lowest-objective decile; vertical lines mark the nested 5%, 10%, and 20% eligibility thresholds. Facet labels report the 50 retained optimizer seeds. **(C)** Qualitative oxygen–CIN–ploidy diagnostics for the primary 50-endpoint ensembles, evaluated separately from the *in vivo* and *in vitro* parameter vectors of C01 and C02. Text reports the modal result and its exact support. The corresponding design and robustness values are reported in Supplementary Tables 7, 9, and 10. This categorical audit summarizes ensemble-level numerical-search behavior; pointwise surface variation is evaluated in the dedicated robustness panels.

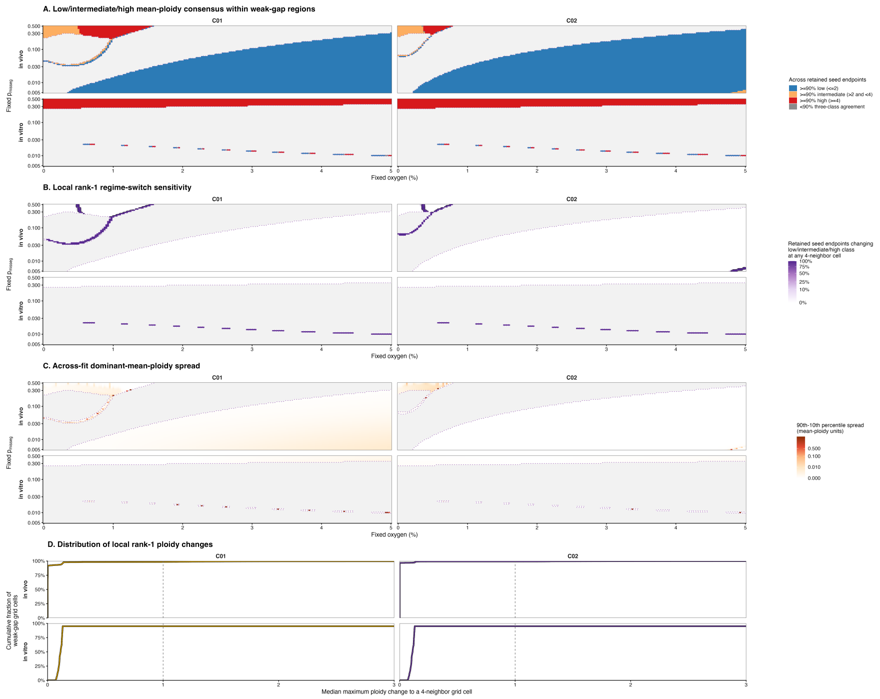


Supplementary Figure 7: **Joint-fit tumor robustness within weak spectral-gap regions.** This figure is assigned to the joint-fit *in vivo* analysis. All panels use the same 201 oxygen concentrations, 60 fixed mechanistic $p_{\text{misseg}}$ values, and family-specific lowest-objective 50-joint-endpoint ensembles as Figure 6A. Columns show C01 and C02. The upper rows show the *in vivo* results that constitute the primary analysis; the lower *in vitro* rows are retained only as paired display context, with the dedicated joint-fit culture analysis reported in Supplementary Figure 8. The dotted purple boundary encloses grid cells where at least half of the context-specific endpoint vectors have dominant spectral gap below 0.005; areas outside that weak-gap region are gray. **(A)** Across-endpoint regime consensus within the weak-gap region. Stable low, intermediate, and high require at least 90% of endpoints to place dominant mean ploidy at or below 2, between 2 and 4, or at or above 4, respectively; gray within the colored region would denote less than 90% three-class agreement. The corresponding *in vivo* consensus fractions are 99.96% for C01 and 99.94% for C02. **(B)** Proportion of endpoints for which the rank-1 mean-ploidy classification changes among low, intermediate, and high at any of the up to four immediately adjacent $O_2$–missegregation grid cells, shown with square-root color spacing and percentage labels in the original scale. The black contour marks a proportion of 0.5. **(C)** Across-endpoint 90th–10th percentile spread of rank-1 dominant mean ploidy, shown in original ploidy units with pseudo-logarithmic color spacing. **(D)** Empirical cumulative distribution of the endpoint-median maximum absolute rank-1 ploidy change to an immediately adjacent grid cell, restricted to weak-gap cells. The vertical dashed line marks one ploidy unit; annotations report the fraction with a three-class change in at least half of endpoints and the fraction with median local ploidy change of at least one unit. The ECDF is calculated from all weak-gap cells, while the displayed horizontal range is limited to 0–3 ploidy units; a curve can therefore remain below 100% at the right boundary when larger changes occur. Exact duplicate context-specific parameter endpoints were evaluated once and restored to their original optimizer-seed multiplicities before aggregation. Family- and context-level counts and robustness summaries are reported in Supplementary Table 10. These post-fit diagnostics quantify numerical robustness across endpoints.

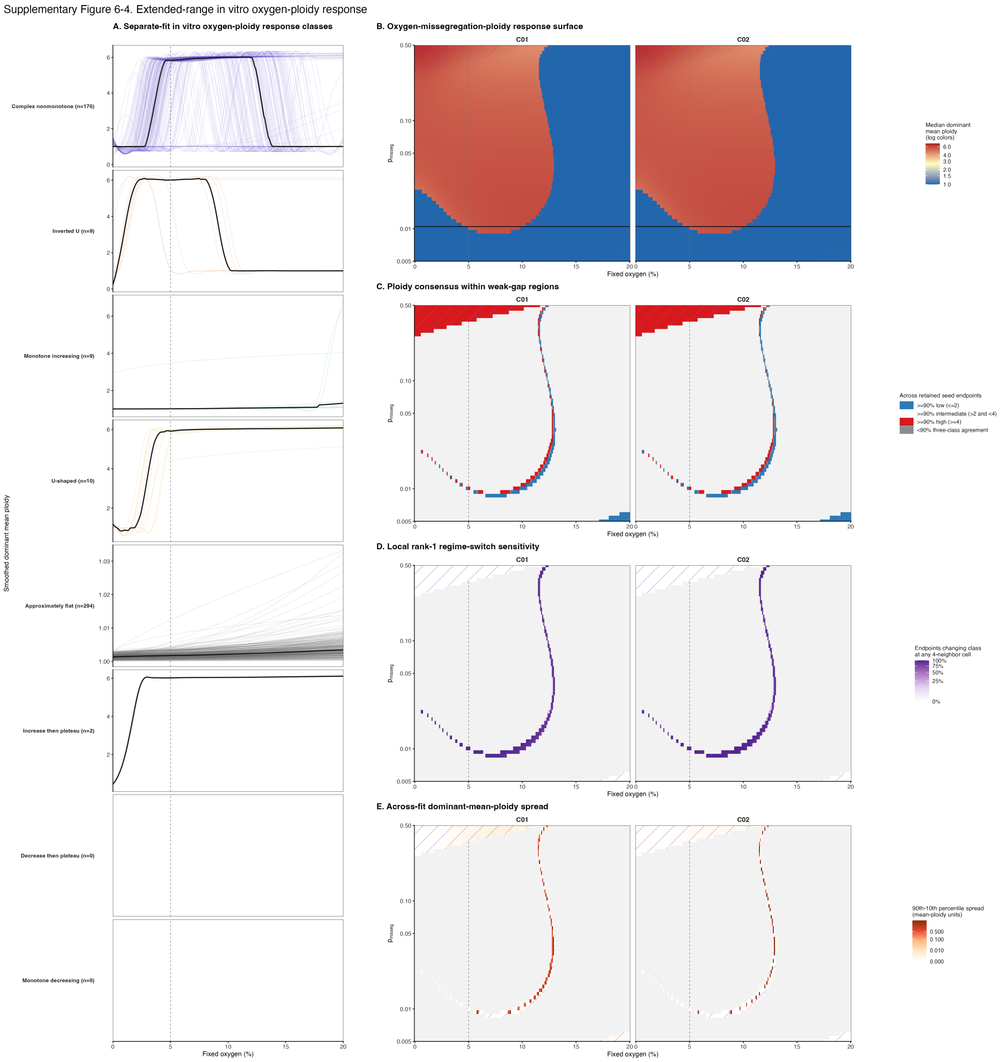


Supplementary Figure 8: **Joint-fit culture response over the extended oxygen domain.** This figure presents the joint-fit *in vitro* analysis. **(A)** Separate-fit culture response classes across 0–20% oxygen provide landscape context; pale curves are individual optimizer endpoints and the thick black curve is the pointwise class median when the class is nonempty. The vertical 5% line marks the upper edge of the primary fixed-oxygen analysis range, not the upper limit of the culture calibration data, which include a 20.5% $O_2$ control. **(B)** C01/C02 joint-fit culture steady-state surfaces over extended oxygen and fixed mechanistic $p_{\text{misseg}}$; fill is the seed-weighted median dominant mean ploidy, displayed with logarithmic color spacing. **(C)** Three-class endpoint consensus within weak-gap regions: low is mean ploidy at or below 2, intermediate is between 2 and 4, and high is at or above 4; gray denotes less than 90% agreement. **(D)** The proportion of endpoints whose rank-1 ploidy class changes at any immediately adjacent oxygen–$p_{\text{misseg}}$ grid cell. **(E)** The across-endpoint 90th–10th percentile spread of dominant mean ploidy. Purple hatching marks weak-gap regions. Values above 5% extend the primary analysis range; the constant-oxygen and altered-missegregation settings remain post-fit counterfactuals rather than additional observations. These panels report continuous steady-state diagnostics; Figure 6B reports passage-aware finite-time culture simulations.

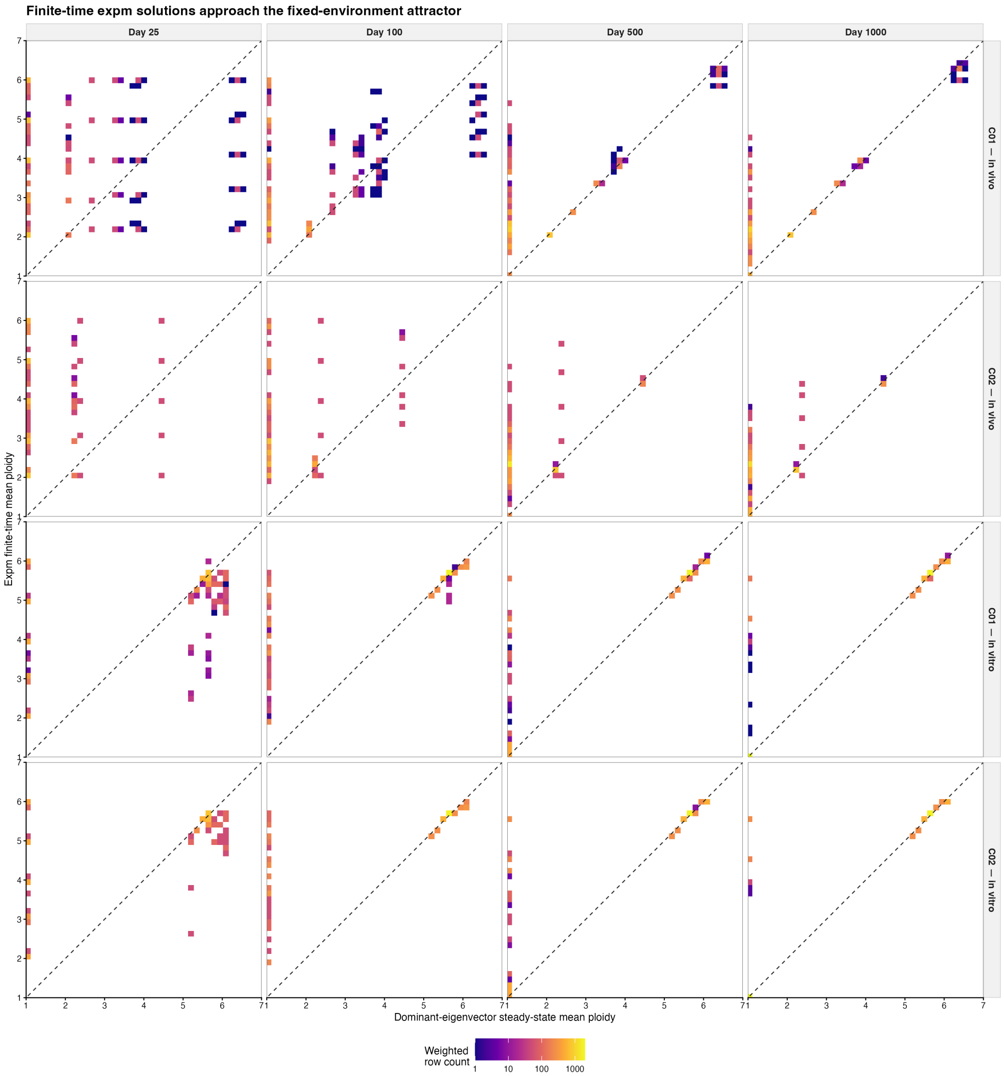


Supplementary Figure 9: **Finite-time response compared with the steady-state attractor.** Finite-time mean ploidy is compared with the corresponding dominant-eigenvector steady-state mean at days 25, 100, 500, and 1,000 for both contexts and both families. Bias and RMSE generally decline and points move toward the identity line with elapsed time; some culture conditions remain separated from the attractor at day 1,000. This numerical comparison quantifies convergence within the evaluated model conditions.

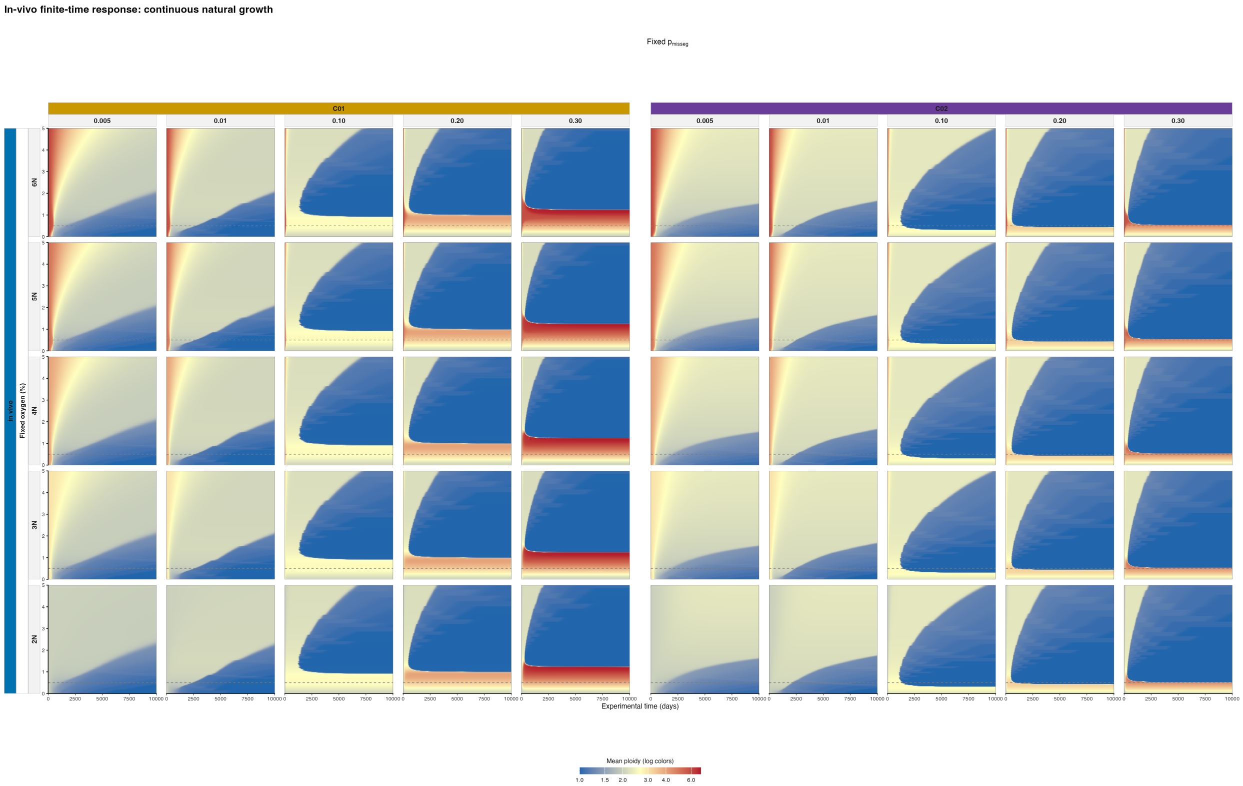


Supplementary Figure 10: **Long-horizon continuous tumor response.** Continuous natural-growth simulations span C01/C02, 2N–6N initial states, five fixed mechanistic $p_{\text{misseg}}$ values, 0–5% oxygen, and 0–10,000 days. This long-horizon model extrapolation compares initial-state persistence and asymptotic behavior under the tumor operator.

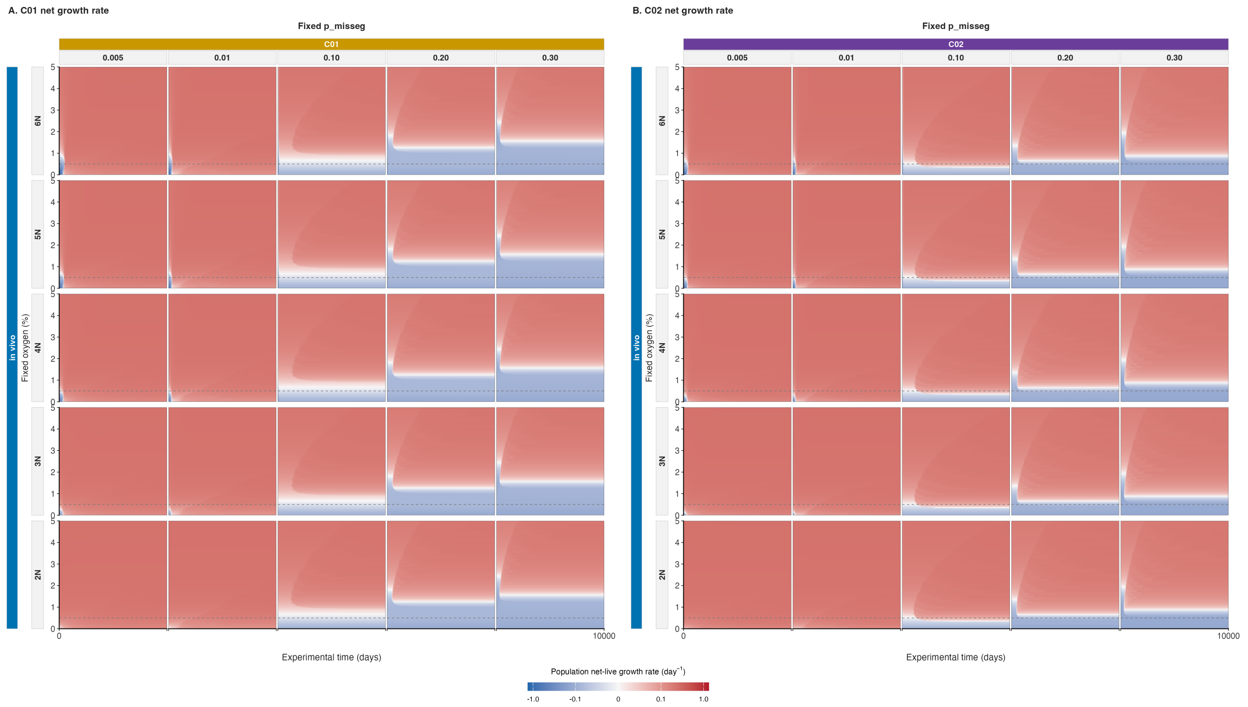


Supplementary Figure 11: **Long-horizon population net-live growth rate in tumors.** Panels **(A)** and **(B)** display C01 and C02, respectively. The continuous *in vivo* projections span 2N–6N initial states, fixed $p_{\mathrm{misseg}}$ values of 0.005, 0.01, 0.10, 0.20, and 0.30, 0–5% $O_2$, and 0–10,000 days. Fill gives the population-weighted net-live growth rate (day$^{-1}$), averaged across the same lowest-objective 50 endpoints per family used for Figure 6. A signed pseudo-log$_{10}$ color scale with a 0.01 day$^{-1}$ linear threshold shows positive growth in red, negative growth in blue, and zero in white. The dashed horizontal line marks 0.5% $O_2$. These maps pair with the long-horizon mean-ploidy projections in Supplementary Figure 10; negative values indicate declining viable populations under the modeled fixed conditions.

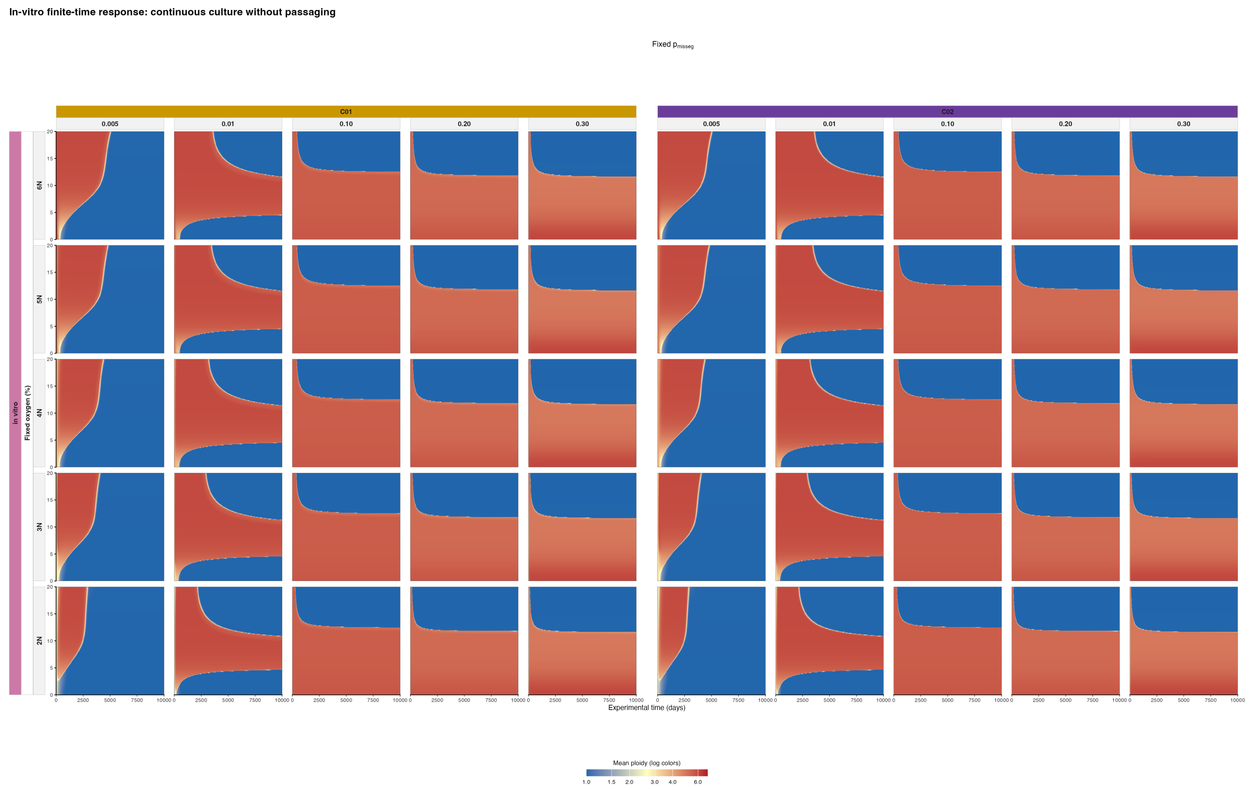


Supplementary Figure 12: **Long-horizon continuous culture response without passaging.** Continuous simulations span C01/C02, 2N–6N initial states, five fixed mechanistic $p_{\text{misseg}}$ values, 0–20% oxygen, and 0–10,000 days. Family patterns are similar, with oxygen-dependent low/high-ploidy transitions that vary with fixed input. This counterfactual isolates continuous propagation without threshold-triggered passage and extends the primary 0–5% analysis range; its prescribed conditions and long time horizon are distinct from the measured passage schedule.

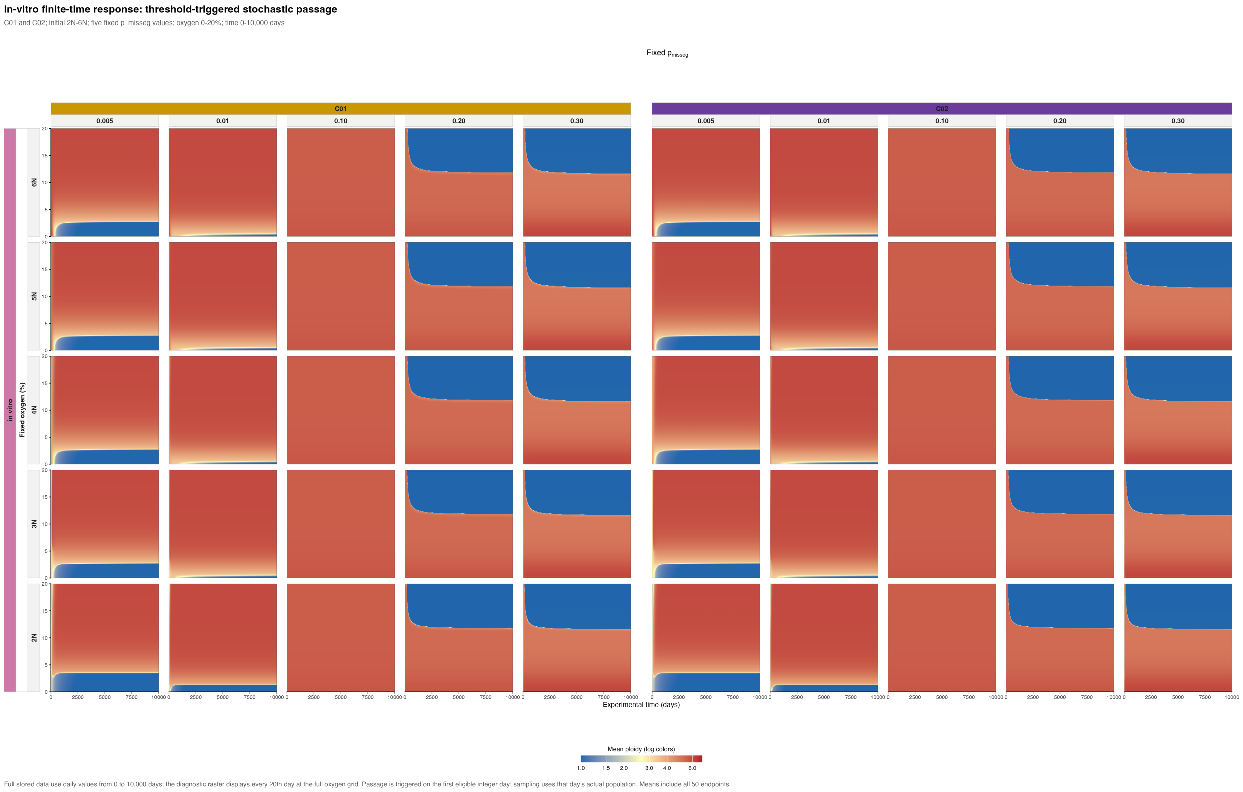


Supplementary Figure 13: **Long-horizon passage-aware culture response.** Threshold-triggered stochastic-passage simulations span C01/C02, 2N–6N initial states, five fixed mechanistic $p_{\text{misseg}}$ values, 0–20% oxygen, and 0–10,000 days. Passage occurs on the first eligible integer day and samples the actual population; curves are averaged first over stochastic repeats and then over optimizer endpoints. Initial-state effects are most visible early, and high-oxygen low-ploidy regions persist for selected inputs. These counterfactual model simulations extend beyond the measured time range and the primary 0–5% analysis range. The culture calibration includes a 20.5% control but does not independently validate the intervening constant-oxygen response surface.

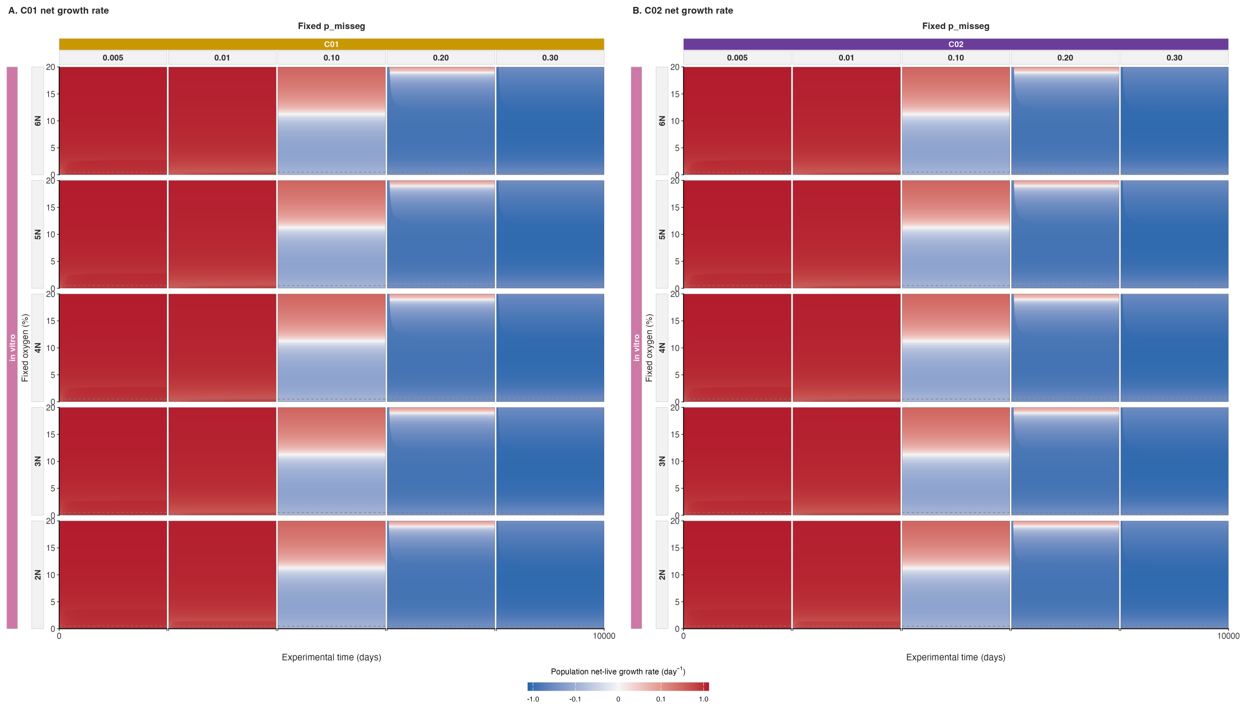


Supplementary Figure 14: **Long-horizon population net-live growth rate in passage-aware culture.** Panels **(A)** and **(B)** display C01 and C02, respectively. The *in vitro* projections span 2N–6N initial states, fixed $p_{\mathrm{misseg}}$ values of 0.005, 0.01, 0.10, 0.20, and 0.30, 0–20% $O_2$, and 0–10,000 days. Fill gives the population-weighted net-live growth rate (day$^{-1}$), averaged first over stochastic passage repeats and then over the same lowest-objective 50 optimizer endpoints per family used for Figure 6. A signed pseudo-$\log_{10}$ color scale with a 0.01 day$^{-1}$ linear threshold shows positive growth in red, negative growth in blue, and zero in white. Daily threshold-triggered stochastic passage is applied to the trajectories, but sampling and dilution are excluded from the plotted growth rate. The dashed horizontal line marks 0.5% $O_2$. These maps pair with the long-horizon passage-aware mean-ploidy projections in Supplementary Figure 13. The prescribed fixed-oxygen and missegregation settings extend beyond the measured passage schedule.

#### 1.3 A Mechanistic Model of Chromosome-Number Evolution

We model tumor population dynamics as a deterministic state-structured ODE system over chromosome-number classes. The formulation separates viable cells from non-viable biomass, couples mean-field effective oxygen to resource-dependent growth and stress-associated death, and uses chromosome missegregation and whole-genome doubling to move viable mass across chromosome-number states.

**State space and compartments.** We consider tumor cells stratified by modeled total autosomal chromosome count; sex chromosomes are not included in this state variable. Let

$$\mathcal{N} = \{N_{\min}, \ldots, N_{\max}\}. \tag{3}$$

This set denotes the allowed autosomal chromosome-count grid. For the simulations described here, $N_{\min} = 22$ and $N_{\max} = 154$. Let $N_{\text{dip}} = 44$ denote the diploid autosomal chromosome-count reference state, and let $N_{\text{unit}} = 22$ denote the one-ploidy autosomal chromosome-count unit, so that $2N_{\text{unit}} = N_{\text{dip}}$.

The viable abundance vector is

$$\mathbf{x}^L(t) = \{x_N^L(t)\}_{N\in\mathcal{N}}, \tag{4}$$

where $x_N^L(t)$ is the number of viable cells with modeled autosomal chromosome count $N$ at time $t$. The ODE state is indexed only by autosomal chromosome count and by live/dead compartment. Unless explicitly marked as a dead compartment, chromosome-number distributions in the model are defined over the viable population. The total viable population size is

$$N_{\text{tot}}(t) = \sum_{N\in\mathcal{N}} x_N^L(t). \tag{5}$$

To represent non-viable biomass contributing to macroscopic tumor volume, we track two dead pools:

$$\mathbf{x}^{D,\text{hyp}}(t) = \{x_N^{D,\text{hyp}}(t)\}_{N\in\mathcal{N}}, \qquad \mathbf{x}^{D,\text{CIN}}(t) = \{x_N^{D,\text{CIN}}(t)\}_{N\in\mathcal{N}}. \tag{6}$$

Here $\mathbf{x}^{D,\text{hyp}}$ receives cells assigned to the modeled pathway for death associated with resource stress, and $\mathbf{x}^{D,\text{CIN}}$ receives non-viable daughter mass generated by chromosome missegregation or WGD; the superscript CIN denotes missegregation-origin loss or out of grid WGD. The superscript hyp is an internal label for a model assigned compartment. It does not indicate that the biological cause of death was directly observed or uniquely identified. Their sum is

$$\mathbf{x}^D(t) = \mathbf{x}^{D,\text{hyp}}(t) + \mathbf{x}^{D,\text{CIN}}(t). \tag{7}$$

The model's bulk tumor burden is therefore

$$N_{\text{burden}}(t) = \sum_{N\in\mathcal{N}} \left( x_N^L(t) + x_N^{D,\text{hyp}}(t) + x_N^{D,\text{CIN}}(t) \right), \tag{8}$$

which includes viable cells and dead biomass. This distinction is used throughout: tumor-volume observations are compared with total burden, whereas single-cell chromosome-number observations sample the viable distribution.

**Environmental stress functions.** Let $O_2(t)$ denote the mean-field approximation of the effective oxygen level experienced by the tumor population. Oxygen-dependent resource stress is represented by the bounded Hill function

$$h(O_2) = \frac{O_{2,c}^{n_O}}{O_{2,c}^{n_O} + O_2^{n_O}}, \tag{9}$$

where $O_{2,c}$ is the critical oxygen level at which the stress term is half-maximal, and $n_O$ controls the steepness of the transition from low stress at high oxygen to strong stress under oxygen limitation. A Hill-type stress function is biologically justified because the literature indicates that oxygen-dependent cell fate is graded but threshold-like rather than linear: "The severity of hypoxia determines whether cells become apoptotic or adapt to hypoxia and survive"[36], and recent spheroid work identifies a "critical survival $pO_2$"[37]; accordingly, Eq. (9) provides a smooth, bounded approximation to an oxygen-linked resource-stress threshold.

**Proliferation and stress-associated death rates.** Given chromosome count $N$ and effective oxygen level $O_2$, the effective division rate is

$$\lambda_{\text{eff}}(N, O_2) = \lambda_{\max} \frac{1}{1 + \alpha_{O_2}\, h(O_2) \left(\frac{N}{N_{\text{dip}}}\right)^{\gamma_{\text{growth}}}}. \tag{10}$$

Here $\alpha_{O_2}$ controls the strength of resource-stress-dependent growth damping, and $\gamma_{\text{growth}}$ controls how strongly this damping increases with chromosome content. When oxygen-linked stress is small, $h(O_2) \approx 0$ and the reciprocal factor approaches one; as oxygen decreases, the reciprocal factor suppresses proliferation more strongly, especially for high-chromosome-count states.

Stress-associated death is applied as an explicit live-to-dead flow. The per-capita death hazard is

$$\mu_{\text{eff}}(N, O_2) = \mu_{\text{hp}}\, h(O_2) \left(\frac{N}{N_{\text{dip}}}\right)^{\gamma_\mu}. \tag{11}$$

Thus $\mu_{\text{hp}}$ sets the stress-associated death scale, $h(O_2)$ makes death depend on the same oxygen-linked stress function (Eq. (9)), and $\gamma_\mu$ controls the chromosome-number penalty on stress-associated death.

**Cell division, WGD, and chromosomal transitions.** With the stress-modulated rates defined, divisions are represented by a live-state generator matrix

$$G_{\text{div}}(O_2) \in \mathbb{R}^{R\times R}, \qquad R = N_{\max} - N_{\min} + 1. \tag{12}$$

Let

$$L(O_2) = \operatorname{diag}\big(\lambda_{\text{eff}}(N, O_2)\big) \tag{13}$$

be the diagonal matrix of state-specific division rates. A division follows the ordinary chromosome-missegregation branch with probability $1 - p_{\text{wgd}}$ and follows a whole-genome-doubling branch with probability $p_{\text{wgd}}$. The corresponding division generator is

$$G_{\text{div}}(O_2) = (1 - p_{\text{wgd}})\, B(O_2)L(O_2) + p_{\text{wgd}}\, W\, L(O_2) - L(O_2), \tag{14}$$

where $B(O_2)$ is the viable offspring-allocation matrix generated by chromosome missegregation and post-missegregation viability filtering, $W$ maps a mother state $N$ to the within-grid doubled state $2N$, and the final $-L(O_2)$ term removes the dividing mother cell. The ordinary branch $B$ accounts for the two daughter branches of a division and therefore has total mass up to two before viability losses. By contrast, the WGD branch is implemented as an effective tracked-lineage conversion: $W$ has unit mass in the doubled state, so a WGD event replaces the mother lineage by one viable doubled lineage when $2N$ lies in the modeled grid. Out-of-grid WGD mass is not retained in the viable state space and is routed to the CIN-associated dead compartment.

**Chromosome missegregation and daughter viability.** For a mother cell with chromosome count $N$, the per-chromosome missegregation probability increases with the stress-associated death hazard defined in Eq. (11):

$$p_{\text{mis}}(N, O_2) = p_{\text{mis},base} + p_{\text{misseg}} \frac{\mu_{\text{eff}}(N, O_2)}{\mu_{\text{eff}}(N, O_2) + k_{o,\text{mis}}}, \tag{15}$$

with values clipped to $[0, 1]$. Here $p_{\text{mis},base}$ is the baseline per-chromosome missegregation probability under low stress, $p_{\text{misseg}}$ is the maximal stress-induced increment above baseline, and $k_{o,\text{mis}}$ is the death-hazard scale at which the stress-induced increase in missegregation is half-maximal.

Let $m$ denote the number of missegregated chromosome copies in a division of a mother cell with chromosome count $N$. We model

$$m \sim \text{Binomial}\{N, p_{\text{mis}}(N, O_2)\}. \tag{16}$$

For each value of $m$, the two daughter branches are assigned coarse chromosome-count shifts $+m$ and $-m$. Before applying the daughter-viability modifier, the shift kernel is

$$d_N(\Delta; O_2) = \sum_{m=0}^{N} \binom{N}{m} \{p_{\text{mis}}(N, O_2)\}^m \{1 - p_{\text{mis}}(N, O_2)\}^{N-m} [\mathbf{1}\{\Delta = m\} + \mathbf{1}\{\Delta = -m\}]. \tag{17}$$

This kernel sums over two daughter branches, so its total mass is two before viability filtering.

For a daughter with chromosome-count shift $\Delta$, post-missegregation survival is approximated by the symmetric ploidy-dependent modifier

$$S_N(\Delta) = s_N^{|\Delta|}, \qquad s_N = s_{\max} \exp\left[-\beta_{\text{buf}} \left(\frac{N_{\text{dip}}}{N}\right)^{n_{\text{exp}}}\right]. \tag{18}$$

The parameter $s_{\max}$ is the maximum per-copy survival factor, $\beta_{\text{buf}}$ controls the severity of post-missegregation viability loss at lower ploidy, and $n_{\text{exp}}$ controls the steepness of the ploidy dependence. This form lets higher-chromosome-count states tolerate a given chromosome-count shift more readily than lower-chromosome-count states. This survival modifier enters the retained-grid offspring allocation and dead-pool accounting described in the boundary-handling subsection below.

**Boundary handling on the chromosome-count domain.** The chromosome-count state space is finite. We define the viable chromosome-count domain as

$$\mathcal{N} = \{N_{\min}, N_{\min} + 1, \ldots, N_{\max}\},$$

where $N_{\min} = 22$ and $N_{\max} = 154$ in the simulations described here. The viable state vector

$$\mathbf{x}^L(t) = \{x_N^L(t)\}_{N \in \mathcal{N}}$$

therefore tracks only cells whose modeled autosomal chromosome count lies within this retained grid.

This finite state space requires an explicit rule for division events that would produce daughter or doubled-lineage states outside $\mathcal{N}$. We use absorbing boundary handling on the viable chromosome-count grid. That is, daughter cells or WGD-derived lineages whose chromosome count would fall below $N_{\min}$ or exceed $N_{\max}$ are removed from the viable offspring allocation and routed to the CIN-associated dead compartment.

For ordinary missegregating divisions, the viable offspring-allocation matrix is defined only over retained daughter states $N' \in \mathcal{N}$:

$$B_{N',N}(O_2) = d_N(N' - N; O_2) S_N(N' - N), \qquad N, N' \in \mathcal{N}.$$

Here $d_N(\Delta; O_2)$ specifies the expected daughter allocation at chromosome-count shift $\Delta$ before viability filtering, and $S_N(\Delta)$ specifies the probability that a daughter with that shift remains viable. Any daughter allocation corresponding to $N + \Delta \notin \mathcal{N}$ is excluded from $B_{\cdot,N}$. Similarly, for whole-genome-doubling events, the doubling operator $W$ retains only within-grid transitions $N \mapsto 2N \in \mathcal{N}$; out-of-grid WGD offspring mass is excluded from $W_{\cdot,N}$ and routed to the CIN-associated dead compartment.

Because viability filtering or finite-grid exclusion can reduce the ordinary-division allocation below two daughters and the WGD allocation below one doubled lineage, we define the CIN-associated event-level nonviable inflow as

$$\phi_{\mathrm{CIN},N}(\mathbf{x}^L, O_2) = \lambda_{\mathrm{eff}}(N, O_2) x_N^L \left\{ (1 - p_{\mathrm{wgd}}) \left[ 2 - \sum_{N' \in \mathcal{N}} B_{N',N}(O_2) \right] + p_{\mathrm{wgd}} \left[ 1 - \sum_{N' \in \mathcal{N}} W_{N',N} \right] \right\}. \tag{19}$$

The first bracket is the ordinary-division deficit between two potential daughters and the viable daughter mass retained within $\mathcal{N}$. The second bracket accounts for WGD events for which the doubled state $2N$ lies outside the modeled grid. This term is distinct from the modeled continuous death hazard associated with resource stress, $\mu_{\mathrm{eff}}(N, O_2)$; it is added to the CIN-associated dead compartment $x^{D,\mathrm{CIN}}$.

Thus, the chromosome-count boundaries are absorbing for viable offspring mass, with out-of-domain and non-buffered daughter mass transferred to the dead compartment.

**Oxygen demand and supply dynamics.** To couple tumor size to oxygen demand, we define the chromosome-number-weighted oxygen-demand proxy

$$N_{\mathrm{eff}}(t) = \sum_N x_N^L(t) \left(\frac{N}{N_{\mathrm{dip}}}\right)^{\eta}, \tag{20}$$

where $\eta$ controls how oxygen demand scales with chromosome content. The reference scale $N_{\mathrm{ref}}$ is a fixed cell-count scale used to nondimensionalize demand, so that $N_{\mathrm{eff}}/N_{\mathrm{ref}}$ is a relative effective tumor size.

The oxygen supply target is the phenomenological supply–demand relation

$$O_2^*(N_{\mathrm{eff}}) = \max\left(O_{2,\min},\, S_0 - \kappa_O \log\left(1 + \frac{N_{\mathrm{eff}}}{N_{\mathrm{ref}}}\right)\right), \tag{21}$$

where $S_0$ is the low-burden oxygen supply level, $O_{2,\min}$ is the oxygen floor, and $\kappa_O$ controls the oxygen-drop amplitude per unit log-burden increase. The logarithmic term is dimensionless because of the normalization by $N_{\mathrm{ref}}$. Oxygen values are interpreted as percentages and are restricted to the physical interval $[0, 100]$.

Effective oxygen is updated by first-order relaxation toward this target:

$$\alpha_\tau = 1 - \exp\left(-\frac{\Delta t}{\tau_{O_2}}\right), \qquad O_{2,k+1} = O_{2,k} + \alpha_\tau (O_2^*(N_{\mathrm{eff},k}) - O_{2,k})\,. \tag{22}$$

Thus oxygen changes gradually toward the demand-adjusted target rather than instantaneously matching it at each simulation step.

**Discrete-time simulation scheme.** The simulations use a time step of $\Delta t = 0.05$ days. At each step $k \to k+1$, the oxygen state is first relaxed toward the supply–demand target in Eq. (22), using the current viable population to calculate $N_{\mathrm{eff},k}$. The resulting relaxed value, $O_{2,k+1}$, is then used to evaluate division-linked chromosome transitions and explicit stress-associated death for that population step:

$$\mathbf{x}_{k+1}^L = \mathbf{x}_k^L + \Delta t\, G_{\mathrm{div}}(O_{2,k+1})\mathbf{x}_k^L - \Delta t\, \mu_{\mathrm{eff}}(N, O_{2,k+1}) \odot \mathbf{x}_k^L, \tag{23}$$

where $\odot$ denotes element-wise multiplication over chromosome-count states.

Mass that becomes nonviable through the modeled stress pathway or failed divisions is transferred into the dead compartments, which contribute to tumor burden. The dead pool assigned by the model to resource stress is updated as

$$\mathbf{x}_{k+1}^{D,\mathrm{hyp}} = \mathbf{x}_k^{D,\mathrm{hyp}} + \Delta t\, \mu_{\mathrm{eff}}(N, O_{2,k+1}) \odot \mathbf{x}_k^L - \Delta t\, k_{\mathrm{clear}} \mathbf{x}_k^{D,\mathrm{hyp}}, \tag{24}$$

and the CIN-associated dead pool is updated as

$$\mathbf{x}_{k+1}^{D,\mathrm{CIN}} = \mathbf{x}_k^{D,\mathrm{CIN}} + \Delta t\, \boldsymbol{\phi}_{\mathrm{CIN}}(\mathbf{x}_k^L, O_{2,k+1}) - \Delta t\, k_{\mathrm{clear}} \mathbf{x}_k^{D,\mathrm{CIN}}. \tag{25}$$

Here $k_{\mathrm{clear}}$ is the first-order clearance rate for dead biomass, and $\boldsymbol{\phi}_{\mathrm{CIN}}$ is defined component-wise in Eq. (19). Together, the oxygen relaxation and Eqs. (23)–(25) define the coupled viable/dead population dynamics used in the simulations.

##### 1.3.1 Model calibration and observation models

Model calibration uses a common transformed-parameter optimizer for *in vivo*, *in vitro*, and joint fits. Natural-scale parameter specifications provide initialization values and admissible bounds; each component maps its fitted parameters to the optimizer scale before evaluating its objective. The mechanistic dynamics described above are shared, while the observation models differ between the *in vivo* and *in vitro* data.

***In vivo* observation model and likelihood.** The *in vivo* component uses three data modalities: longitudinal tumor burden, terminal necrosis where matched histology is available, and terminal single-cell chromosome-number profiles.

Tumor burden. For each mouse $i \in \mathcal{I}$, where $\mathcal{I}$ denotes the eight retained tumors used for calibration (section 1.1), tumor burden is measured at observation times $\{t_{ij}^{(V)}\}_{j=1}^{m_i}$, yielding observed tumor volumes

$$V_{ij}^{\text{obs}} \equiv V_i^{\text{obs}}(t_{ij}^{(V)}). \tag{26}$$

Let $x_{i,N}^{L}(t)$, $x_{i,N}^{D,\text{hyp}}(t)$, and $x_{i,N}^{D,\text{CIN}}(t)$ denote viable abundance, dead abundance assigned by the model to loss associated with resource stress, and CIN-associated dead abundance, respectively, for tumor $i$ in chromosome-count state $N$ at time $t$. The model predicts tumor volume by scaling total burden:

$$\widehat{V}_{ij} = \sum_{N} \left( x_{i,N}^{L}(t_{ij}^{(V)}) + x_{i,N}^{D,\text{hyp}}(t_{ij}^{(V)}) + x_{i,N}^{D,\text{CIN}}(t_{ij}^{(V)}) \right) v_{\text{cell}}, \tag{27}$$

where each cell is assigned the effective volume

$$v_{\text{cell}} = \frac{1}{\rho_{2N}}. \tag{28}$$

This observation model does not impose a ploidy-dependent cell-volume correction: low- and high-ploidy cells contribute the same effective per-cell volume to the bulk burden. Tumor-burden observations are modeled on the logarithmic scale,

$$\log V_{ij}^{\text{obs}} \sim \mathcal{N}\left(\log \widehat{V}_{ij}, \sigma_{\text{burden}}^2\right), \tag{29}$$

where $\sigma_{\text{burden}}$ is the burden measurement-noise scale. For longitudinal tumor burden, the fitted loss is a Gaussian negative log-likelihood on the logarithmic scale, averaged first within tumor and then across tumors. Let $\mathcal{J}_i$ denote the retained observation times for tumor $i$ after excluding day 0. Then

$$\mathcal{L}_B(\theta) = \frac{1}{|\mathcal{I}|} \sum_{i \in \mathcal{I}} \frac{1}{|\mathcal{J}_i|} \sum_{j \in \mathcal{J}_i} \left[ \log \sigma_{\text{burden}} + \frac{1}{2}\log(2\pi) + \frac{\left(\log V_{ij}^{\text{obs}} - \log \widehat{V}_{ij}\right)^2}{2\sigma_{\text{burden}}^2} \right]. \tag{30}$$

In implementation, each volume entering a logarithm is replaced by $\max(V, \varepsilon_{\log B})$ with $\varepsilon_{\log B} = 10^{-12}$ mm$^3$; this numerical floor is suppressed in Eqs. (29)–(30) for readability. Because $\sigma_{\text{burden}}$ is estimated jointly, the $\log \sigma_{\text{burden}}$ term is retained in the likelihood contribution. Omitting it would allow the optimizer to reduce the burden loss by inflating the assumed measurement noise.

Terminal necrosis. For tumors with matched terminal histology, necrosis is compared with the simulated terminal fraction of total dead biomass. This is an observation-model term, not an additional live-to-dead flux. The stress-associated death hazard $\mu_{\text{eff}}(N, O_2)$ contributes to $\mathbf{x}^{D,\text{hyp}}$, while mitotic nonviability and boundary-dropped offspring contribute to $\mathbf{x}^{D,\text{CIN}}$. The terminal necrosis comparison uses the sum of these two dead pools.

For tumor $i$, define the terminal total-dead volume and terminal total tumor volume as

$$\widehat{V}_i^{D,\text{tot}}(\theta) = v_{\text{cell}} \sum_N \left[ x_{i,N}^{D,\text{hyp}}(T_i^{(H)};\theta) + x_{i,N}^{D,\text{CIN}}(T_i^{(H)};\theta) \right], \qquad \widehat{V}_i^{\text{tot}}(\theta) = v_{\text{cell}} \sum_N \left[ x_{i,N}^{L}(T_i^{(H)};\theta) + x_{i,N}^{D,\text{hyp}}(T_i^{(H)};\theta) + \right. \tag{31}$$

The predicted terminal necrotic fraction is

$$\widehat{f}_i^{\text{nec}}(\theta) = \frac{\widehat{V}_i^{D,\text{tot}}(\theta)}{\widehat{V}_i^{\text{tot}}(\theta)}. \tag{32}$$

Because the same effective cell-volume factor appears in both numerator and denominator, it cancels in this ratio. Thus the fitted necrosis term constrains the terminal total-dead fraction of the simulated tumor burden, after both dead pools have accumulated their respective inflows and have been cleared by the shared first-order clearance process.

Let $f_i^{\text{nec,obs}}$ be the observed terminal necrotic fraction for tumors in the matched necrosis set $\mathcal{I}_{\text{nec}}$. Fractions are compared on a clipped logit scale. Define

$$\text{logit}_\epsilon(f) = \log\left(\frac{\widetilde{f}}{1-\widetilde{f}}\right), \qquad \widetilde{f} = \min\{1-\epsilon_{\text{nec}}, \max(\epsilon_{\text{nec}}, f)\}. \tag{33}$$

The terminal necrosis loss is the mean standardized squared discrepancy on this scale,

$$\mathcal{L}_{\text{nec}}(\theta) = \frac{1}{|\mathcal{I}_{\text{nec}}|} \sum_{i \in \mathcal{I}_{\text{nec}}} \left[ \frac{\text{logit}_\epsilon(\widehat{f}_i^{\text{nec}}(\theta)) - \text{logit}_\epsilon(f_i^{\text{nec,obs}})}{\sigma_{\text{nec}}} \right]^2. \tag{34}$$

Here $\sigma_{\text{nec}}$ sets the tolerated discrepancy in terminal necrosis on the logit-fraction scale, and $\epsilon_{\text{nec}}$ prevents fractions at 0 or 1 from producing infinite logits. In the run configuration these quantities are named `sigma_necrosis_logit` and `necrosis_fraction_eps`, respectively.

Ploidy. Each mouse has an initial ploidy group $g_i \in \{2N, 4N\}$ and harvest time $T_i^{(H)}$. Retained burden observations are nonzero-time measurements up to harvest, so the number and timing of observations may differ across mice.

For each tumor $i \in \mathcal{I}$, endpoint single-cell copy-number profiling provides modeled total autosomal chromosome-number estimates. If $n_i$ cells are profiled from tumor $i$, the observed chromosome-number values are denoted

$$z_{i\ell}, \qquad \ell = 1, \dots, n_i. \tag{35}$$

These single-cell observations sample the viable population at harvest. The model-predicted viable chromosome-number distribution at harvest is

$$\widehat{\pi}_{i,N}(\theta) = \frac{x^L_{i,N}(T_i^{(H)};\theta)}{\sum_{N'} x^L_{i,N'}(T_i^{(H)};\theta)}. \tag{36}$$

This distribution represents the predicted probability that a viable cell sampled from tumor $i$ at harvest has true chromosome count $N$.

Observed single-cell chromosome-number estimates are then modeled conditional on this predicted viable distribution. Specifically, each observed value $z_{i\ell}$ is treated as a noisy measurement of an unobserved true chromosome-count state $N$, with mixture weights given by $\widehat{\pi}_{i,N}(\theta)$:

$$p(z_{i\ell} \mid \theta) = \sum_N \widehat{\pi}_{i,N}(\theta)\, \mathcal{N}\big(z_{i\ell}; N, \sigma^2_{\text{ploidy}}\big)\,. \tag{37}$$

where $\sigma_{\text{ploidy}}$ represents measurement variability in single-cell chromosome-number estimates. For terminal single-cell chromosome-number data, the fitted loss first computes a tumor-level average negative log-likelihood. Define

$$\ell_i(\theta) = -\frac{1}{n_i}\sum_{\ell=1}^{n_i} \log p(z_{i\ell} \mid \theta). \tag{38}$$

Let $\mathcal{I}_{2N}$ and $\mathcal{I}_{4N}$ denote the retained tumors initiated from the near-diploid and near-tetraploid lineages, respectively. With both cohorts present, the endpoint contribution gives equal weight to the two initial-ploidy cohorts and equal weight to tumors within each cohort:

$$\mathcal{L}_C(\theta) = \frac{1}{2}\left[\frac{1}{|\mathcal{I}_{2N}|}\sum_{i\in\mathcal{I}_{2N}} \ell_i(\theta) + \frac{1}{|\mathcal{I}_{4N}|}\sum_{i\in\mathcal{I}_{4N}} \ell_i(\theta)\right]. \tag{39}$$

*Maximum-a-posteriori objective.* The *in vivo* component is formulated as a maximum-a-posteriori objective combining burden likelihood, chromosome-number likelihood, terminal necrosis loss, and soft-prior penalties:

$$\mathcal{L}_{\text{vivo}}(\theta) = \mathcal{L}_B(\theta) + \mathcal{L}_C(\theta) + \lambda_{\text{nec}}\,\mathcal{L}_{\text{nec}}(\theta) + \lambda_{\text{prior}}\,\mathcal{L}_{\text{prior}}(\theta), \tag{40}$$

where $\theta$ denotes the fitted parameter vector for this component, $\mathcal{L}_B$ is the burden contribution, $\mathcal{L}_C$ is the terminal chromosome-number contribution, $\mathcal{L}_{\text{nec}}$ is the terminal total-dead-fraction loss, $\lambda_{\text{nec}}$ is its relative objective weight, and $\mathcal{L}_{\text{prior}}$ is the soft-prior penalty.

For each prior-constrained parameter, let $z_i = T_i(\theta_i)$ denote the calibration-scale value used by the optimizer. The soft-prior penalty is

$$\mathcal{L}_{\text{prior}}(\theta) = \frac{1}{2}\sum_{i\in\mathcal{P}}\left(\frac{z_i-\mu_i}{\sigma_i}\right)^2, \tag{41}$$

where $\mathcal{P}$ is the subset of parameters with configured soft priors. The transform $T_i$ is the parameter-specific optimizer transform (including $\log_{10}$ transforms for several positive parameters and the identity transform where configured), and $\mu_i$ and $\sigma_i$ are the prior center and scale on that same calibration scale.

***In vitro* observation model and likelihood.** The *in vitro* component uses the same chromosome-number transition model but applies it to passage-structured cell-line data. Each passage segment $s$ has an initial cohort label, a fixed oxygen level during the segment, and a duration $\Delta t_s$; available growth observations are represented as segment-level growth-rate data rather than being restricted to a single initial–final cell-count pair. The model propagates the corresponding viable chromosome-number distribution through the segment and compares the simulated passage outcome with three measured data modalities: passage growth-rate observations, passage-level chromosome-number observations, and flow-density measurements. For each passage, the propagated chromosome-number distribution used for reseeding and endpoint comparison was taken from the simulated time point whose live-cell count was closest to the observed final or next-passage initial cell count.

For growth-rate data, let $g_s^{\text{obs}}$ be the observed passage growth rate and $\widehat{g}_s(\theta)$ the model-predicted growth rate for segment $s$. The averaged growth log-likelihood is

$$\bar{\ell}_{\text{growth}}(\theta) = \frac{1}{|\mathcal{S}_g|} \sum_{s \in \mathcal{S}_g} \log \mathcal{N}\Big(g_s^{\text{obs}}; \widehat{g}_s(\theta), \sigma_{\text{growth}}^2\Big), \tag{42}$$

where $\mathcal{S}_g$ is the set of segments with observed growth-rate data and $\sigma_{\text{growth}}$ is the growth measurement-noise scale.

For chromosome-number observations in segment $s$, let $\pi_{s,N}$ denote the predicted viable chromosome-number distribution at the comparison time and let $z_{s\ell}$ be the observed integer chromosome-number value for cell $\ell$. The implementation first smooths the predicted distribution on the finite modeled chromosome grid $\mathcal{N}$. For an observed-grid state $m$ and true modeled state $N$, define the column-normalized Gaussian kernel

$$K_{\sigma_{\text{kary}}}(m \mid N) = \frac{\phi\Big(m; N, \sigma_{\text{kary}}^2\Big)}{\sum_{r \in \mathcal{N}} \phi\big(r; N, \sigma_{\text{kary}}^2\big)}, \qquad m, N \in \mathcal{N}, \tag{43}$$

where $\phi(\cdot; N, \sigma_{\text{kary}}^2)$ is a Gaussian density evaluated only on the discrete chromosome grid. The smoothed discrete probability mass is

$$\widetilde{\pi}_{s,m} = \sum_{N \in \mathcal{N}} K_{\sigma_{\text{kary}}}(m \mid N) \pi_{s,N}. \tag{44}$$

The averaged chromosome-number log-likelihood is then

$$\bar{\ell}_{\text{ploidy}}(\theta) = \frac{1}{|\mathcal{S}_z|} \sum_{s \in \mathcal{S}_z} \frac{1}{n_s} \sum_{\ell=1}^{n_s} \log\left[\max\left\{\widetilde{\pi}_{s,z_{s\ell}}, 10^{-12}\right\}\right], \tag{45}$$

where $\mathcal{S}_z$ is the set of passage comparisons with chromosome-number observations, $n_s$ is the number of observations in segment $s$, and $\sigma_{\text{kary}}$ is the chromosome-number smoothing scale for the *in vitro* karyotype term. Observations that do not match a modeled grid state receive the same $10^{-12}$ probability floor. Thus, this term is a discrete grid likelihood after finite-grid kernel smoothing, not a continuous Gaussian-mixture density.

For flow-density data, let $\omega_{sp}^{\text{obs}}$ be the normalized observed density mass at flow grid point $p$ for segment $s$, and let $\widehat{f}_s(p;\theta)$ be the model-predicted density after Gaussian smoothing of the simulated viable chromosome-state distribution onto the observed flow grid and renormalization over that grid. The Gaussian width was set separately from the fitted karyotype smoothing parameter to the larger of twice the median flow-grid spacing and 1/22 ploidy units; for the seed-144 grid this was 0.0804020100502587. The averaged flow log-likelihood is

$$\bar{\ell}_{\text{flow}}(\theta) = \frac{1}{|\mathcal{S}_f|} \sum_{s \in \mathcal{S}_f} \sum_{p \in \mathcal{G}_s} \omega_{sp}^{\text{obs}} \log \left[ \max \left\{ \widehat{f}_s(p;\theta), 10^{-12} \right\} \right], \tag{46}$$

where $\mathcal{S}_f$ is the set of flow-profile comparisons and $\mathcal{G}_s$ is the observed flow grid for segment $s$. The $10^{-12}$ floor is a numerical safeguard applied after smoothing and normalization.

The fitted *in vitro* objective is the negative weighted mean log-likelihood

$$\mathcal{L}_{\text{vitro}}(\theta) = -\left[ w_{\text{growth}}\, \bar{\ell}_{\text{growth}}(\theta) + w_{\text{ploidy}}\, \bar{\ell}_{\text{ploidy}}(\theta) + w_{\text{flow}}\, \bar{\ell}_{\text{flow}}(\theta) \right]. \tag{47}$$

The three terms compare the simulated passage lineages with growth-rate, chromosome-number, and flow-density data, respectively. The reported fits used equal component weights, $w_{\text{growth}} = w_{\text{ploidy}} = w_{\text{flow}} = 1$ (Supplementary Table 12).

**Joint fitting of *in vivo* and *in vitro* data.** The joint fit uses a single optimizer vector while evaluating separate transformed parameter vectors for the *in vivo* and *in vitro* likelihood components. Soft-coupled parameters are represented by a center–delta parameterization: for each soft-coupled parameter, the optimizer carries one transformed center variable, $c_Q$, and one additional offset variable, $\delta_Q$. The context-specific transformed values are reconstructed from these two optimizer variables during objective evaluation. Parameters that appear only in one observation model are retained as context-specific observation-model parameters. In the analysis reported here, all 14 overlapping active biological parameters—$O_{2,c}$, $\mu_{\text{hp}}$, $p_{\text{misseg}}$, $k_{o,\text{mis}}$, $s_{\max}$, $\beta_{\text{buf}}$, $n_{\exp}$, $n_O$, $\alpha_{O_2}$, $\gamma_{\text{growth}}$, $\lambda_{\max}$, $p_{\text{mis,base}}$, $p_{\text{wgd}}$, and $\gamma_\mu$—were soft-coupled across contexts; no overlapping active biological parameter was hard-shared (Supplementary Table 11). Thus, the strength and chromosome-number dependence of resource-stress growth suppression were allowed to differ between the *in vivo* and *in vitro* components, while regularization discouraged unsupported divergence.

The minimized joint objective is

$$\mathcal{L}_{\text{joint}}(\theta) = w_{\text{vivo}}\mathcal{L}_{\text{vivo}}(\theta_{\text{vivo}}) + w_{\text{vitro}}\mathcal{L}_{\text{vitro}}(\theta_{\text{vitro}}) + \mathcal{L}_{\text{soft}}(\theta), \tag{48}$$

where $\theta_{\text{vivo}}$ and $\theta_{\text{vitro}}$ are the context-specific parameter vectors reconstructed from the joint optimizer vector and passed to the two likelihood components. The reported analysis used $w_{\text{vivo}} = w_{\text{vitro}} = 1$ (Supplementary Table 12). For each soft-coupled natural-scale parameter $Q$, let

$$z_Q = T_Q(Q)$$

denote its transformed optimizer-scale value, where $T_Q$ is the transformation used for that parameter during optimization. The separate *in vivo* and *in vitro* fits define context-specific transformed bounds $[l_Q^{\text{vivo}}, u_Q^{\text{vivo}}]$ and $[l_Q^{\text{vitro}}, u_Q^{\text{vitro}}]$. In the joint fit, these component-specific bounds are used only to define a single active joint admissible interval:

$$l_Q^{\text{joint}} = \min\bigl(l_Q^{\text{vivo}}, l_Q^{\text{vitro}}\bigr), \qquad u_Q^{\text{joint}} = \max\bigl(u_Q^{\text{vivo}}, u_Q^{\text{vitro}}\bigr). \tag{49}$$

Thus, the original component-specific bounds record the provenance of the separate fits, whereas the joint optimizer uses the union interval $[l_Q^{\text{joint}}, u_Q^{\text{joint}}]$ as the active transformed-scale admissible range. During objective evaluation, the transformed values passed to the two components are reconstructed as

$$z_Q^{\text{vivo}} = c_Q + \frac{\delta_Q}{2}, \qquad z_Q^{\text{vitro}} = c_Q - \frac{\delta_Q}{2}. \tag{50}$$

The reconstructed values must satisfy

$$z_Q^{\text{vivo}} \in [l_Q^{\text{joint}}, u_Q^{\text{joint}}], \qquad z_Q^{\text{vitro}} \in [l_Q^{\text{joint}}, u_Q^{\text{joint}}]. \tag{51}$$

Parameter vectors that do not yield admissible reconstructed transformed values are outside the joint feasible domain and are not evaluated as valid likelihood points. The soft-coupling penalty is applied to the offset variable,

$$\mathcal{L}_{\text{soft}}(\theta) = \sum_{Q \in \mathcal{S}} \frac{c^2}{2} \left[ 1 - \exp\left\{ -\left( \frac{\delta_Q}{c\sigma_Q} \right)^2 \right\} \right], \tag{52}$$

where $\mathcal{S}$ is the set of soft-coupled parameters, $\sigma_Q$ is the transformed-scale soft-coupling scale, and $c$ controls the transition from near-quadratic behavior to saturation. For the reported fits, $\sigma_Q = 0.65$ for all 14 parameters and $c = 0.4$ (Supplementary Table 12). For $|\delta_Q| \ll c\sigma_Q$, each term approaches $\frac{1}{2}(\delta_Q/\sigma_Q)^2$; for large separations it saturates at $c^2/2$, limiting the influence of a single strongly context-specific parameter. Since

$$z_Q^{\text{vivo}} - z_Q^{\text{vitro}} = \delta_Q,$$

this penalty regularizes the transformed difference between the *in vivo* and *in vitro* parameter values. For log-transformed parameters, this corresponds to regularization of fold differences rather than absolute natural-scale differences.

**Differential-evolution initial-population and optimizer-endpoint analysis.** For Figure 5D, "initial distribution" denotes the population passed to differential evolution at the start of each joint optimization. It is not a distribution induced from the configured fitting intervals and is not obtained by weighting candidate values with the Welsch objective penalty. Endpoint ensembles were defined for C01 and C02 using separate-*in vivo* seeds 366 and 25, respectively. Each retained *in vivo* warm start was paired with the common separate-*in vitro* seed-144 anchor (Supplementary Tables 7 and 8).

The initial populations were deterministically reconstructed with the fitting code and saved run inputs. For each C family, joint seeds 1–500 were reset individually and the same `joint_deoptim_initial_population` routine used by the fit was called. The optimizer contained 40 coordinates, so the enforced population size was

$$N_P = \max(80, 10 \times 40) = 400.$$

Warm-start initialization was enabled with $\sigma_N = 0.1$. For an ordinary transformed coordinate with warm-start value $z_0$, the routine sampled a normal distribution centered at $z_0$, with standard deviation $\sigma_N|z_0|$, truncated to the active optimizer bounds. When $|z_0| \leq 10^{-12}$, the active bound width replaced $|z_0|$ as the scale reference. For each soft-coupled parameter, the center was sampled first; the feasible bounds for the transformed context difference $\delta_Q$ were then recomputed from that sampled center so that both reconstructed context values remained inside the joint union interval. The delta coordinate was sampled from a truncated normal centered on the warm-start delta after clipping that mean to its center-dependent feasible interval, using the same scale rule. The first member of every 400-member population was replaced by the exact warm-start vector. Thus, each family–parameter initial distribution contained 200,000 values (500 seeds times 400 population members; Supplementary Table 8).

For every soft-coupled coordinate, the transformed context values were reconstructed as

$$z_Q^{\text{vivo}} = c_Q + \frac{\delta_Q}{2}, \qquad z_Q^{\text{vitro}} = c_Q - \frac{\delta_Q}{2}. \tag{53}$$

For $\log_{10}$-transformed parameters, natural values were obtained as

$$Q_{\text{vivo}} = 10^{z_Q^{\text{vivo}}}, \qquad Q_{\text{vitro}} = 10^{z_Q^{\text{vitro}}}; \tag{54}$$

identity-scale coordinates already equaled the natural context values. Figure 5D displays these two natural-scale marginals separately rather than plotting their ratio. For every C family and coupled parameter, all 500 endpoints were required to have finite positive context values, to satisfy the joint feasible-domain check, and to require no post-optimization projection (Supplementary Table 8).

For each context-specific DE initial population and its corresponding endpoint ensemble, the natural-scale median and type-8 5th and 95th percentiles were calculated. The percentiles were retained for the numerical concentration audit and its source tables, but Figure 5D displays only optimizer-endpoint medians, as asterisks on the shared coordinate axis; initial-population median points and separate percentile spans are not drawn. Kernel densities were evaluated on a 401-point parameter-specific grid shared by the two contexts, both numerical roles, and both families. Density estimation used the configured optimizer scale—$\log_{10}$ for log-transformed parameters and identity otherwise—while logarithmic axes were labeled with natural parameter values. Each density was divided by its own maximum; therefore only horizontal location, width, and shape, not vertical peak height, were compared (Supplementary Table 8).

The row order in Figure 5D used the single best retained joint winner from each C family. For parameter $Q$ and family $f \in \{\text{C01}, \text{C02}\}$, the natural-scale ratio was $R_{Qf} = Q_f^{\text{vivo}}/Q_f^{\text{vitro}}$. Parameters were ordered by increasing $\left|\frac{1}{2}\sum_f R_{Qf} - 1\right|$, with parameter name as the deterministic tie breaker. This ordering is a display rule for proximity to cross-context parity, not an inferential test.

Boundary proximity in Figure 5D was defined from the complete joint-union fitting interval, not from a distribution quantile. For lower and upper bounds $L_Q$ and $U_Q$ on the optimizer coordinate, the two red background regions were

$$[L_Q,\, L_Q + 0.05(U_Q - L_Q)] \quad \text{and} \quad [L_Q + 0.95(U_Q - L_Q),\, U_Q]\,. \tag{55}$$

Thus each region occupies exactly 5% of the plotted horizontal interval. For $\log_{10}$-optimized parameters these limits were calculated in $\log_{10}$ optimizer space and the tick labels were then converted back to natural units; identity-scale parameters required no conversion. The two retained families used the same joint-union bounds for each parameter.

As an auxiliary directional audit, the paired values were also transformed to $\log_2(Q_{\text{vivo}}/Q_{\text{vitro}})$. A parameter-family contrast was called higher *in vivo* when its endpoint 5th percentile exceeded zero, lower *in vivo* when its 95th percentile was below zero, and overlapping equality otherwise. Cross-family direction agreement required the same non-overlap classification in both families. Paired-ratio fitted-solution concentration relative to initialization was summarized as

$$R_{W,Qf} = \frac{q_{0.95,Qf}^{\text{endpoint}} - q_{0.05,Qf}^{\text{endpoint}}}{q_{0.95,Qf}^{\text{DE init}} - q_{0.05,Qf}^{\text{DE init}}}. \tag{56}$$

We additionally reported, separately for *in vivo* and *in vitro*, the fraction of the 500 endpoints that lay at the corresponding active parameter bound (Supplementary Table 8).

The fixed red edge bands show whether a fitted marginal approaches the extremes of the declared joint fitting domain, while the source-table 5th–95th percentile summaries and $R_{W,Qf}$ provide the separate numerical-concentration audit. Both characterize numerical-search behavior under the tested warm starts, bounds, objective, and search settings. A zero endpoint span can result from many starts converging to the same endpoint, whereas high active-bound occupancy indicates optimizer pressure against the declared fitting domain. These summaries assess optimizer concentration rather than posterior uncertainty, biological replication, or structural identifiability.

**Warm-start initialization for the joint fit.** The joint analysis used two landscape-informed separate-*in vivo* fits as warm-start representatives. C01 and C02 denote the two saved primary regions obtained by clustering the 500 final *in vivo* positions in the pooled 14-parameter t-SNE embedding described below; no secondary clustering was used. The retained $k = 2$ partition had average silhouette 0.673. This embedding and its labels are distinct from the *in vivo*-only 18-parameter landscape in Figure 4C. The cluster coordinates are numerical-search design variables rather than mechanistic variables, biological subtypes, or experimental groups (Supplementary Figure 6; Supplementary Table 7).

Within the two selected primary regions, the retained separate-*in vivo* representatives were seeds 366 and 25 for C01 and C02, respectively. Each was paired with the same lowest-objective separate-*in vitro* fit (seed 144) on the transformed optimizer scale. The main design therefore compares two *in vivo* primary regions against one fixed *in vitro* anchor (Supplementary Table 7).

For each soft-coupled parameter $Q$, let $z_Q^{\text{vivo},0}$ and $z_Q^{\text{vitro},0}$ denote the transformed values from one such C-family configuration. The initial center and offset are set to

$$c_Q^0 = \frac{z_Q^{\text{vivo},0} + z_Q^{\text{vitro},0}}{2}, \qquad \delta_Q^0 = z_Q^{\text{vivo},0} - z_Q^{\text{vitro},0}. \tag{57}$$

This initialization exactly reconstructs the two separate-fit transformed values:

$$z_Q^{\text{vivo},0} = c_Q^0 + \frac{\delta_Q^0}{2}, \qquad z_Q^{\text{vitro},0} = c_Q^0 - \frac{\delta_Q^0}{2}.$$

Parameters that appear only in one component, such as context-specific observation-model parameters, are initialized from the corresponding separate-fit solution.

The warm start must be representable under the active joint union bounds in Eq. (49); otherwise, initialization fails explicitly. This ensures that the joint fit begins from the selected separate-fit solutions only when those solutions are valid under the declared joint feasible domain.

**Optimization and data inclusion.** Optimization attempted a parallel differential-evolution global stage followed by serial L-BFGS-B refinement; a local result was retained only when it passed the implementation checks and improved the objective. Separate *in vivo* and *in vitro* searches each used 500 numerical seeds. In the main joint analysis, C01 and C02 each used 500 numerical seeds, for 1,000 optimizer runs. Both selected joint winners retained the differential-evolution endpoint after the local step was not accepted (code 52); completed differential-evolution iterations ranged from 26–95 in C01 and 26–45 in C02. The selected winners had two and one of 20 active parameters at configured bounds, respectively. In the separate *in vitro* search, all 500 runs recorded the differential-evolution flag, 495 accepted a local result, and the retained seed-144 local step was accepted with code 1 while five of 20 active parameters were at bounds. In the separate *in vivo* search, all 500 runs recorded the differential-evolution flag, 460 accepted a local result, 13 objectives were within 1% and 225 were within 5% of seed 25, and the selected seed-25 local step was accepted with code 0 while one of 20 active parameters was at a bound. The corresponding diagnostics are summarized in Figure 3G, Supplementary Figures 3 and 2, and Supplementary Tables 6, 4, and 7. Throughout, "best" denotes the lowest retained objective among the evaluated searches, and seed-to-seed spread quantifies numerical-search behavior.

For the Figure 6A multi-seed analysis, eligibility was defined independently within C01 and C02 because the families had different objective baselines. A retained endpoint first had to have a finite recorded full joint MAP objective, a valid unique seed identifier, exactly one finite value for each of the 14 response parameters in both contexts, feasibility both before projection and at the reported solution, no applied parameter projection, and an available local model configuration. Eligible endpoints were ordered by the complete retained joint MAP objective, including the fitted-data terms and soft-coupling penalty. The primary ensemble was the lowest objective decile within each family (50 of 500 endpoints); nested lowest-5% (25 endpoints) and lowest-20% (100 endpoints) sets were used as threshold-sensitivity analyses. Within an eligibility set, endpoints received equal seed weight; objective values determined inclusion without posterior-like weighting. Exact natural-scale parameter signatures identified 50 unique C01 endpoints and 16 unique C02 endpoints in the q10 sets. Each unique representative was evaluated once and original optimizer-seed multiplicity was restored for seed-weighted summaries; equal-weight unique-endpoint summaries were retained as sensitivity analyses. All 200 q20 endpoint/operator combinations passed, all 16 audited modal results were invariant from q05 through q20, and all 48 seed-weighted versus unique-endpoint modal comparisons agreed (Supplementary Figure 6; Supplementary Tables 7, 9, and 10). These thresholds and weights quantify numerical-search robustness.

**Optimizer-ensemble sensitivity analyses.** For coupled parameter $Q$, warm-start family $f \in \{\mathrm{C01}, \mathrm{C02}\}$, and numerical start $s \in \{1, \ldots, 500\}$, the context contrast was

$$D_{Qfs} = \log_2\left(\frac{Q_{fs}^{\mathrm{vivo}}}{Q_{fs}^{\mathrm{vitro}}}\right). \tag{58}$$

Within each C family, the 500 endpoints contributed equal weight to the context-specific initialization and optimizer-endpoint distributions shown in Figure 5D. Natural-scale ratios were classified as higher *in vitro* for $0 < Q_{\text{vivo}}/Q_{\text{vitro}} \leq 0.8$, approximately equal for $0.8 < Q_{\text{vivo}}/Q_{\text{vitro}} < 1.2$, and higher *in vivo* for ratios at least 1.2. Figure 5D reports C01 and C02 separately. Because both families share the same *in vitro* anchor, cross-family agreement quantifies numerical-search robustness across the two *in vivo* regions. These analyses characterize optimization and warm-start sensitivity (Supplementary Tables 7 and 8).

Retained *in vivo* observation times satisfy $0 < t_{i,1} < \cdots < t_{i,n_i} \leq T_i^{(H)}$, so day-0 burden points are excluded and longitudinal follow-up is administratively truncated at harvest. No censoring likelihood was used. Only tumors with both burden and terminal chromosome-number data are retained. Terminal endpoint discrepancies are evaluated directly from the single-cell chromosome-number observations under the Gaussian-mixture likelihood in Eq. (37), and tumor-level endpoint losses are aggregated equally within each initial-ploidy cohort before applying cohort balancing as in Eq. (39).

**Representative in vivo trajectory and terminal-distribution visualization.** The lowest-objective retained separate *in vivo* fit (seed 25) was propagated at one-day reporting intervals for the eight paired tumors. The four 2N-initiated and four 4N-initiated trajectories were summarized separately (Supplementary Tables 1 and 2). At each day, the viable model state, the dead model state assigned to loss associated with resource stress, and the CIN-associated dead model state were converted to tumor-volume components and averaged over the tumor trajectories available on that day before being plotted as stacked areas. The fitted effective oxygen state was plotted directly in percent. Viable-cell mean ploidy was calculated within each tumor as

$$\overline{P}_i(t) = \frac{\sum_{N\in\mathcal{N}} N\, x_{i,N}^L(t)}{N_{\text{unit}} \sum_{N\in\mathcal{N}} x_{i,N}^L(t)}, \qquad N_{\text{unit}} = 22. \tag{59}$$

The tumor-specific effective-$O_2$ and $\overline{P}_i(t)$ trajectories were then averaged over the same available trajectories at each day. Both cohort means were displayed without standardization on the numerical 0–5 left axis, while burden used the right axis. The shared coordinate axis visually aligns quantities with different physical units. Both longitudinal panels use a 0–100-day x-axis, and model curves end at the last available modeled time.

Observed tumor burden was summarized only at recorded measurement days. For cohort $c$, day $t$, and $n_{ct}$ observed tumors, the black curve is the arithmetic mean $\overline{B}_c^{\text{obs}}(t)$, and the gray band is $\overline{B}_c^{\text{obs}}(t) \pm s_c(t)$, where $s_c(t)$ is the between-tumor sample standard deviation, truncated at zero for the lower display boundary. All available observations at each recorded day contributed to these descriptive summaries. The band is not measurement error, a confidence interval, or model uncertainty. The 120 burden records inside the configured cohort and time scope were cross-checked by tumor and day between the independently prepared Figure 1 and Figure 4 analysis tables, with exact agreement; the eight day-0 records were excluded from the burden likelihood, leaving 112 loss-contributing observations (Supplementary Table 1).

For the terminal comparison, the observed single-cell chromosome-number values and predicted terminal chromosome-state weights from the seed-25 fit were converted to ploidy by division by $N_{\text{unit}} = 22$. Within each cohort and source, each tumor was normalized to unit total weight and then assigned one-quarter of the cohort distribution. This equal-tumor weighting prevents tumors with more profiled cells from dominating the observed cohort distribution. Observed and predicted distributions were displayed on the common ploidy range 1–7 with weighted box summaries (Supplementary Tables 1 and 2).

**Fixed-oxygen attractor analysis.** For each of the 500 separate *in vivo* parameter sets, the chromosome-number model was evaluated at fixed oxygen without parameter refitting. At each oxygen level, the chromosome-transition generator was combined with the state-specific effective death rate to form

$$M(O_2) = G_{\text{div}}(O_2) - \text{diag}\{\mu_{\text{eff}}(N, O_2)\}\,. \tag{60}$$

The eigenvalue of $M$ with the largest real part, $\lambda_1$, was calculated numerically, and the real part of its associated eigenvector was retained. The eigenvector sign was oriented to have a non-negative sum, negative numerical components were truncated to zero, and the vector was normalized to unit mass. With normalized dominant eigenvector $v_N$, dominant mean ploidy was

$$\overline{P}_{\text{dom}}(O_2) = \frac{1}{N_{\text{unit}}} \sum_{N\in\mathcal{N}} N v_N. \tag{61}$$

The spectral gap was defined as the difference between the largest and second-largest eigenvalue real parts, $\text{Re}(\lambda_1) - \text{Re}(\lambda_2)$. The operator was evaluated at 201 fixed oxygen concentrations from 0% through 5% $O_2$ in increments of 0.025 percentage points. The continuous dominant mean ploidy from Eq. (61), without thresholding at ploidy 2, was retained for the Figure 4B association analysis (Supplementary Tables 3 and 4). These are analytical evaluations of the fitted linear operator, not stochastic simulations or oxygen-intervention experiments.

**Post-fit continuous single-parameter ploidy-association analysis.** At each of the 201 oxygen concentrations, the analysis units were 500 parameter sets obtained from distinct numerical optimizer seeds. Each of 18 prespecified fitted *in vivo* parameters was analyzed separately; the observation/volume parameters $\rho_{2N}$ and $\sigma_{\text{burden}}$ were excluded (Supplementary Tables 3 and 4). Let $R_{\theta,ij}$ be the average rank of fitted parameter $\theta$ across the 500 endpoints and let $R_{P,ij}$ be the average rank of continuous dominant mean ploidy at oxygen value $O_{2,j}$. Spearman association was calculated as

$$\rho_\theta(O_{2,j}) = \frac{\text{cov}(R_{\theta,ij}, R_{P,ij})}{\text{sd}(R_{\theta,ij})\,\text{sd}(R_{P,ij})}. \tag{62}$$

Ties received average ranks. Natural-scale parameter values were used for this calculation; applying the configured $\log_{10}$ transformation to strictly positive parameters gives the same ranks and therefore the same $\rho$.

For each parameter, the displayed peak association magnitude was

$$M_\theta = \max_{j=1,\ldots,201} |\rho_\theta(O_{2,j})|\,. \tag{63}$$

Let $j^*_\theta$ be the lowest oxygen-grid index attaining $M_\theta$. Figure 4B rows were first divided by the sign of $\rho_\theta(O_{2,j^*_\theta})$, with the positive group displayed before the negative group, and were then ordered by decreasing $M_\theta$ within each group; configured parameter order was used only to break an exact tie.

The adjacent dot position reports $M_\theta$, while its fill reports the signed value $\rho_\theta(O_{2,j^*_\theta})$. Selecting a maximum over 201 oxygen values emphasizes the strongest oxygen-specific association and does not measure persistence across the grid. The full signed heatmap is therefore retained to show whether that peak direction persists or reverses with oxygen. Neither $M_\theta$ nor the signed peak correlation is a causal effect, hypothesis-test statistic, multivariable importance estimate, or out-of-sample prediction measure.

**Separate *in vivo* parameter-landscape embedding and exploratory clustering.** For Figure 4C, the differential-evolution initial population was deterministically reconstructed for each of the 500 separate *in vivo* fits from its optimizer seed, ordered 20-variable parameter table, transformed bounds, and effective initialization rule. The saved configuration recorded hybrid initialization (`de_init_mode` = `hybrid`). The single-stage backend explicitly used uniform initialization. All 500 fit logs recorded `init_mode` = `uniform`, zero local candidates, and 255 uniform candidates after the parameter-table initializer. The full 20-variable population was reconstructed before feature selection to preserve the optimizer random-number sequence. Candidate 1, which reproduced the parameter-table initializer, was then removed from every 256-member population. The observation/calibration variables $\rho_{2N}$ and $\sigma_{\text{burden}}$ were excluded, leaving 18 mechanistic parameters and 127,500 retained initial-population rows. These were combined with the 500 best endpoints, giving 128,000 *in vivo*-only parameter vectors. Each parameter was transformed on its configured natural or $\log_{10}$ optimizer scale and z-scored across the combined data. The vectors were embedded in two dimensions by t-SNE using random seed 123, perplexity 30, Barnes–Hut approximation parameter $\theta = 0.5$, and 1,000 iterations (Supplementary Table 4).

The 500 best endpoints were retained and k-means clustering was applied to their two-dimensional t-SNE coordinates for $k = 2, \ldots, 8$. The value maximizing the average silhouette coefficient was selected; this procedure selected $k = 2$ with average silhouette 0.685055 and cluster sizes of 53 and 447 solutions (Supplementary Table 4). Figure 4B displays each original parameter value on a shared $\log_{10}$ coordinate, independently of whether its optimizer transformation was logarithmic or identity. Strictly positive values were mapped to $\log_{10}(x)$. Because one configured fitting interval had a zero lower bound, a dedicated display coordinate one decade below the smallest positive bound, initial value, or fitted endpoint was labeled "0"; in this analysis the display floor was $10^{-7}$. The upper blue half violin is a kernel-density estimate of the 500 best retained endpoints on the displayed $\log_{10}$ coordinate, and its green point marks seed 25, verified as rank 1 under the canonical 500-seed fit objective. The lower gray glyph spans the configured fitting bounds and its black point marks the configured initial value. Together, these elements summarize calibration settings and optimizer-endpoint variation.

For Supplementary Figure 1, each natural-scale endpoint was instead transformed on its configured optimizer scale and mapped to the unit interval defined by its configured transformed lower and upper bounds. The plotted prior-referenced position was $z_{\text{prior}} = (u - 0.5)/\sqrt{1/12}$, where $u$ is the bound-scaled value; zero therefore denotes the midpoint of the configured interval and one unit is the standard deviation of a uniform distribution over that interval. The endpoint values were stratified by the two exploratory t-SNE groups and shown in the same positive-peak then negative-peak order as Figure 4B, with decreasing $M_\theta$ within each group. These distributions and percentiles summarize optimizer-derived fitted endpoints on the configured scale. Solid t-SNE boundaries are convex hulls of all assigned endpoints and provide descriptive membership outlines. Because the clusters and displayed parameters were derived from the same fitted ensemble, cluster-associated parameter differences quantify internal fitted-landscape structure (Supplementary Table 4).

For the exploratory Figure 4D stratification, the two cluster distributions were compared separately for each of the 18 parameters with a Kruskal–Wallis rank-sum test on $z_{\text{prior}}$ (Supplementary Table 4). This transformation is monotone within each parameter, so it preserves the ranks and test statistic obtained from the corresponding configured transformed parameter values. The 18 omnibus $P$ values were adjusted together by the Benjamini–Hochberg procedure. For parameter $\theta$, the displayed effect size was

$$\epsilon_\theta^2 = \max\left\{0, \frac{H_\theta - k + 1}{n - k}\right\}, \tag{64}$$

where $H_\theta$ is the Kruskal–Wallis statistic, $k = 2$ groups, and $n = 500$ fitted endpoints. Parameters with adjusted $q < 0.05$ were ranked by decreasing $\epsilon_\theta^2$; adjusted $q$ and then the Figure 4B display order were prespecified tie breakers, and the first six were shown. The analysis units are numerical optimizer endpoints. Because the tested parameters were also inputs to the t-SNE and cluster assignment, the adjusted values and effect sizes quantify internal cluster separation.

**Pooled parameter-landscape embedding and joint-fit warm-start selection.** A separate pooled landscape was constructed from optimizer population samples and the final fitted solutions from the separate *in vivo* and *in vitro* calibrations. This landscape was used to design the joint-fitting warm starts shown in Figure 5C and is distinct from both the optimized joint-fit endpoints and the 18-parameter *in vivo*-only landscape above. Fourteen parameters shared by the two contexts were used: $\lambda_{\max}$, $p_{\text{mis,base}}$, $p_{\text{misseg}}$, $k_{o,\text{mis}}$, $s_{\max}$, $\beta_{\text{buf}}$, $n_{\exp}$, $p_{\text{wgd}}$, $\alpha_{O_2}$, $\gamma_{\text{growth}}$, $\mu_{\text{hp}}$, $\gamma_\mu$, $O_{2,c}$, and $n_O$. Parameters were transformed on their configured natural or $\log_{10}$ scales and standardized jointly across the pooled data. The resulting 256,000 parameter vectors—127,500 population points and 500 final solutions from each context—were embedded in two dimensions by t-SNE[38] using random seed 123, perplexity 30, Barnes–Hut approximation parameter $\theta = 0.5$, and 1,000 iterations. The unitless t-SNE axes approximate local neighborhood structure and have no direct mechanistic interpretation. The selected C01/C02 regions and their robustness audit are specified above, in Supplementary Figure 6, and in Supplementary Table 7.

**Weak-gap dominant-ploidy regime robustness.** For Supplementary Figure 7, we reused the same 201 oxygen values, 60 log-spaced fixed mechanistic $p_{\text{misseg}}$ values, and C01/C02 lowest-objective 50-endpoint ensembles used in Figure 6A. The upper *in vivo* rows constitute the primary result in this figure; the lower *in vitro* rows are retained as paired context, while the dedicated joint-fit culture extension is reported in Supplementary Figure 8. Exact duplicate 14-parameter endpoints were evaluated once and restored to their original optimizer-seed multiplicities before pointwise aggregation; the q10 ensembles contained 50 unique C01 endpoints and 16 unique C02 endpoints (Supplementary Tables 9 and 10). The rank-1 eigenvector was oriented to have positive total mass and normalized with the same nonnegative rule used for Figure 6A. Its chromosome-number-weighted mean divided by $N_{\text{unit}}$ defined dominant mean ploidy $P_{esc}(O_2, p)$ for endpoint $e$, family $s$, context $c$, oxygen concentration $O_2$, and mechanistic input $p = p_{\text{misseg}}$. Pointwise rank-1 medians and percentiles were required to agree with the corresponding context-specific Figure 6A source surface to numerical tolerance.

A grid cell was classified as weak-gap when at least half of its 50 seed-weighted endpoints had

$\mathrm{Re}(\lambda_1) - \mathrm{Re}(\lambda_2) < 0.005$, matching the Figure 6A overlay. For each endpoint, dominant mean ploidy was classified as low when $P_{esc} \leq 2$, intermediate when $2 < P_{esc} < 4$, and high when $P_{esc} \geq 4$. At each grid cell, the largest of the three corresponding endpoint fractions defined regime consensus; a cell was assigned a stable class at consensus at least 0.9 and was called mixed otherwise. Exact across-fit dispersion was reported as the 90th minus 10th percentile of $P_{esc}$ (Supplementary Table 10).

Local grid sensitivity was calculated independently for every context-specific endpoint vector before aggregation. The neighborhood of a grid cell contained its available immediately adjacent oxygen values and immediately adjacent fixed-missegregation values, for at most four neighbors. The endpoint-level regime-switch indicator equaled one when any neighbor changed among the low, intermediate, and high classes defined above. The corresponding local-change amplitude was the largest absolute difference in $P_{esc}$ among those neighbors. Supplementary Figure 7B reports the endpoint fraction with a local three-class switch, panel C reports the across-endpoint percentile spread at the focal cell, and panel D reports the empirical cumulative distribution of the endpoint-median local-change amplitude across weak-gap cells. These quantities were computed for both displayed rows, but the manuscript interpretation of this supplementary figure is based on the *in vivo* rows. Boundary cells used only their available neighbors. These grid-local quantities measure sensitivity to one numerical evaluation step; they are not derivatives, finite-time transition rates, probabilities that cells switch state, or biological confidence levels. Higher eigenmodes were retained in the reproducibility audit but were not used as alternative population distributions because non-dominant eigenvectors may be signed or complex.

**Extended joint-fit culture oxygen response.** Supplementary Figure 8 is the dedicated joint-fit *in vitro* extension. Panel A applies the separate-fit response classifier over 0–20% oxygen to provide landscape context. Panels B–E evaluate the C01/C02 joint-fit culture parameter vectors over the extended oxygen domain using the same fixed mechanistic-$p_{\mathrm{misseg}}$, weak-gap, endpoint-multiplicity, and local-sensitivity definitions as the corresponding primary analyses. The 5% cutoff defines the primary analysis range rather than the culture calibration limit: the culture data include a 20.5% $O_2$ control. The extended grid consists of prescribed constant-oxygen and altered-missegregation scenarios, not additional fitted observations or a replay of the measured passage schedule.

**Fixed-oxygen response-curve classification and reliability.** For each final separate-fit parameter set in each context, dominant mean ploidy and spectral gap were evaluated from 0% to 5% $O_2$ in increments of 0.025 percentage points, giving 201 oxygen values per solution and 100,500 seed–oxygen evaluations per context. The same context-appropriate operator implementation and response classifier were used for both sets. Each curve was smoothed with degree-2 Gaussian-family LOESS with span 0.20 under the versioned rule `loess_persistent_v1`; a smoothing spline with `spar`=0.65 was used when required, and raw finite values were retained if smoothing remained invalid. Persistent slope segments had to satisfy prespecified duration and amplitude thresholds; curves were then classified as approximately flat, monotone, plateauing, U-shaped, inverted U-shaped, or complex nonmonotone (Supplementary Figure 5A–B).

For seed-level reliability summaries, a response curve was flagged as unreliable when at least 25% of its oxygen-grid values had a spectral gap below 0.005. Curves were flagged for caution when any grid value had a gap below 0.005 or when at least 10% of grid values had a gap below 0.01; all remaining curves were labeled reliable. These diagnostics indicate weak separation between the first two eigendirections and are not additional fitted-data hypothesis tests.

For the family-specific steady-state response surfaces in Figure 6A, every q10 endpoint supplied context-specific *in vivo* and *in vitro* parameter vectors. Each vector was evaluated under its own model context on 201 oxygen values from 0% to 5% and 60 log-spaced fixed mechanistic $p_{\text{misseg}}$ values from 0.005 to 0.5. At each grid point, $p_{\text{misseg}}$ was set to the requested value while $p_{\text{mis,base}}$ and all other parameters retained that endpoint's fitted context-specific values. Thus, the state-specific effective probability $p_{\text{mis}}(N, O_2)$ remained model-derived rather than being forced to the input value (Supplementary Table 9).

At each oxygen–$p_{\text{misseg}}$ grid point and within each context, the displayed surface is the seed-weighted median dominant mean ploidy across the 50 eligible endpoints. Weak-gap boundaries identify cells where at least half of the endpoint vectors had dominant spectral gap below 0.005. The fitted mechanistic $p_{\text{misseg}}$ mean and the corresponding population-average state-specific missegregation probability were calculated as separate overlays, preserving the distinction between fitted input parameters and derived effective rates.

Figure 6B–C used finite-time simulations rather than steady-state eigendecomposition. For each context, family, 2N/4N initial state, and fixed mechanistic $p_{\text{misseg}}$ value of 0.005, 0.01, 0.10, 0.20, or 0.30, mean ploidy (panel B) was followed over 0–1,000 days across fixed oxygen. *In vivo* simulations used continuous natural growth. *In vitro* simulations used daily threshold-triggered stochastic passage with fixed-inoculum sampling, averaged over repeats and then over optimizer endpoints. The paired population-weighted net-live growth rate in panel C (day$^{-1}$) was mapped over the same finite-time grids and summarized with the same optimizer-endpoint and stochastic-repeat averaging; passage-day sampling and dilution were excluded from this rate. Continuous and passage-aware 0–10,000-day extensions were retained as supplementary numerical comparators and sensitivity analyses rather than being substituted for the main finite-time panels (Supplementary Figures 9 and 10–13; Supplementary Table 9).

**Embedding of oxygen-response classes and fit objectives.** For the response-class diagnostic, the 500 final endpoints from each context were taken from the same pooled 14-parameter joint-warm-start t-SNE embedding described immediately above; no second t-SNE embedding was estimated. Supplementary Figure 5C therefore uses two-dimensional pooled-embedding coordinates and does not use the *in vivo*-only 18-parameter Figure 4C coordinates. The coordinate labels denote only the first and second axes returned by t-SNE: they are unitless, may change with embedding settings, and have no direct mechanistic or biological interpretation. Endpoints were joined by context and optimizer seed to their smoothed fixed-$O_2$ response classes and complete fitted MAP objective values. Circles and triangles distinguish *in vivo* and *in vitro*; within every nonempty context–class combination, a black outline marks its lowest-objective endpoint. Supplementary Figure 5D subtracts the minimum objective within each context before comparing class distributions, so its vertical scale is relative fit quality and does not equate the two context-specific objectives. The class and objective overlays were used descriptively to assess landscape overlap, not as independent tests of class separation (Supplementary Table 4).

**Visualization and reporting.** For visualization and reporting in cell-number units, observed tumor volume (Eq. (26)) is converted to an approximate cell-count range using the specified diploid packing-density bounds:

$$N_{ij}^{\mathrm{obs,low}} = V_{ij}^{\mathrm{obs}}\,\rho_{2N,\min}, \qquad N_{ij}^{\mathrm{obs,mid}} = V_{ij}^{\mathrm{obs}}\,\sqrt{\rho_{2N,\min}\rho_{2N,\max}}, \qquad N_{ij}^{\mathrm{obs,high}} = V_{ij}^{\mathrm{obs}}\,\rho_{2N,\max}. \quad (65)$$

This conversion is used for plotting/interpretation only and is not part of the fitted likelihood, which is evaluated directly in the volume domain via Eq. (29).

#### 1.4 Parameter Tables

The active fitted parameters and their calibration bounds are summarized in Table 11; fixed numerical quantities are summarized in Table 12. Values and bounds are reported on the natural scale. The bounds define the calibration domain, while likelihood-based confidence intervals were not estimated. Some ranges were informed by published biological scales, whereas parameters without reliable direct measurements were assigned engineering bounds for numerical calibration.

For the reported joint fits, the *in vivo*/*in vitro* bounds for each overlapping active biological parameter are merged into the joint union interval in Eq. (49), and all 14 such parameters are represented by the center–delta construction in Eq. (50). Parameters used only by one context retain their component-specific bounds.

Supplementary Table 11: Active fitted parameters and natural-scale bounds used by the separate and joint analyses reported here. A dash means that the parameter is not active in that component. For the 14 overlapping biological parameters, the joint admissible interval is the union of the separate *in vivo* and *in vitro* intervals. The entries were source-authenticated against the current Figure 3 separate-*in vitro* ranges, Figure 4 separate-*in vivo* configuration, and Figure 5 joint-fit configurations. These bounds are calibration constraints, not confidence intervals.

| Parameter | *In vivo* bounds | *In vitro* bounds | Joint bounds | Role in the reported model |
|---|---|---|---|---|
| $\lambda_{\max}$ | 0.005–0.300 | 0.005–1.500 | 0.005–1.500 | Maximum division rate (day$^{-1}$); soft-coupled. |
| $p_{\mathrm{mis,base}}$ | $10^{-6}$–0.100 | $10^{-6}$–0.005 | $10^{-6}$–0.100 | Baseline per-chromosome missegregation probability; soft-coupled. |
| $p_{\mathrm{misseg}}$ | 0.005–0.500 | 0.005–0.400 | 0.005–0.500 | Maximal stress-induced missegregation increment; soft-coupled. |
| $k_{o,\mathrm{mis}}$ | 0.001–0.020 | $10^{-5}$–0.020 | $10^{-5}$–0.020 | Death-hazard half-saturation scale for stress-induced missegregation (day$^{-1}$); soft-coupled. |
| $s_{\max}$ | 0–1 | 0–1 | 0–1 | Maximum per-copy post-missegregation survival factor; soft-coupled. |
| $\beta_{\mathrm{buf}}$ | 0.01–10 | 0.01–10 | 0.01–10 | Ploidy-dependent post-missegregation viability-loss strength; soft-coupled. |
| $n_{\exp}$ | 0.1–10 | 0.1–10 | 0.1–10 | Exponent controlling ploidy dependence of post-missegregation survival; soft-coupled. |
| $p_{\mathrm{wgd}}$ | $10^{-5}$–$10^{-3}$ | $10^{-5}$–$10^{-3}$ | $10^{-5}$–$10^{-3}$ | Constant per-division WGD probability; soft-coupled. |
| $\alpha_{O_2}$ | 0.5–4.0 | 0.5–4.0 | 0.5–4.0 | Strength of resource-stress-dependent growth damping; soft-coupled. |
| $\gamma_{\mathrm{growth}}$ | 1.5–4.5 | 1.5–4.5 | 1.5–4.5 | Chromosome-number dependence of growth damping; soft-coupled. |
| $\mu_{\mathrm{hp}}$ | 0.005–0.300 | 0.005–0.300 | 0.005–0.300 | Stress-associated death scale (day$^{-1}$); soft-coupled. |

| Parameter | *In vivo* bounds | *In vitro* bounds | Joint bounds | Role in the reported model |
|---|---|---|---|---|
| $\gamma_\mu$ | 1.2–3.5 | 1.2–3.5 | 1.2–3.5 | Chromosome-number penalty on stress-associated death; soft-coupled. |
| $O_{2,c}$ | 0.1–2.5 | $10^{-4}$–2.5 | $10^{-4}$–2.5 | Critical level for the oxygen-linked resource-stress function (% $O_2$); soft-coupled. |
| $n_O$ | 0.5–5.0 | $10^{-4}$–5.0 | $10^{-4}$–5.0 | Hill exponent for the oxygen-linked resource-stress function; soft-coupled. |
| $S_0$ | 2.0–5.0 | – | *In vivo* only | Low-burden effective oxygen supply level (% $O_2$). |
| $\kappa_O$ | 0.1–1.2 | – | *In vivo* only | Oxygen-drop amplitude in the supply–demand target. |
| $\eta$ | 1.0–2.0 | – | *In vivo* only | Chromosome-number-weighted oxygen-demand exponent. |
| $\rho_{2N}$ | $5.0\times10^4$–$5.0\times10^5$ | – | *In vivo* only | Diploid-cell density (cells $mm^{-3}$). |
| $k_{\text{clear}}$ | 0.005–0.300 | – | *In vivo* only | Dead-biomass clearance rate ($day^{-1}$). |
| $\sigma_{\text{burden}}$ | 0.100–0.150 | – | *In vivo* only | Log-scale burden observation standard deviation. |
| $\sigma_{\text{growth}}$ | – | 0.001–1.000 | *In vitro* only | Growth-rate observation standard deviation. |
| $\sigma_{\text{kary}}$ | – | 0.01–10.0 | *In vitro* only | Chromosome-number observation smoothing/error scale. |
| $\mu_{2N,0}$ | – | 45.725–49.725 | *In vitro* only | Initial near-diploid Gaussian chromosome-count mean. |
| $\sigma_{2N,0}$ | – | 0.5–3.0 | *In vitro* only | Initial near-diploid Gaussian chromosome-count SD. |
| $\mu_{4N,0}$ | – | 83.667–91.667 | *In vitro* only | Initial near-tetraploid Gaussian chromosome-count mean. |
| $\sigma_{4N,0}$ | – | 1.5–6.0 | *In vitro* only | Initial near-tetraploid Gaussian chromosome-count SD. |

Supplementary Table 12: Fixed numerical settings used by the reported resource-stress supply–demand model and calibration workflow. Values are reported on the natural scale; settings for inactive optional modules are identified explicitly. The entries were source-authenticated against the current Figure 4 fit configuration: the 5% oxygen cap is inherited from the fitted-supply upper bound, and the optional logistic crowding term was disabled in the reported fits.

| Parameter | Value | Unit | Description | Reference |
|---|---|---|---|---|
| $\tau_{O_2}$ | 0.1 | day | Oxygen relaxation timescale. | Fixed numerical relaxation setting; empirical workflow choice rather than a literature-derived parameter. |
| $O_{2,\mathrm{cap}}$ | 5.0 | % $O_2$ | Maximum oxygen level used to cap the supply-demand oxygen trajectory, inherited from the $S_0$ upper bound. | Bertout et al. (2008)[39]; McKeown (2014)[40]. |
| $O_{2,\mathrm{min}}$ | 0 | % $O_2$ | Lower oxygen floor used in the logarithmic oxygen target relation. | Physical/numerical anoxia floor used to bound oxygen values; model constraint rather than literature-estimated threshold. |
| $N_{\mathrm{ref}}$ | $10^6$ | cells | Reference viable burden scale used in the oxygen supply-demand relation. | Oxygen-feedback normalization constant; empirical model scaling value rather than a measured biological quantity. |
| $N_{\mathrm{unit}}$ | 22 | autosomal chromosomes | Modeled autosomal chromosome count corresponding to ploidy 1. | Model convention defining one haploid autosome set as ploidy 1; definition rather than literature-estimated parameter. |
| $N_{\mathrm{min}}$ | 22 | autosomal chromosomes | Minimum autosomal chromosome-count state retained in the discrete state space. | Discrete state-space lower bound anchored to the model ploidy convention; grid-design choice. |
| $N_{\mathrm{max}}$ | 154 | autosomal chromosomes | Maximum autosomal chromosome-count state retained in the discrete state space. | Bielski et al. (2018)[1]; Zack et al. (2013)[41]. |
| $\Delta t$ | 0.05 | day | Simulation time step. | Simulation time step selected for numerical resolution and stability; computational setting rather than biological parameter. |
| $K$ | $10^{12}$ | cells | Carrying-capacity setting for the optional logistic crowding term; crowding was disabled in the reported fits. | Waclaw et al. (2015)[42]. |
| $N(0)$ | $10^6$ | cells | Initial total tumor population size. | Initial-condition scale for the simulation/calibration workflow; empirical setup choice. |
| $\sigma_{\mathrm{ploidy}}$ | 0.12 | chromosomes | Fixed endpoint autosomal chromosome-number observation-noise scale in the harvest likelihood. | Fixed likelihood observation-noise term; empirical measurement-error setting in the calibration workflow. |
| $\lambda_{\mathrm{nec}}$ | 1 | unitless | Relative weight of the terminal necrosis loss in the *in vivo* objective (`lambda_necrosis` in the run configuration). | Reported calibration setting. |
| $\sigma_{\mathrm{nec}}$ | 0.75 | logit-fraction units | Scale used to standardize terminal necrosis discrepancies (`sigma_necrosis_logit` in the run configuration). | Reported calibration setting. |
| $\epsilon_{\mathrm{nec}}$ | $10^{-4}$ | fraction | Clipping constant for observed and predicted necrosis fractions before the logit transform (`necrosis_fraction_eps` in the run configuration). | Numerical safeguard in the reported calibration. |

| Parameter | Value | Unit | Description | Reference |
|---|---|---|---|---|
| $\lambda_{\mathrm{prior}}$ | 0.03 | unitless | Relative weight of the soft-prior contribution to the *in vivo* objective. | Reported calibration setting. |
| $\varepsilon_{\log B}$ | $10^{-12}$ | $\mathrm{mm}^3$ | Volume-domain floor used inside $\log\{\max(V, \varepsilon_{\log B})\}$ for burden comparisons. | Numerical floor preventing log(0); computational safeguard. |
| $N_{\mathrm{min,pop}}$ | $10^{-12}$ | cells | Minimum population floor used for numerical stability. | Population floor used to avoid numerical underflow or zero states; computational safeguard. |

Supplementary Table 13: Function-level contrasts reconstructed from the 2 selected joint-fit winners displayed in Figure 5. Ranges span C01–C02. Survival columns report the *in vivo* and shared *in vitro* values; missegregation and proliferation rows report the *in vivo*/*in vitro* ratio. The optimizer-selected fits are neither biological replicates nor confidence-interval samples.

| Derived quantity | Reference state | *In vivo* value or ratio range | *In vitro* reference value | Direction across winners |
|---|---|---|---|---|
| Per-copy post-missegregation survival | $N = 44$ | 0.660–0.771 | 0.202 | higher *in vivo*, 2/2 |
| Per-copy post-missegregation survival | $N = 88$ | 0.899–0.909 | 0.836 | higher *in vivo*, 2/2 |
| Survival gradient, $s_{88} - s_{44}$ | $44 \rightarrow 88$ | 0.138–0.239 | 0.634 | larger *in vitro*, 2/2 |
| Proliferation-rate ratio | 0% $O_2$, $N = 44$ | 0.07–0.10 | not applicable | $< 1$, 2/2 |
| Proliferation-rate ratio | 0% $O_2$, $N = 88$ | 0.02–0.03 | not applicable | $< 1$, 2/2 |
| Proliferation-rate ratio | 1% $O_2$, $N = 44$ | 0.22–0.23 | not applicable | $< 1$, 2/2 |
| Proliferation-rate ratio | 1% $O_2$, $N = 88$ | 0.18–0.24 | not applicable | $< 1$, 2/2 |
| Proliferation-rate ratio | 5% $O_2$, $N = 44$ | 0.19–0.19 | not applicable | $< 1$, 2/2 |
| Proliferation-rate ratio | 5% $O_2$, $N = 88$ | 0.20–0.20 | not applicable | $< 1$, 2/2 |
| Effective missegregation-probability ratio | 0% $O_2$, $N = 44$ | 11.22–14.94 | not applicable | $> 1$, 2/2 |
| Effective missegregation-probability ratio | 0% $O_2$, $N = 88$ | 11.29–15.66 | not applicable | $> 1$, 2/2 |
| Effective missegregation-probability ratio | 1% $O_2$, $N = 44$ | 5.96–7.48 | not applicable | $> 1$, 2/2 |
| Effective missegregation-probability ratio | 1% $O_2$, $N = 88$ | 10.46–12.49 | not applicable | $> 1$, 2/2 |
| Effective missegregation-probability ratio | 5% $O_2$, $N = 44$ | 0.74–0.82 | not applicable | $> 1$, 0/2 |
| Effective missegregation-probability ratio | 5% $O_2$, $N = 88$ | 3.24–3.65 | not applicable | $> 1$, 2/2 |
